# Optimising Chemotherapy for Advanced High-Grade Serous Ovarian Cancer via Delay-Differential Equations

**DOI:** 10.1101/2025.06.05.658173

**Authors:** Cristina Koprinski, Georgio Hawi, Peter S. Kim

## Abstract

Ovarian cancer is the deadliest gynaecological cancer and the fourth leading cause of cancer deaths in women. High-grade serous ovarian cancer (HGSOC) accounts for 75% of cases, and chemotherapy resistance and relapse occur in 85% of patients, leading to a 5-year survival of 45%. Currently, the literature lacks comprehensive immunobiological models of HGSOC, and developing such models could provide critical insights into the disease’s underlying mechanisms and interactions within the tumour microenvironment. We address this by constructing an immunobiological model using delay differential equations and then optimise chemotherapy regimens to maximise efficacy, minimise toxicity, and improve treatment efficiency for first-line treatment. The model consists of two compartments, the tumour site and tumour-draining lymph node, with immune processes such as DC maturation, T cell priming and proliferation, and cytokine interactions modelled. Parameter values are estimated using experimental data from ovarian cancer tissue samples as well as the TCGA OV database. The results indicate that low-dose dose more frequent chemotherapy provides comparable results to the standard regimen with a lower toxicity, and alternative dosing strategies with rest weeks can allow patients to recover from the toxic side effects of chemotherapy.

## 1 Introduction

High-grade serous ovarian cancer (HGSOC) is the most common subtype of ovarian cancer and the fourth leading cause of cancer-related deaths in women worldwide, with a 5-year survival rate of 45% [1]. Its poor prognosis is partly due to asymptomatic early stages and a lack of screening programs, resulting in approximately 80% of cases being diagnosed at Stages III or IV [2]. At the same time, 70–80% of HGSOC patients experience recurrence after primary surgery and resistance to initial chemotherapy, contributing to poor long-term survival outcomes [3–5]. Of new HGSOC cases, half of the diagnoses occur in women over 64, and the disease burden is expected to increase due to ageing populations and increased life expectancy [6].

The primary first-line treatment for patients with HGSOC is neoadjuvant chemotherapy (NACT) followed by surgery or adjuvant chemotherapy (ACT) after surgery aimed at removing all visible tumours within the peritoneal cavity, with the degree of cytoreduction being a critical prognostic factor [7, 8]. Randomised clinical trials, including EORTC 55971, CHORUS, JCOG0602, and SCORPION-NCT01461850, have shown no significant difference in overall survival and progression-free survival (PFS) between NACT and ACT in ovarian cancer [9–12]. However, in advanced cancer, NACT is preferred due to its potential to facilitate increased rates of optimal cytoreduction [13, 14]. The standard chemotherapy regimen (SR) has been the combination of paclitaxel (175 mg/m^2^; 3h infusion) and carboplatin (AUC 5 or 6; 30 min infusion), administered intravenously every three weeks for three/- four cycles in the NACT and five/six cycles in ACT [9, 15–17]. Trials in the mid-1990s established that this regimen is superior to cisplatin and cyclophosphamide [18, 19], with later studies confirming that substituting cisplatin for carboplatin offers similar efficacy with improved safety [20, 21]. The SR dosing is based on the principle that higher doses increase the effectiveness of cancer cell eradication while maintaining tolerability [22, 23]. While aiming to maximise tumour depletion, chemotherapy’s cytotoxic effects on healthy tissues and high toxicity, including neurotoxicity, require prolonged rest periods and impair patients’ quality of life [24].

In the past decade, there has been growing interest in alternative dosing, including more frequent, low-dose drug administrations (dose-dense and metronomic therapies), due to their potential efficacy and low toxicity in various cancers [25]. In the Japanese JGOG3016 trial, patients with Stage II, III and IV epithelial ovarian cancer were randomised to receive triweekly carboplatin (AUC 6) and weekly paclitaxel (80 mg/m2), or the SR, with results showing significantly improved PFS compared to the SR [26]. However, the GOG-0262 trial, which trialled the same regime in a European population, did not find a significantly improved PFS to the SR [27]. Separately, a metronomic dosing regimen of weekly carboplatin (AUC 2) and paclitaxel (60 mg/m2) was trialled in elderly patients in the MITO-5 trial, concluding comparable results to the SR with reduced toxicity, and was further supported by the MITO-7 trial in a larger cohort [28, 29]. This prompted the ICON8 study, which found comparable results to the SR for both these dosing regimens; however, as no significant improvement in PFS was observed, the SR remains the standard of care in European populations [30]. Despite this, dose-dense and metronomic regimens remain useful for patients who cannot tolerate the SR, such as elderly and vulnerable patients [31].

An important question is the appropriate dosing and spacing of paclitaxel and carboplatin therapies to balance tumour reduction with factors such as toxicity and efficacy. This can be achieved through the construction of immunobiological mathematical models and has been employed in cancers such as colorectal, breast and pancreatic to optimise chemotherapy and other treatment regimens [32–34]. These models also provide insights into the underlying biological processes guiding further research and clinical trials [35]. Our work builds on [36] and [37] by applying a two-compartment model, incorporating the tumour site (TS) and the tumour-draining lymph node (TDLN), to HGSOC and chemotherapy. This enables more accurate modelling of immune cell types and interactions, such as antigen presentation and cell proliferation in the TDLN. To the authors’ knowledge, no existing mathematical models in the literature consider the immunobiology of HGSOC with paclitaxel and carboplatin. Few related models exist, the first by Panetta who used a two-compartment model of proliferating and quiescent cancer cells to investigate the effect of varying dosages and infusion times of paclitaxel, and the second by Kohandel et al. who expanded this model to investigate the effectiveness of NACT and ACT therapies [38, 39]. Although these models provide a modelling foundation, they do not incorporate the role of various immune cells and cytokines in the TME, opting to only consider the cancer cell concentration over time.

Hence, it is important to briefly outline the functions of immune cells and cytokines in the tumour microenvironment (TME), as their interaction with cancer cells directly influences the efficacy of chemotherapy treatments. T cell activation takes place in the lymph node through T-cell receptor (TCR) recognition of MHC class I molecules, which activate CD8+ T cells, and MHC class II molecules, which activate CD4+ T cells, expressed on mature dendritic cells. CD8+ cells and NK cells are some of the most cytotoxic cells involved in cancer cell lysis, and they secrete pro-inflammatory cytokines such as IL-2, IFN-*γ*, and TNF. These are also secreted by Th1 cells, which differentiate from naïve CD4+ T cells, and are important in the activation of macrophages, which can differentiate into two types: classically activated M1 macrophages and alternatively activated M2 macrophages. M1 macrophages contribute to the inflammatory response by releasing cytokines such as TNF and IL-12, while M2 macrophages contribute to wound repair and the anti-inflammatory response by releasing cytokines such as IL-10 and TGF-*β* and promote Th2 responses. Of note is that this M1/M2 dichotomy is a simplification, as macrophages exhibit high plasticity depending on their environmental signals [40]. However, we consider macrophage polarisation and repolarisation between these types to account for this simplification. We also consider Tregs, which mediate inflammatory processes such as reducing the synthesis of pro-inflammatory cytokines by producing immunosuppressive cytokines such as TGF-*β* and IL-10.

In this paper, we construct a mechanistic mathematical model of HGSOC using delay differential equations, incorporating immune cells, including CD4+ and CD8+ T cells, dendritic cells, macrophages, NK cells, cytokines, and damage-associated molecular patterns. We use experimental data to estimate the model’s initial conditions and parameter values, incorporating deconvolution of bulk RNA-sequencing data from the TCGA OV database. Finally, we optimise the first-line neoadjuvant treatment of advanced HGSOC with carboplatin and paclitaxel, focusing on maximising efficacy, minimising toxicity, and improving treatment efficiency.

## 2 Mathematical Model of Ovarian Cancer

### 2.1 Modelling Assumptions

We assume cancer cell necrosis by tumour necrosis factor (TNF), interferon gamma (IFN-*γ*), and from the first-line chemotherapy treatment with paclitaxel. The necrosis of cancer cells releases danger-associated molecular patterns (DAMPs). In our model, we consider the DAMPs High Mobility Group Box 1 (HMGB1) and surface calreticulin, which bind to their respective receptors on immature dendritic cells (DCs), promoting their maturation and the presentation of tumour antigens [41]. Mature DCs migrate from the TS to the tumour-draining lymph node (TDLN) through chemokine circuits, where they prime naïve CD8+ T cells, CD4+ T cells and regulatory T cells (Tregs) [42, 43]. Upon DC activation, naïve CD8+ T cells undergo clonal expansion into mature cytotoxic CD8+ T cells in the TDLN [44]. We assume that naïve CD4+ T cells differentiate into either Th1 or Th2 T cells and then undergo clonal expansion in the TDLN. In the TLDN, naïve Tregs are activated by mature DCs and then undergo clonal expansion and proliferation. Chemotherapy inhibits the proliferation of all T cells in the TDLN, and after clonal expansion, these cells migrate to the TS. A diagram of these processes is provided in Figure 1.

**Figure 1:**
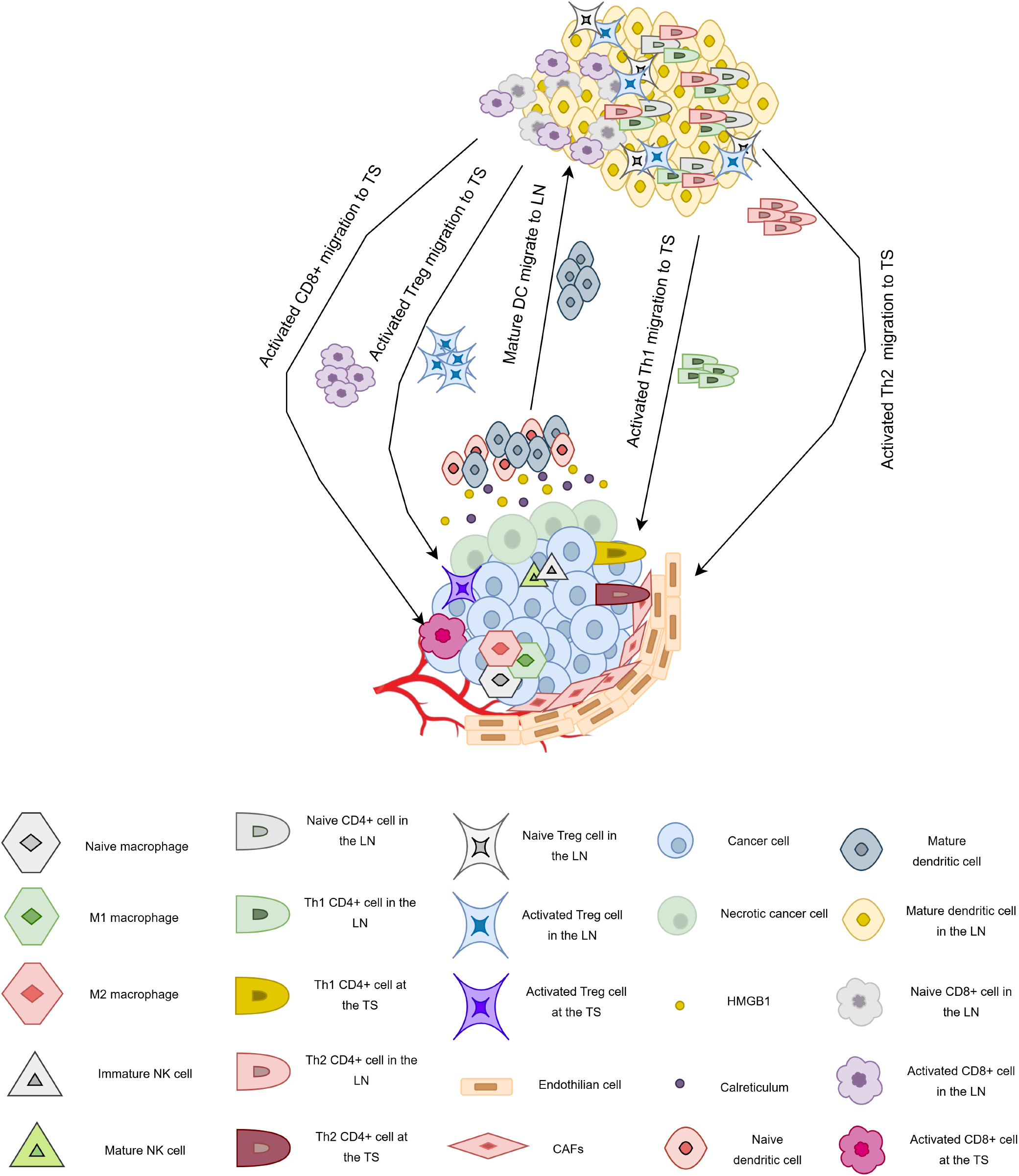
The relationship between the TS (bottom) and TLDN (top). DAMPs released by necrotic cancer cells activate a proportion of native DCs to mature and travel to the LN, with the cancer antigen, where they activate naïve CD8+, CD4+ T and Tregs. In the TDLN, these cells differentiate into effector CD8+ T cells, Th1, Th2 and Tregs, and upon maturation, migrate to the TS.

We also examine seven key cytokines involved in the pathogenesis of ovarian cancer, as shown in Figure 2. The pro-inflammatory cytokines include interleukin-2 (IL-2), interleukin-12 (IL-12), interferon-*γ* (IFN-*γ*), and tumour necrosis factor (TNF). We assume IL-2 is primarily released by CD8+ T cells and Th1 T cells. It plays a crucial role as a growth factor for the proliferation of CD8+ T cells, Th1 T cells, and NK cells [45, 46]. IL-12, produced by dendritic cells and M1 macrophages, is key to the activation and proliferation of NK cells [47]. IFN-*γ* and TNF, produced by Th1 cells, NK cells, and CD8+ T cells, promote the polarisation of macrophages to the M1 pro-inflammatory phenotype and induce the necrosis of cancer cells [48, 49].

**Figure 2:**
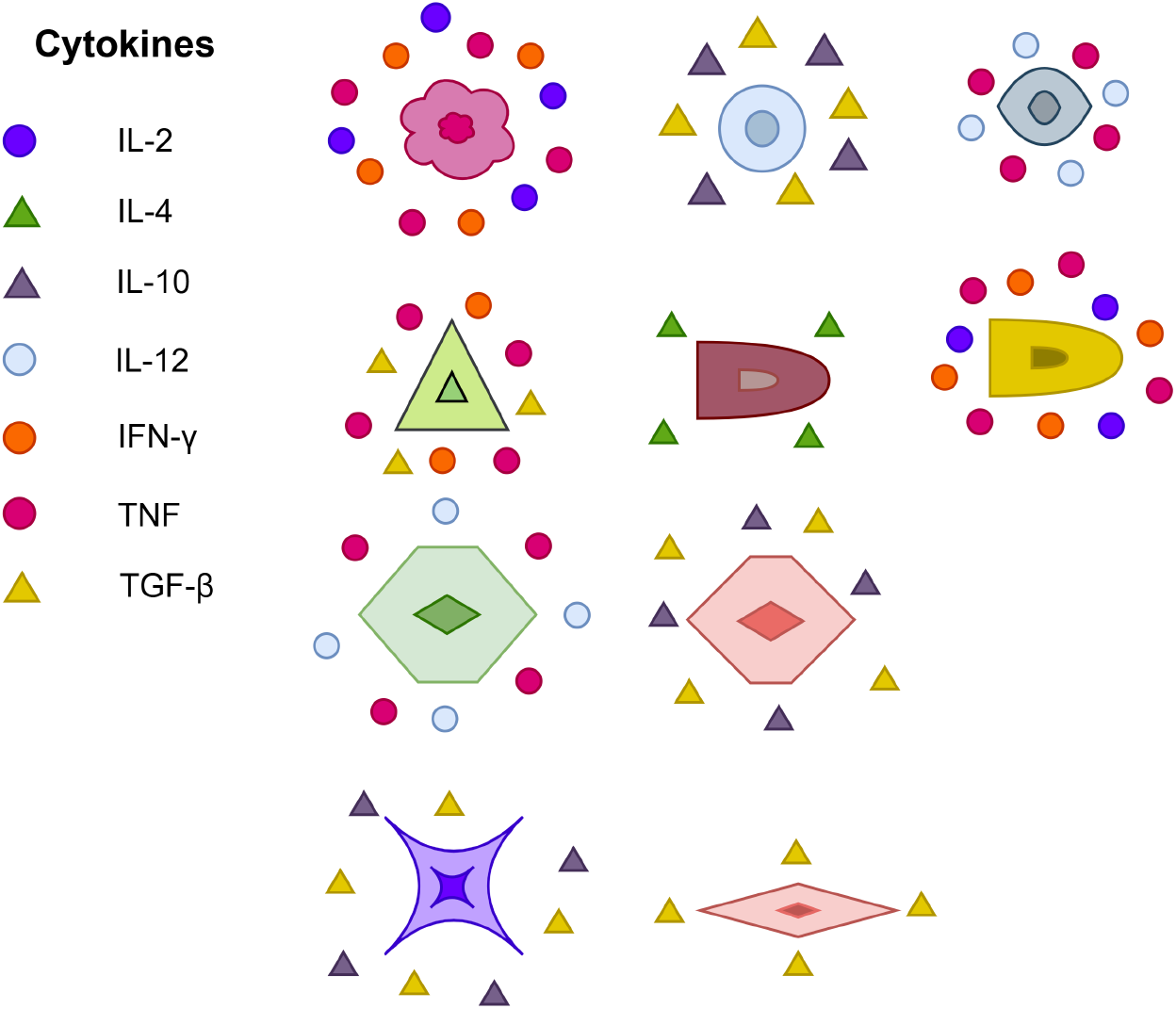
Immune cells and the cytokines they release. Pro-inflammatory cytokines are displayed by a circle, and anti-inflammatory cytokines are displayed by a triangle.

We also consider interleukin-4 (IL-4), interleukin-10 (IL-10), and transforming growth factor-*β* (TGF-*β*) as key anti-inflammatory cytokines. IL-4 and IL-10 promote the polarisation of naïve and M1 macrophages into the M2 type. We also assume, IL-4 is produced by Th2 cells, while IL-10 is produced by Tregs, M2 macrophages, and cancer cells. TGF-*β*, primarily involved in inhibiting CD8+ T cells and NK cells in cancer lysis, is produced by Tregs, cancer cells, M2 macrophages, and NK cells.

Additionally, we note that chemotherapy, including carboplatin and paclitaxel, works by killing rapidly proliferating cells, including cancer cells and T cells. In particular, carboplatin binds to DNA, interfering with DNA synthesis in the S phase of the cell cycle, inhibiting replication and leading to apoptosis [50, 51]. However, paclitaxel induces mitotic arrest at the G2/M phases of the cell cycle, disturbing cell division, and resulting in cell death [52, 53]. We model this by considering carboplatin and paclitaxel apoptosis and necrosis of cancer cells within the growth term, and by inhibiting CD8+ and CD4+ cell proliferation in the TDLN after activation.

We also assume the following biological and modelling conditions. Immature dendritic cells are cancer antigen-bearing and are activated by a proportion of DAMPs released by necrotic cells. This stimulates immature dendritic cells to mature and a proportion of these travel to the TDLN. When mature DCs in the TDLN activate naïve CD8+ and CD4+ T cells, the T cells differentiate and activate instantaneously, with their naïve cell concentrations remaining unchanged during proliferation. Effector and activated cells produce cytokines, and macrophages follow an M1/M2 dichotomy. A constant solution history is used, with each species’ history set to its respective initial condition, and each component *X* degrades at a rate *d*_*X*_, proportional to its concentration. All variables and their units are shown in Table 1.

**Table 1:** Variables used in the model, with variables in the top boxes having units of cell*/*cm^3^, and all other variables having units of g*/*cm^3^. All quantities refer to tumour site density/concentration unless otherwise specified. TDLN denotes the tumour-draining lymph node, whilst TS denotes the tumour site.

| Var | Description | Var | Description |
| --- | --- | --- | --- |
| $C$ | Viable cancer cell density | $N$ | Necrotic cancer cell density |
| $H$ | HMGB1 | $R$ | Calreticulin |
| $D_0$ | Naïve dendritic cell density | $D$ | Mature dendritic cell density |
| $D_L$ | Mature dendritic cell density in the TDLN | $T_0^8$ | Naïve CD8+ T cell density in the TDLN |
| $T_8$ | Effector CD8+ T cell density in the TDLN | $T_8$ | CD8+ T cell density in the TS |
| $T_0^4$ | Naïve CD4+ T cell density in the TDLN | $L_1$ | Effector Th1 cell density in the TDLN |
| $T_1$ | Effector Th1 cell density in the TS | $L_2$ | Effector Th2 cell density in the TDLN |
| $T_2$ | Effector Th2 cell density in the TS | $L_0^r$ | Naïve Treg cell density in the TDLN |
| $L_r$ | Effector Treg cell density in the TDLN | $T_r$ | Effector Treg cell density in the TS |
| $M_0$ | Naïve macrophages | $M_1$ | M1 macrophage density |
| $M_2$ | M2 macrophage density | $K_0$ | Naïve NK cell density |
| $K$ | Activated NK cell density | $I_2$ | IL-2 concentration |
| $I_4$ | IL-4 concentration | $I_{12}$ | IL-12 concentration |
| $I_{10}$ | IL-10 concentration | $I_\gamma$ | IFN-γ concentration |
| $I_\alpha$ | TNF concentration | $I_\beta$ | TGF-β concentration |
| $P$ | Paclitaxel concentration in the TS | $A$ | Carboplatin concentration in the TS |
| $P^{\text{LN}}$ | Paclitaxel concentration in the TDLN | $A^{\text{LN}}$ | Carboplatin concentration in the TDLN |

Finally, we assume that the rate of activation and polarisation of *X* by *Y* follows the Michaelis-Menten kinetic law 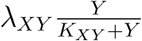, where *λ*_*XY*_ is the rate constant and *K*_*XY*_ is the half-saturation constant. We model the rate of inhibition of *X* by *Y* using 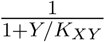, where *λ*_*XY*_ is the rate constant and *K*_*XY*_ is the half-saturation constant. We model the production of *X* by *Y* using mass-action kinetics using *λ*_*XY*_ *XY* for some positive constant *λ*_*XY*_. We also assume that lysis of *X* by *Y* follows mass-action when *Y* is a cell, and Michaelis-Menten when *Y* is a cytokine.

### 2.2 Model Equations

The model equations and structure follow similarly to many of the model equations in [36] and [37].

#### 2.2.1 Equation for Cancer Cells (*C*)

The first term models cancer growth using a logistic function, where *r*_*C*_ is the baseline growth rate and *C*_0_ is the carrying capacity. It also accounts for apoptosis induced by paclitaxel and carboplatin, with paclitaxel additionally inducing immunogenic cell death. The second term accounts for CD8+ T cell lysis of the cancer cells, with this process inhibited by TGF-*β* [54, 55]. The third and fourth terms model CD4+ Th1 and NK cell recognition and lysis of cancer cells, respectively [56–58]. Tregs also inhibit the cytotoxicity of CD8+ T cells, CD4+ T cells and NK cells, and are accounted for in the second, third and fourth terms [59–61]. TGF-*β* also inhibits the cytotoxicity of NK cells, as shown in the fourth term [62]. Lastly, the fifth and sixth terms model cancer cell necroptosis by TNF [63, 64] and IFN-*γ* [65]. This gives us

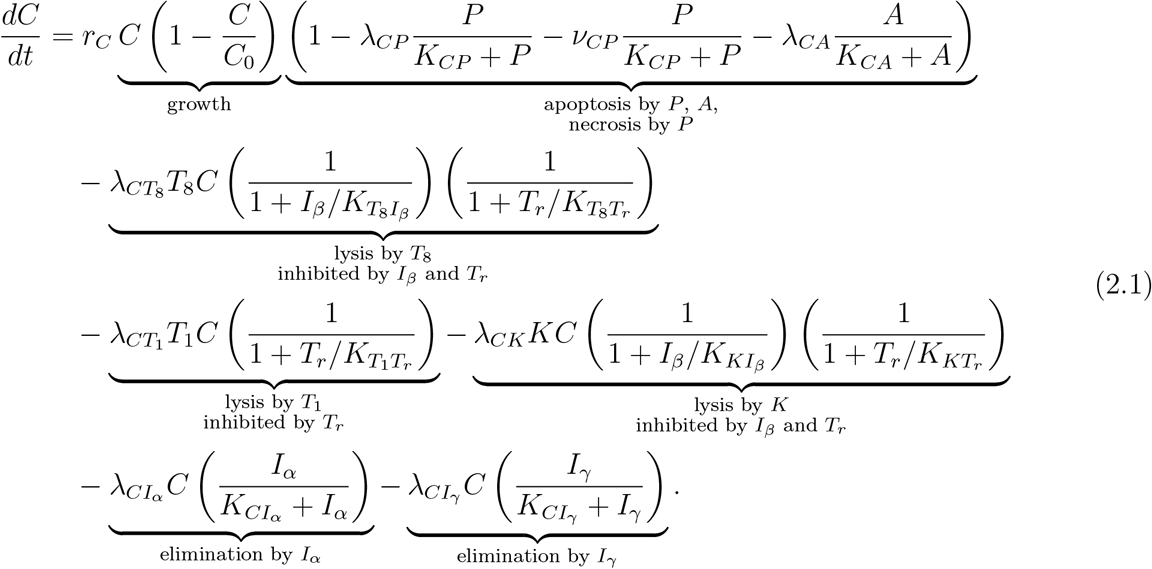

#### 2.2.2 Equation for Necrotic Cells (*N*)

The first to third terms account for cancer cell necrosis by TNF, IFN-*γ* and paclitaxel, respectively [63–65]. This gives us

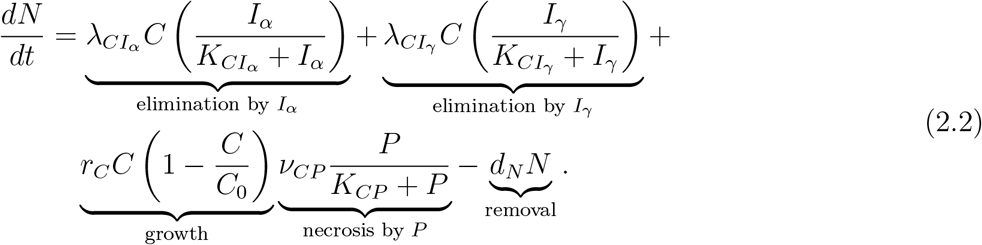

#### 2.2.3 Equation for HMGB1 (*H*)

As shown in the first term, we assume that a proportion of necrotic cells release HMGB1 [66] This gives us

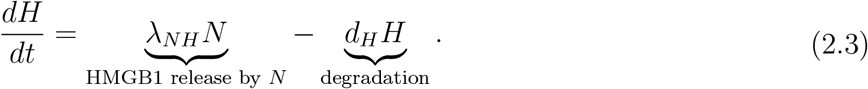

#### 2.2.4 Equation for Calreticulin (*R*)

As shown in the first term, we assume that a proportion of necrotic cells release calreticulin [66]. This gives us

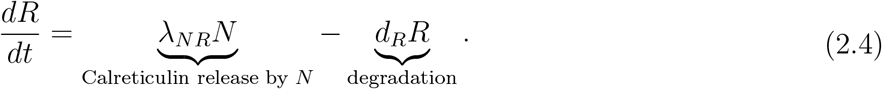

#### 2.2.5 Equation for Immature DCs in the TS (*D*_0_)

In the first term, we assume that immature dendritic cells are supplied at a constant rate 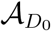. In the second term, DAMPs activate a proportion of DCs in the TS, with DC maturation in the TS inhibited by IL-10 and TGF-*β* [67, 68]. This gives us

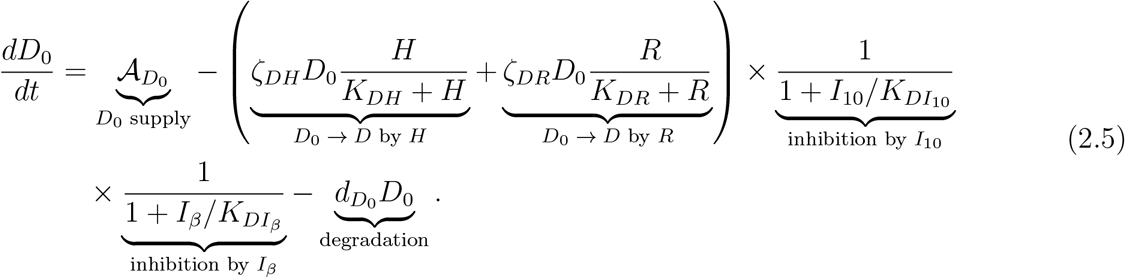

#### 2.2.6 Equation for Mature DCs in the TS (*D*)

The first term accounts for the activation of DCs from (2.5) and the second term assumes that a proportion of mature DCs migrate to the TDLN. This gives us

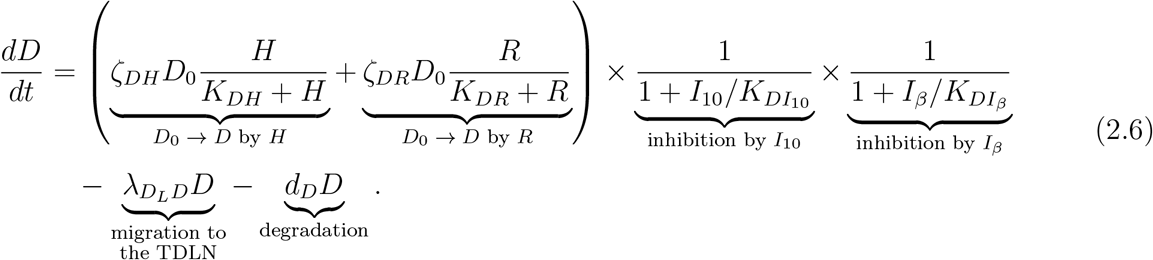

#### 2.2.7 Equation for Mature DCs in the TDLN (*D*_*L*_)

The exponential in the first term accounts for the degradation of DCs during the transit, of time *τ*_*m*_, from the TS to the TDLN. The degradation rate during transit is similar to the degradation of mature dendritic cells in the TS, and we assume that they are the same [69]. As the volume between the TS and TDLN is different, we account for the volume change to maintain the correct concentrations by multiplying by *V*_TS_*/V*_LN_, where *V*_TS_ is the volume of the TS, and *V*_LN_ is the volume of the LN. We also consider the transit time of dendritic cells arriving at the TDLN as having left the TS at time *t − τ*_*m*_. This gives us

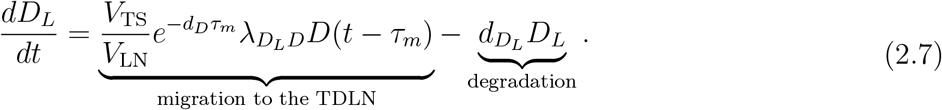

#### 2.2.8 Equation for Naïve CD8+ T Cells in the TDLN (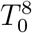)

We assume that naïve CD8+ T cells are supplied at a rate 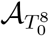 in the first term. The second term accounts for the activation of naïve CD8+ T cells by DCs in the TDLN, inhibited by Tregs [70, 71]. We assume that this activation is instant, and gives us

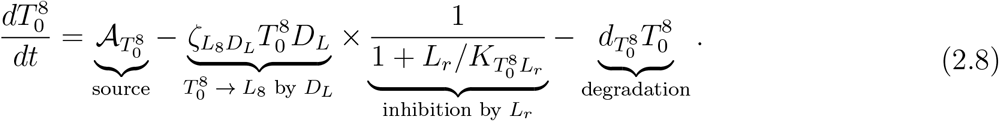

#### 2.2.9 Equation for Effector CD8+ T Cells in the TDLN (*L*_8_)

The exponential in the first term accounts for the degradation of activated CD8+ T cells during proliferation dependent on *τ*_8_, the time for *T*_8_ to complete clonal expansion, and *N*_8_, the number of cell divisions in clonal expansion before proliferation stops. We assume proliferation takes *τ*_8_ time to finish, the degradation rate during proliferation is the same for effector CD8+ T cells in the TDLN, and that the concentration of effector Tregs in the TDLN does not change significantly during proliferation. We also consider the activation of 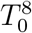 by *D*_*L*_ at time *t−τ*_8_, and the inhibition of CD8+ T cell proliferation due to Tregs, carboplatin and paclitaxel [51, 52, 72].

In particular, since the killing of proliferating CD8+ T cells in the TDLN due to carboplatin and paclitaxel occurs during the whole proliferation process, it is not sufficient to consider point estimates of carboplatin and paclitaxel concentration. Instead, we resort to considering the integrals of the concentrations of the relevant species throughout the entire *τ*_8_ time that it takes for the cell division program to be completed. This is because these integrals are proportional (with a proportionality constant of 1*/τ*_8_) to the average concentration of these species throughout activation, allowing us to properly incorporate the inhibition.

The second term accounts for the migration of effector CD8+ T cells from the TDLN to the TS. This gives us

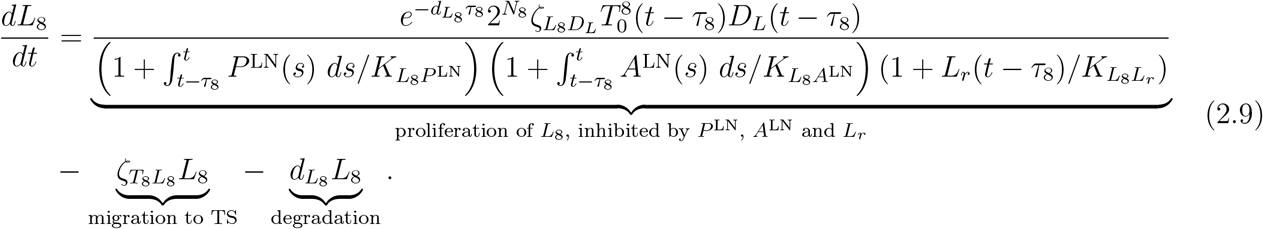

#### 2.2.10 Equation for Effector CD8+ T Cells in the TS (*T*_8_)

The exponential in the first term accounts for the degradation of effector CD8+ T cells during the transit of time *τ*_*a*_ from the TS to the TDLN. We account for the volume difference between the TDLN and TS by multiplying this term by *V*_LN_*/V*_TS_. We further assume that the degradation during transit is the same as the degradation of effector CD8+ T cells in the TS, and consider the transit time of DCs arriving in the TS as having left the TDLN at time *t−τ*_*a*_. The second term accounts for the proliferation of CD8+ T cells in the TS by IL-2, inhibited by Tregs [71, 73]. The third term accounts for the degradation of CD8+ T cells in the TS, inhibited by IL-10 [74]. Thus,

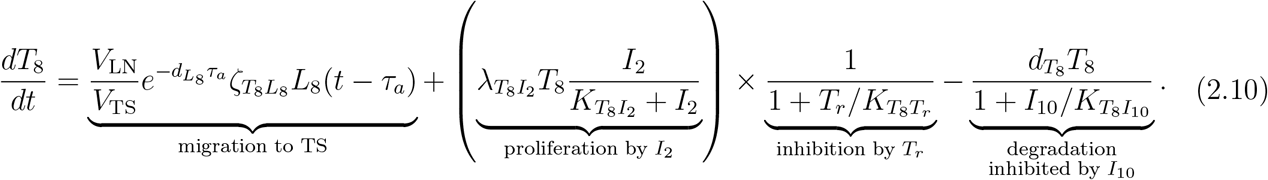

#### 2.2.11 Equation for Naïve CD4+ T Cells in the TDLN 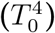

In the first term, we assume that naïve CD4+ T cells are supplied at a rate 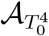. In the second term, we account for DC-induced activation of naïve CD4+ T cells which differentiate into the Th1 and Th2 phenotypes, respectively, in the TDLN inhibited by Tregs [70, 71]. This gives us that

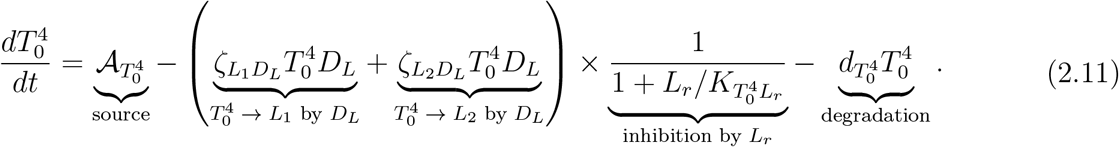

#### 2.2.12 Equation for Effector Th1 Cells in the TDLN (*L*_1_)

The exponential in the first term accounts for the degradation of activated Th1 cells during proliferation dependent on *τ*_1_, the time for *L*_1_ to complete clonal expansion, and *N*_1_, the number of cell divisions in clonal expansion before proliferation stops. We assume that proliferation takes *τ*_1_ time to finish, the degradation during proliferation is the same as the degradation of effector Th1 cells in the TDLN, and that the concentration of effector Tregs in the TDLN does not change significantly during proliferation. We also consider the activation of 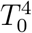to *L*_1_ by *D*_*L*_ at time *t − τ*_1_, and the inhibition of Th1 cell proliferation due to carboplatin, paclitaxel and Tregs [72]. The second term accounts for the migration of effector Th1 cells from the TDLN to the TS. This gives us

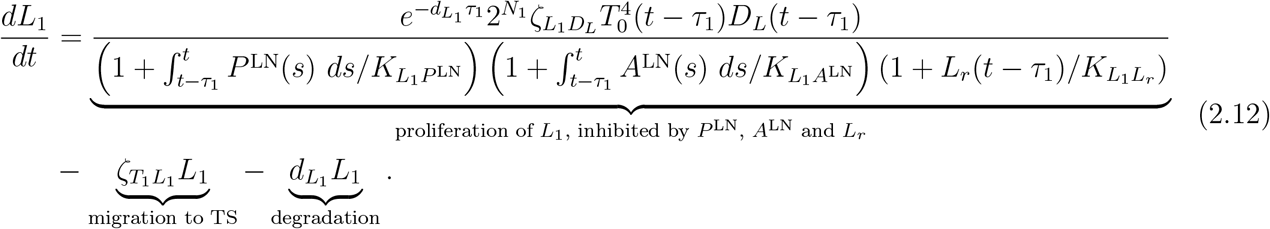

#### 2.2.13 Equation for Effector Th1 Cells in the TS (*T*_1_)

The exponential in the first term accounts for the degradation of effector Th1 cells during the transit of time *τ*_*b*_ from the TS to the TDLN. We account for the volume difference between the TDLN and TS by multiplying this term by *V*_LN_*/V*_TS_ and consider the transit time of DCs arriving in the TS as having left the TDLN at time *t* −*τ*_*b*_. We assume that the degradation rate during transit is the same as the degradation of effector Th1 cells in the TS. The second term accounts for the proliferation of effector Th1 cells in the TS by IL-2 [75]. This gives us

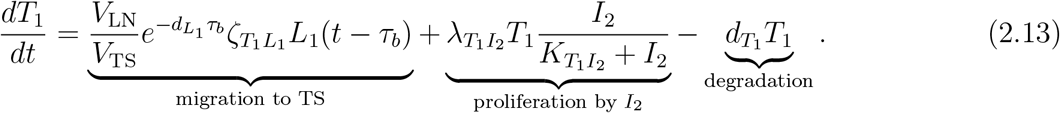

#### 2.2.14 Equation for Effector Th2 Cells in the TDLN (*L*_2_)

The exponential in the first term accounts for the degradation of activated Th2 cells during proliferation dependent on *τ*_2_, the time for *L*_2_ to complete clonal expansion, and *N*_2_, the number of cell divisions in clonal expansion before proliferation stops. We assume that proliferation takes *τ*_2_ time to finish, and that the degradation during proliferation is the same as the degradation of effector Th2 cells in the TDLN. We also consider the activation of 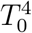 to *L*_2_ by *D*_*L*_ at time *t − τ*_2_, and the inhibition of Th2 cell proliferation due to carboplatin and paclitaxel. The second term accounts for the migration of effector Th2 cells from the TDLN to the TS. This gives us

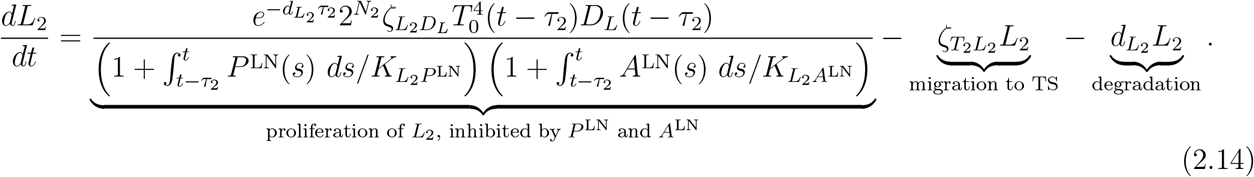

#### 2.2.15 Equation for Effector Th2 Cells in the TS (*T*_2_)

The exponential in the first term accounts for the degradation of activated Th2 cells during the transit of time *τ*_*c*_ from the TS to the TDLN. We account for the volume difference between the TDLN and TS by multiplying this term by *V*_LN_*/V*_TS_. We also consider the transit time of DCs arriving in the TS as having left the TDLN at time *t* −*τ*_*c*_. We assume that the degradation during transit is the same as the degradation of effector Th2 cells in the TS. The second term accounts for the proliferation of effector Th2 cells in the TS by IL-4 [76]. Finally, This gives us

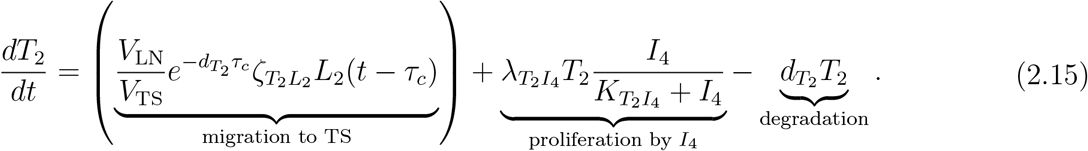

#### 2.2.16 Equation for Naïve Tregs in the TDLN 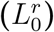

In the first term, we assume that naïve Tregs come from a source 𝒜_*T*_ mainly from the thymus [77]. The second term accounts for the activation of 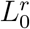by *D*_*L*_. This gives us

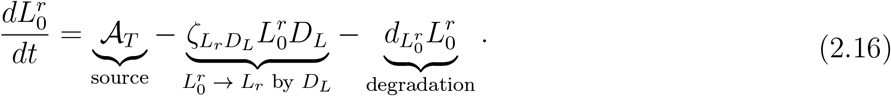

#### 2.2.17 Equation for Effector Tregs in the TDLN (*L*_*r*_)

The exponential in the first term accounts for the degradation of activated Tregs during proliferation dependent on *τ*_*r*_, the time for *L*_*r*_ to complete clonal expansion, and *N*_*r*_, the number of cell divisions in clonal expansion before proliferation stops. We assume that proliferation occurs with a delay of *τ*_*r*_ after activation, and that the degradation during proliferation is the same as the degradation of effector Tregs in the TDLN. We also consider the activation of 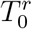 to *L*_*r*_ by *D*_*L*_ at time *t* −*τ*_*r*_, and the inhibition of Treg cell proliferation due to carboplatin and paclitaxel. The second term accounts for the migration of Tregs from the TDLN to the TS. This gives us

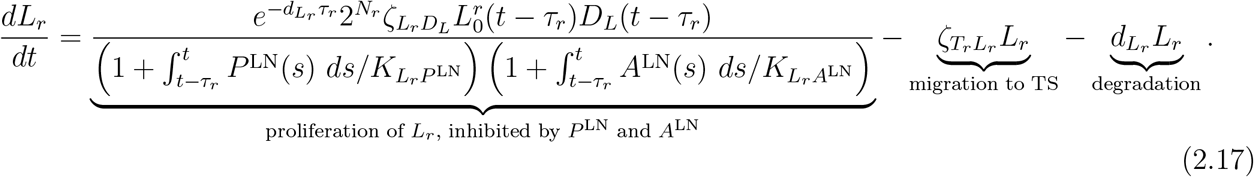

#### 2.2.18 Equation for Effector Tregs in the TS (*T*_*r*_)

The exponential in the first term accounts for the degradation of effector Tregs during the transit of time *τ*_*d*_ from the TS to the TDLN. We account for the volume difference between the TDLN and TS by multiplying this term by *V*_LN_*/V*_TS_. We assume that the degradation during transit is the same as the degradation of effector Tregs in the TS. We also consider the transit time of dendritic cells arriving in the TS as having left the TDLN at time *t* −*τ*_*d*_. The second term accounts for the proliferation of effector Tregs in the TS by TGF-*β* [78]. This gives us

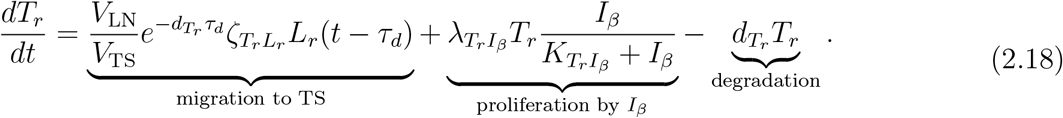

#### 2.2.19 Equations for Naïve (*M*_0_), M1 (*M*_1_) and M2 (*M*_2_) Macrophages

In the first term, we assume that naïve macrophages are supplied at a rate 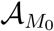. The second and third terms account for IFN-*γ* and TNF stimulating the polarisation of naïve macrophages toward the pro-inflammatory M1 phenotype, respectively, and the fourth and fifth terms account for IL-4 and IL-10 stimulating the polarisation of naïve macrophages toward the anti-inflammatory M2 phenotype [79–81]. This gives us

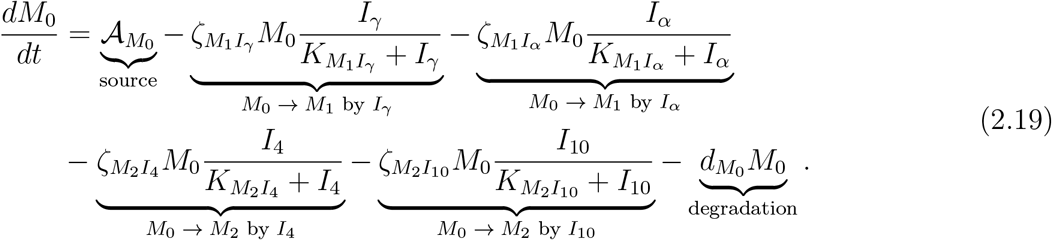

The first and second terms account for IFN-*γ* and TNF stimulating the polarisation of naïve macrophages toward the pro-inflammatory M1 phenotype from (2.19). The third and fourth terms account for IFN-*γ* and TNF stimulating the re-polarisation of M2 macrophages toward the pro-inflammatory M1 phenotype, and the fifth term accounts for TGF-*β* stimulating the re-polarisation of M1 macrophages toward the M2 anti-inflammatory phenotype [82–84]. This gives us

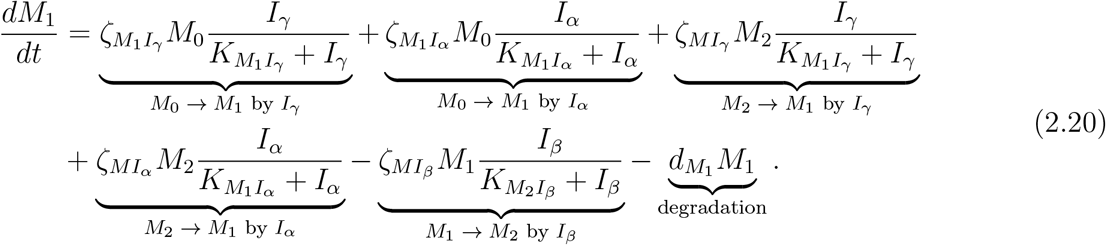

The first and second terms account for IL-4 and IL-10 stimulating the polarisation of naïve macrophages toward the anti-inflammatory M2 phenotype from (2.19). The third and fourth terms account for IFN-*γ* and TNF stimulating the re-polarisation of M2 macrophages toward the pro-inflammatory M1 phenotype from (2.20). The fifth term accounts for TGF-*β* re-polarisation of M1 macrophages toward the anti-inflammatory M2 phenotype from (2.20). This gives us

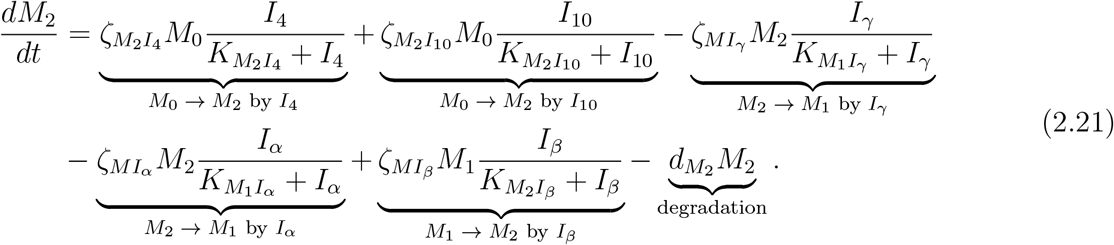

#### 2.2.20 Equations for Naïve (*K*_0_) and Activated (*K*) NK cells

In the first term, we assume naïve NK cells are supplied at a rate 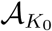. The second and third terms account for NK cell activation by IL-2 and IL-12, respectively [85, 86]. This gives us

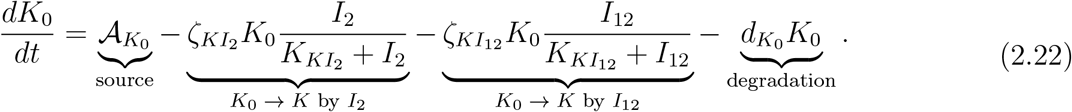

The first and second terms account for NK cell activation by IL-2 and IL-12 from (2.22). IL-2 [85] and IL-12 [87] also promote NK cell proliferation, shown in the third and fourth terms, respectively. This gives us

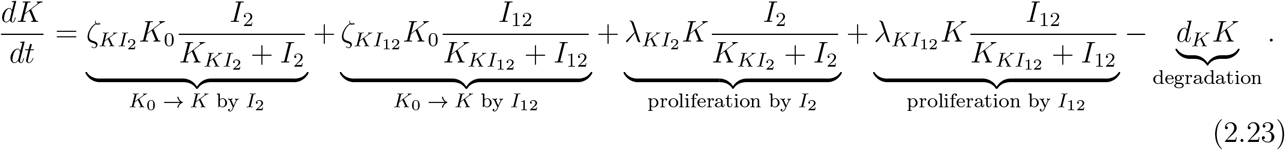

#### 2.2.21 Equation for IL-2 (*I*_2_)

The equation accounts for IL-2 production by CD4+ Th1 and CD8+ T cells inhibited by Tregs [88, 89]. This gives us

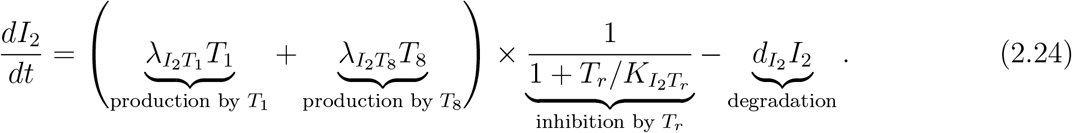

#### 2.2.22 Equation for IL-4 (*I*_4_)

The equation accounts for the release of IL-4 by Th2 CD4+ T cells [80]. This gives us

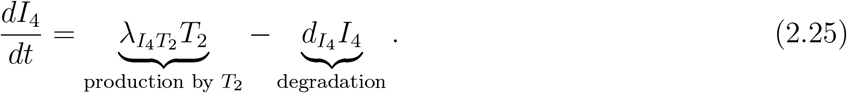

#### 2.2.23 Equation for IL-10 (*I*_10_)

The first to third terms account for IL-10 production by Tregs, M2 macrophages and cancer cells, respectively [80, 90–92]. This gives us

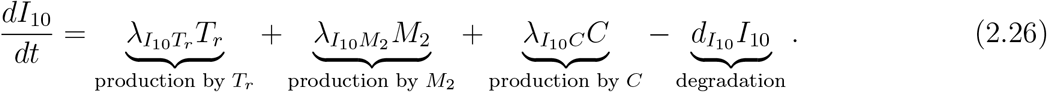

#### 2.2.24 Equation for IL-12 (*I*_12_)

The first and second terms account for IL-12 production by mature DCs and activated M1 macrophages [93–95], respectively. This gives us

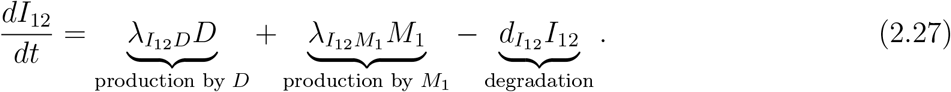

#### 2.2.25 Equation for IFN-*γ* (*I*_*γ*_)

The first term accounts for IFN-*γ* production by CD4+ Th1 and NK cells inhibited by Tregs [80, 87, 96–98]. The second term accounts for IFN-*γ* production by CD8+ T cells [99]. This gives us

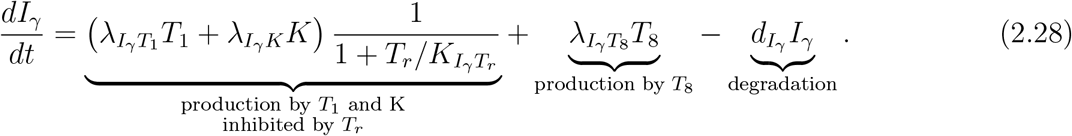

#### 2.2.26 Equation for TNF (*I*_*α*_)

The first to fifth terms account for TNF production by CD8+ T cells, CD4+ Th1 cells, NK cells, M1 polarised macrophages and mature DCs [82, 94, 96, 99–101], respectively. This gives us

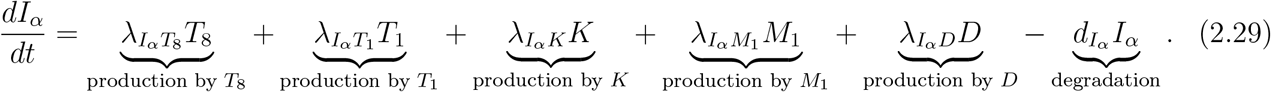

#### 2.2.27 Equation for TGF-*β* (*I*_*β*_)

The first to fourth terms account for TGF-*β* production by Tregs, cancer cells, M2 polarised macrophages and NK cells, respectively [87, 102–105]. This gives us

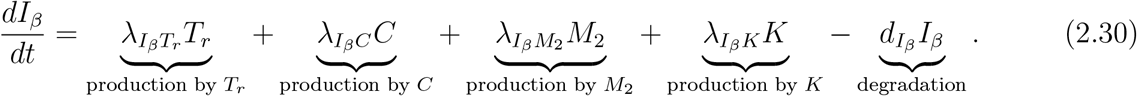

#### 2.2.28 Equations for Paclitaxel (*P* and *P* ^LN^)

We assume that paclitaxel is administered at a constant rate over a duration Δ_*P*_ intravenously at times 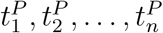 with doses 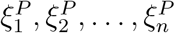 respectively. We account for the systemic nature of paclitaxel distribution and depletion due to killing cancer cells in the TS and killing CD8+ T cells, Th1 and Th2 cells, and Tregs in the TDLN. It is important to note that the administered dose is not equal to the corresponding change in concentration in the TS or the TDLN. For simplicity, we assume linear pharmacokinetics so that, for some scaling factor *f*_*P*_, we have that

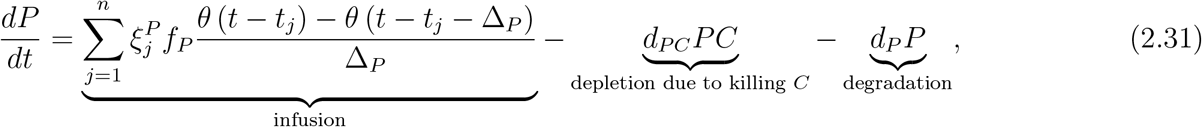

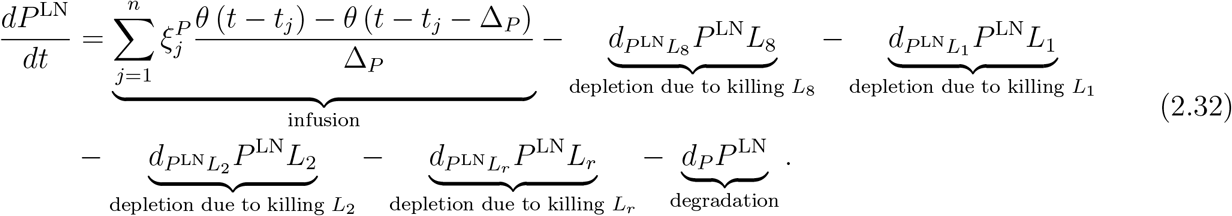

where *θ*(*t*) is the Heaviside function which equals 1 if *t* ≥ 0, and 0 otherwise.

#### 2.2.29 Equations for Carboplatin (*A* and *A*^LN^)

We assume that carboplatin is administered at a constant rate over a duration Δ_*A*_ intravenously at times 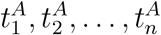 with doses 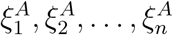 respectively. We account for the systemic nature of carboplatin distribution and depletion due to killing cancer cells in the TS and due to killing CD8+ T cells, Th1 and Th2 cells, and Tregs in the TDLN. It is important to note that the administered dose is not equal to the corresponding change in concentration in the TS or the TDLN. For simplicity, we assume linear pharmacokinetics so that, for some scaling factor *f*_*A*_, we have that

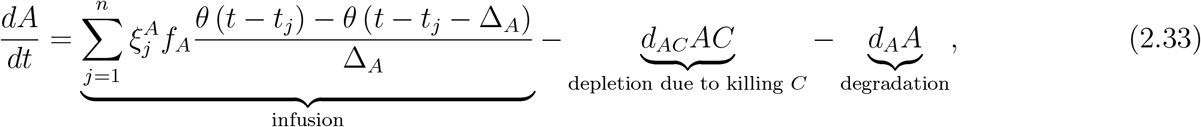

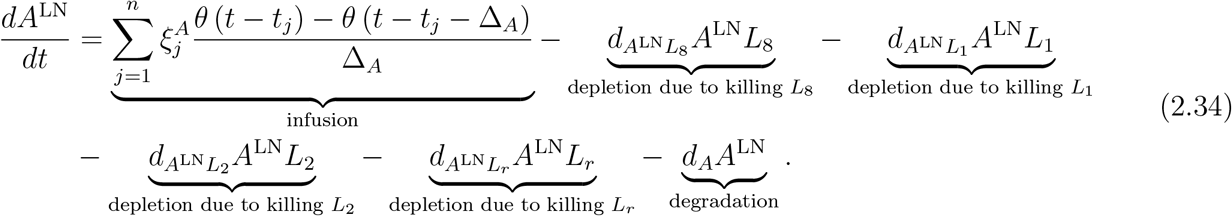

#### 2.2.30 Model Reduction via QSSA

The model parameters are listed in Table 2 and are estimated in Appendix A. We note that the degradation rates of cytokines and DAMPs are generally several orders of magnitude higher than those of immune and cancer cells. Specifically, IL-2, IL-4, IFN-*γ*, TNF, and TGF-*β* evolve on much faster timescales due to their comparatively rapid degradation, causing them to reach equilibrium significantly quicker than other species in the model. Consequently, we apply a quasi-steady state approximation (QSSA) by setting (2.24), (2.25), (2.28), (2.2.26), and (2.2.27) to zero and solving for *I*_2_, *I*_4_, *I*_*γ*_, *I*_*α*_, and *I*_*β*_ as functions of the remaining model variables and parameters. This simplification introduces minimal impact on the system’s dynamics beyond a brief transient phase [106], and we support its validity by noting that system trajectories exhibit negligible variation under nearby parameter sets [36]. Implementing the QSSA results in

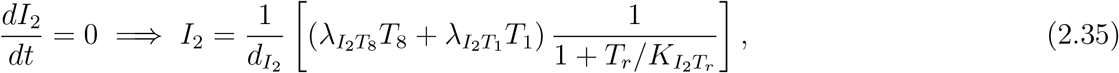

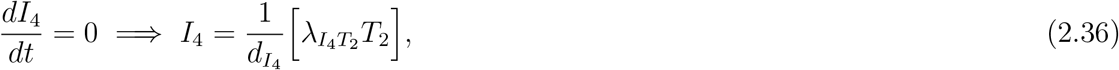

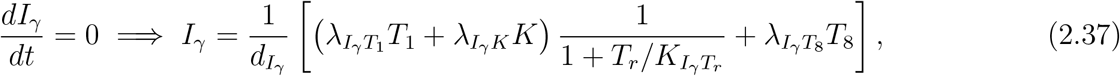

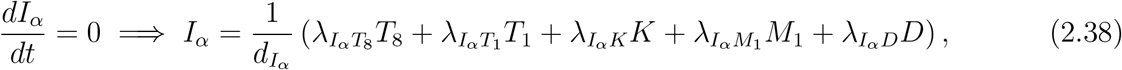

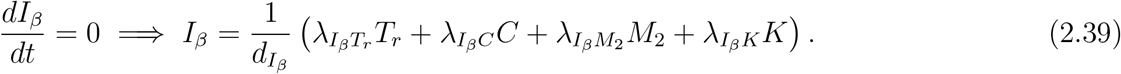

**Table 2:** Parameter values for the model. est. denotes estimated parameters.

| Parameter | Description | Value | Unit | Reference |
| --- | --- | --- | --- | --- |
| $\zeta_{DH}$ | Rate of maturation of $D$ by $H$ | 8.831 | $\text{day}^{-1}$ | est. |
| $\zeta_{DR}$ | Rate of maturation of $D$ by $R$ | 0.883 | $\text{day}^{-1}$ | est. |
| $\zeta_{L_8 D_L}$ | Rate of activation of $T_0^8 \rightarrow L_8$ by $D_L$ | $1.103 \times 10^{-14}$ | $(\text{cell cm}^{-3})^{-1} \text{day}^{-1}$ | est. |
| $\zeta_{T_8 L_8}$ | Migration of $L_8$ to the TS | $2.891 \times 10^{-2}$ | $\text{day}^{-1}$ | est. |
| $\zeta_{L_1 D_L}$ | Rate of activation of $T_0^4 \rightarrow L_1$ by $D_L$ | $2.967 \times 10^{-14}$ | $(\text{cell cm}^{-3})^{-1} \text{day}^{-1}$ | est. |
| $\zeta_{L_2 D_L}$ | Rate of activation of $T_0^4 \rightarrow L_2$ by $D_L$ | $1.517 \times 10^{-14}$ | $(\text{cell cm}^{-3})^{-1} \text{day}^{-1}$ | est. |
| $\zeta_{T_1 L_1}$ | Migration of $L_1$ to the TS | $2.816 \times 10^{-2}$ | $\text{day}^{-1}$ | est. |
| $\zeta_{T_2 L_2}$ | Migration of $L_2$ to the TS | $2.898 \times 10^{-2}$ | $\text{day}^{-1}$ | est. |
| $\zeta_{L_r D_L}$ | Rate of activation of $L_0^r \rightarrow L_r$ by $D_L$ | $1.970 \times 10^{-11}$ | $(\text{cell cm}^{-3})^{-1} \text{day}^{-1}$ | est. |
| $\zeta_{T_r L_r}$ | Migration of $L_r$ to the TS | $1.098 \times 10^{-2}$ | $\text{day}^{-1}$ | est. |
| $\zeta_{M_1 I_\gamma}$ | Rate of polarisation of $M_0 \rightarrow M_1$ by $I_\gamma$ | $5.371 \times 10^{-1}$ | $\text{day}^{-1}$ | [107] est. |
| $\zeta_{M_1 I_\alpha}$ | Rate of polarisation of $M_0 \rightarrow M_1$ by $I_\alpha$ | $4.613 \times 10^{-1}$ | $\text{day}^{-1}$ | [107] est. |
| $\zeta_{M_2 I_4}$ | Rate of polarisation of $M_0 \rightarrow M_2$ by $I_4$ | $3.165 \times 10^{-1}$ | $\text{day}^{-1}$ | [107] est. |
| $\zeta_{M_2 I_{10}}$ | Rate of polarisation of $M_0 \rightarrow M_2$ by $I_{10}$ | $2.197 \times 10^{-1}$ | $\text{day}^{-1}$ | [107] est. |
| $\zeta_{M I_\alpha}$ | Rate of polarisation of $M_2 \rightarrow M_1$ by $I_\alpha$ | $2.154 \times 10^{-2}$ | $\text{day}^{-1}$ | [107] est. |
| $\zeta_{M I_\beta}$ | Rate of polarisation of $M_1 \rightarrow M_2$ by $I_\beta$ | $1.337 \times 10^{-2}$ | $\text{day}^{-1}$ | [107] est. |
| $\zeta_{M I_\gamma}$ | Rate of polarisation of $M_2 \rightarrow M_1$ by $I_\gamma$ | $2.585 \times 10^{-2}$ | $\text{day}^{-1}$ | [107] est. |
| $\zeta_{K I_2}$ | Rate of activation of $K$ by $I_2$ | $2.385 \times 10^{-1}$ | $\text{day}^{-1}$ | [107] est. |
| $\zeta_{K I_{12}}$ | Rate of activation of $K$ by $I_{12}$ | $2.297 \times 10^{-1}$ | $\text{day}^{-1}$ | [107] est. |
| $V_{\text{TS}}$ | Volume of the TS | 51 | $\text{cm}^3$ | [108] |
| $V_{\text{LN}}$ | Volume of the TDLN | $1.85 \times 10^{-1}$ | $\text{cm}^3$ | [109] |
| $\tau_m$ | Time for $D$ to travel to the TDLN | 0.75 | day | [36, 110] est. |
| $\tau_8$ | Time for $L_8$ to complete clonal expansion | 4.85 | day | est. |
| $N_8$ | Number of cell divisions of $L_8$ to complete clonal expansion | 10 | dimensionless | [36, 111, 112] |
| $\tau_a$ | Time for $L_8$ to travel to the TS | 0.27 | day | [113] |
| $\tau_1$ | Time for $L_1$ to complete clonal expansion | 4.13 | day | [36] |
| $N_1$ | Number of cell divisions of $L_1$ to complete clonal expansion | 9 | dimensionless | [114] est. |
| $\tau_b$ | Time for $L_1$ to travel to the TS | 0.27 | day | [113] |
| $\tau_2$ | Time for $L_2$ to complete clonal expansion | 4.13 | day | [36] |
| $N_2$ | Number of cell divisions of $L_2$ to complete clonal expansion | 9 | dimensionless | [114] est. |
| $\tau_c$ | Time for $L_2$ to travel to the TS | 0.27 | day | [113] |
| $\tau_r$ | Time for $L_r$ to complete clonal expansion | 2.87 | day | [36] |
| $N_r$ | Number of cell divisions of $L_r$ to complete clonal expansion | 6 | dimensionless | [115] est. |
| $\tau_d$ | Time for $L_r$ to travel to the TS | 0.27 | day | [113] |
| $r_C$ | Growth constant of $C$ | 0.534 | day<sup>-1</sup> | est. |
| $\lambda_{CP}$ | Rate of apoptosis of $C$ by $P$ | $2.857 \times 10^{-1}$ | day<sup>-1</sup> | est. |
| $\lambda_{CA}$ | Rate of apoptosis of $C$ by $A$ | $7.00 \times 10^{-1}$ | day<sup>-1</sup> | est. |
| $\nu_{CP}$ | Rate of necrosis of $C$ by $P$ | $1.429 \times 10^{-2}$ | day<sup>-1</sup> | est. |
| $\lambda_{CT_8}$ | Rate of lysis of $C$ by $T_8$ | $4.00 \times 10^{-8}$ | (cell/cm<sup>3</sup>)<sup>-1</sup>day<sup>-1</sup> | est. |
| $\lambda_{CT_1}$ | Rate of lysis of $C$ by $T_1$ | $1.25 \times 10^{-9}$ | (cell/cm<sup>3</sup>)<sup>-1</sup>day<sup>-1</sup> | est. |
| $\lambda_{CK}$ | Rate of lysis of $C$ by $K$ | $4.00 \times 10^{-8}$ | (cell/cm<sup>3</sup>)<sup>-1</sup>day<sup>-1</sup> | est. |
| $\lambda_{CI_\alpha}$ | Rate of lysis of $C$ by $I_\alpha$ | 0.704 | day<sup>-1</sup> | est. |
| $\lambda_{CI_\gamma}$ | Rate of lysis of $C$ by $I_\gamma$ | $14.08 \times 10^{-2}$ | day<sup>-1</sup> | est. |
| $\lambda_{NH}$ | Rate of $H$ release by $N$ | $1.263 \times 10^{-12}$ | (g/cell) day<sup>-1</sup> | est. |
| $\lambda_{NR}$ | Rate of $R$ release by $N$ | $2.128 \times 10^{-13}$ | (g/cell) day<sup>-1</sup> | est. |
| $\lambda_{D_L D}$ | Rate of $D$ migration to TDLN | $6.114 \times 10^{-1}$ | day<sup>-1</sup> | est. |
| $\lambda_{T_8 I_2}$ | Rate of proliferation of $T_8$ by $I_2$ | $9 \times 10^{-4}$ | day<sup>-1</sup> | est. |
| $\lambda_{T_1 I_2}$ | Rate of proliferation of $T_1$ by $I_2$ | $7.970 \times 10^{-4}$ | day<sup>-1</sup> | est. |
| $\lambda_{T_2 I_4}$ | Rate of proliferation of $T_2$ by $I_4$ | $3.985 \times 10^{-4}$ | day<sup>-1</sup> | est. |
| $\lambda_{T_r I_\beta}$ | Rate of proliferation of $T_r$ by $I_\beta$ | $6.30 \times 10^{-3}$ | day<sup>-1</sup> | est. |
| $\lambda_{K I_2}$ | Rate of proliferation of $K$ by $I_2$ | $1.518 \times 10^{-3}$ | day<sup>-1</sup> | est. |
| $\lambda_{K I_{12}}$ | Rate of proliferation of $K$ by $I_{12}$ | $1.462 \times 10^{-3}$ | day<sup>-1</sup> | est. |
| $\lambda_{I_2 T_1}$ | Rate of production of $I_2$ by $T_1$ | $9.957 \times 10^{-14}$ | (g/cell) day<sup>-1</sup> | [107] est. |
| $\lambda_{I_2 T_8}$ | Rate of production of $I_2$ by $T_8$ | $3.399 \times 10^{-14}$ | (g/cell) day<sup>-1</sup> | [107] est. |
| $\lambda_{I_4 T_2}$ | Rate of production of $I_4$ by $T_2$ | $8.749 \times 10^{-15}$ | (g/cell) day<sup>-1</sup> | est. |
| $\lambda_{I_{10} T_r}$ | Rate of production of $I_{10}$ by $T_r$ | $2.204 \times 10^{-15}$ | (g/cell) day<sup>-1</sup> | [107] est. |
| $\lambda_{I_{10} M_2}$ | Rate of production of $I_{10}$ by $M_2$ | $4.667 \times 10^{-15}$ | (g/cell) day<sup>-1</sup> | [107] est. |
| $\lambda_{I_{10} C}$ | Rate of production of $I_{10}$ by $C$ | $2.334 \times 10^{-15}$ | (g/cell) day<sup>-1</sup> | [107] est. |
| $\lambda_{I_{12} D}$ | Rate of production of $I_{12}$ by $D$ | $7.05 \times 10^{-18}$ | (g/cell) day<sup>-1</sup> | [107] est. |
| $\lambda_{I_{12} M_1}$ | Rate of production of $I_{12}$ by $M_1$ | $8.03 \times 10^{-18}$ | (g/cell) day<sup>-1</sup> | [107] est. |
| $\lambda_{I_\gamma T_1}$ | Rate of production of $I_\gamma$ by $T_1$ | $4.202 \times 10^{-17}$ | (g/cell) day<sup>-1</sup> | [107] est. |
| $\lambda_{I_\gamma K}$ | Rate of production of $I_\gamma$ by $K$ | $2.224 \times 10^{-15}$ | (g/cell) day<sup>-1</sup> | [107] est. |
| $\lambda_{I_\gamma T_8}$ | Rate of production of $I_\gamma$ by $T_8$ | $6.001 \times 10^{-17}$ | (g/cell) day<sup>-1</sup> | [107] est. |
| $\lambda_{I_\alpha T_8}$ | Rate of production of $I_\alpha$ by $T_8$ | $2.712 \times 10^{-17}$ | (g/cell) day<sup>-1</sup> | [107] est. |
| $\lambda_{I_\alpha T_1}$ | Rate of production of $I_\alpha$ by $T_1$ | $4.483 \times 10^{-17}$ | (g/cell) day<sup>-1</sup> | [107] est. |
| $\lambda_{I_\alpha K}$ | Rate of production of $I_\alpha$ by $K$ | $4.729 \times 10^{-17}$ | (g/cell) day<sup>-1</sup> | [107] est. |
| $\lambda_{I_\alpha M_1}$ | Rate of production of $I_\alpha$ by $M_1$ | $1.643 \times 10^{-17}$ | (g/cell) day<sup>-1</sup> | [107] est. |
| $\lambda_{I_\alpha D}$ | Rate of production of $I_\alpha$ by $D$ | $8.256 \times 10^{-17}$ | (g/cell) day<sup>-1</sup> | [107] est. |
| $\lambda_{I_\beta T_r}$ | Rate of production of $I_\beta$ by $T_r$ | $1.191 \times 10^{-12}$ | (g/cell) day<sup>-1</sup> | [107] est. |
| $\lambda_{I_\beta C}$ | Rate of production of $I_\beta$ by $C$ | $2.057 \times 10^{-13}$ | (g/cell) day<sup>-1</sup> | est. |
| $\lambda_{I_\beta M_2}$ | Rate of production of $I_\beta$ by $M_2$ | $1.48 \times 10^{-12}$ | (g/cell) day<sup>-1</sup> | [107] est. |
| $\lambda_{I_\beta K}$ | Rate of production of $I_\beta$ by $K$ | $1.007 \times 10^{-12}$ | (g/cell) day<sup>-1</sup> | [107] est. |
| $f_P$ | $P/P^{\text{LN}}$ dose scaling factor | $1.043 \times 10^{-8}$ | g cm<sup>-3</sup> mg<sup>-1</sup> | [116] est. |
| $f_A$ | $A/A^{\text{LN}}$ dose scaling factor | $4.229 \times 10^{-8}$ | g cm<sup>-3</sup> mg<sup>-1</sup> | [116] est. |
| $K_{CP}$ | Half-saturation of $P$ for $C$ | $3.064 \times 10^{-7}$ | g cm<sup>-3</sup> | [116] est. |
| $K_{CA}$ | Half-saturation of $A$ for $C$ | $3.713 \times 10^{-6}$ | g cm<sup>-3</sup> | [116] est. |
| $K_{CI_\alpha}$ | Half-saturation of $I_\alpha$ for $C$ | $2.62 \times 10^{-12}$ | g cm<sup>-3</sup> | [117] est. |
| $K_{CI_\gamma}$ | Half-saturation of $I_\gamma$ for $C$ | $5.2 \times 10^{-11}$ | g cm<sup>-3</sup> | [117] est. |
| $K_{DH}$ | Half-saturation of $H$ for $D$ | $6.69 \times 10^{-8}$ | $\text{g cm}^{-3}$ | [118] est. |
| $K_{DR}$ | Half-saturation of $H$ for $R$ | $4.5 \times 10^{-8}$ | $\text{g cm}^{-3}$ | [119] est. |
| $K_{T_8 I_2}$ | Half-saturation of $I_2$ for $T_8$ | $2.91 \times 10^{-11}$ | $\text{g cm}^{-3}$ | [120] est. |
| $K_{T_1 I_2}$ | Half-saturation of $I_2$ for $T_1$ | $2.91 \times 10^{-11}$ | $\text{cell cm}^{-3}$ | [121] est. |
| $K_{T_2 I_4}$ | Half-saturation of $I_4$ for $T_2$ | $9.46 \times 10^{-12}$ | $\text{g cm}^{-3}$ | [122] est. |
| $K_{T_r I_\beta}$ | Half-saturation of $I_\beta$ for $T_r$ | $1.75 \times 10^{-8}$ | $\text{g cm}^{-3}$ | [123, 124] est. |
| $K_{M_1 I_\gamma}$ | Half-saturation of $I_\gamma$ for $M_1$ | $5.2 \times 10^{-11}$ | $\text{g cm}^{-3}$ | [117] est. |
| $K_{M_1 I_\alpha}$ | Half-saturation of $I_\alpha$ for $M_1$ | $2.62 \times 10^{-12}$ | $\text{g cm}^{-3}$ | [117] est. |
| $K_{M_2 I_4}$ | Half-saturation of $I_4$ for $M_2$ | $9.46 \times 10^{-12}$ | $\text{g cm}^{-3}$ | [122] est. |
| $K_{M_2 I_{10}}$ | Half-saturation of $I_{10}$ for $M_2$ | $1.1 \times 10^{-8}$ | $\text{g cm}^{-3}$ | [125, 126] est. |
| $K_{M_2 I_\beta}$ | Half-saturation of $I_\beta$ for $M_2$ | $1.75 \times 10^{-8}$ | $\text{g cm}^{-3}$ | [123, 124] est. |
| $K_{K I_2}$ | Half-saturation of $I_2$ for $K$ | $2.91 \times 10^{-11}$ | $\text{g cm}^{-3}$ | [120] est. |
| $K_{K I_{12}}$ | Half-saturation of $I_{12}$ for $K$ | $3.1 \times 10^{-12}$ | $\text{g cm}^{-3}$ | [127] est. |
| $C_0$ | Carrying capacity of $C$ | $1.75 \times 10^8$ | $\text{cell cm}^{-3}$ | est. |
| $K_{T_8 I_\beta}$ | Inhibition constant of $T_8$ elimination of $C$ by $I_\beta$ | $1.75 \times 10^{-8}$ | $\text{g cm}^{-3}$ | [123, 124] est. |
| $K_{T_8 T_r}$ | Inhibition constant of $T_8$ elimination of $C$ by $T_r$ | $1.25 \times 10^5$ | $\text{cell cm}^{-3}$ | [121] est. |
| $K_{T_1 T_r}$ | Inhibition constant of $T_1$ elimination of $C$ by $T_r$ | $1.25 \times 10^5$ | $\text{cell cm}^{-3}$ | [121] est. |
| $K_{K I_\beta}$ | Inhibition constant for $K$ elimination of $C$ by $I_\beta$ | $1.75 \times 10^{-8}$ | $\text{g cm}^{-3}$ | [123, 124] est. |
| $K_{K T_r}$ | Inhibition constant for $K$ elimination of $C$ by $T_r$ | $1.25 \times 10^5$ | $\text{cell cm}^{-3}$ | [121] est. |
| $K_{D I_{10}}$ | Inhibition constant for $D_0$ maturation by $I_{10}$ | $1.1 \times 10^{-8}$ | $\text{g cm}^{-3}$ | [125, 126] est. |
| $K_{D I_\beta}$ | Inhibition constant for $D_0$ maturation by $I_\beta$ | $1.75 \times 10^{-8}$ | $\text{g cm}^{-3}$ | [123, 124] est. |
| $K_{T_0^8 L_r}$ | Inhibition constant for $T_0^8$ activation by $L_r$ | $1.91 \times 10^8$ | $\text{cell cm}^{-3}$ | [121] est. |
| $K_{L_8 P^{LN}}$ | Inhibition constant of activated CD8+ T cell proliferation by $P^{LN}$ | $4.995 \times 10^{-7}$ | $(\text{g cm}^{-3}) \text{ day}$ | [116] est. |
| $K_{L_8 A^{LN}}$ | Inhibition constant of activated CD8+ T cell proliferation by $A^{LN}$ | $6.053 \times 10^{-6}$ | $(\text{g cm}^{-3}) \text{ day}$ | [116] est. |
| $K_{L_8 L_r}$ | Inhibition constant for activated CD8+ T cell proliferation by $L_r$ | $1.91 \times 10^8$ | $\text{cell cm}^{-3}$ | [121] est. |
| $K_{T_8 I_{10}}$ | Inhibition constant of $T_8$ death by $I_{10}$ | $1.1 \times 10^{-8}$ | $\text{g cm}^{-3}$ | [125, 126] est. |
| $K_{T_0^4 L_r}$ | Inhibition constant for $T_0^4$ activation by $L_r$ | $1.91 \times 10^8$ | $\text{cell cm}^{-3}$ | [121] est. |
| $K_{L_1 P^{LN}}$ | Inhibition constant of activated Th1 cell proliferation by $P^{LN}$ | $1.266 \times 10^{-6}$ | $(\text{g cm}^{-3}) \text{ day}$ | [116] est. |
| $K_{L_1 A^{LN}}$ | Inhibition constant of activated Th1 cell proliferation by $A^{LN}$ | $1.534 \times 10^{-5}$ | $(\text{g cm}^{-3}) \text{ day}$ | [116] est. |
| $K_{L_1 L_r}$ | Inhibition constant for activated Th1 cell proliferation by $L_r$ | $1.91 \times 10^8$ | $\text{cell cm}^{-3}$ | [121] est. |
| $K_{L_2 P^{LN}}$ | Inhibition constant of activated Th2 cell proliferation by $P^{LN}$ | $1.266 \times 10^{-6}$ | $(\text{g cm}^{-3}) \text{ day}$ | [116] est. |
| $K_{L_2 A^{LN}}$ | Inhibition constant of activated Th2 cell proliferation by $A^{LN}$ | $1.534 \times 10^{-5}$ | $(g \text{ cm}^{-3}) \text{ day}$ | [116] est. |
| $K_{L_r P^{LN}}$ | Inhibition constant of activated Treg proliferation by $P^{LN}$ | $8.795 \times 10^{-7}$ | $(g \text{ cm}^{-3}) \text{ day}$ | [116] est. |
| $K_{L_r A^{LN}}$ | Inhibition constant of activated Treg proliferation by $A^{LN}$ | $1.066 \times 10^{-5}$ | $(g \text{ cm}^{-3}) \text{ day}$ | [116] est. |
| $K_{I_2 T_r}$ | Inhibition constant of production of $I_2$ by $T_r$ | $1.25 \times 10^5$ | $\text{cell cm}^{-3}$ | [121] est. |
| $K_{I_\gamma T_r}$ | Inhibition constant of production of $I_\gamma$ by $T_r$ | $1.25 \times 10^5$ | $\text{cell cm}^{-3}$ | [121] est. |
| $\mathcal{A}_{D_0}$ | Source of $D_0$ | $7.20 \times 10^5$ | $\text{cell cm}^{-3} \text{ day}^{-1}$ | [128] est. |
| $\mathcal{A}_T$ | Source of $T_r$ | $3.572 \times 10^5$ | $\text{cell cm}^{-3} \text{ day}^{-1}$ | est. |
| $\mathcal{A}_{T_8^0}$ | Source of $T_8^0$ | $2.486 \times 10^6$ | $\text{cell cm}^{-3} \text{ day}^{-1}$ | est. |
| $\mathcal{A}_{T_4^0}$ | Source of $T_4^0$ | $2.547 \times 10^6$ | $\text{cell cm}^{-3} \text{ day}^{-1}$ | est. |
| $\mathcal{A}_{M_0}$ | Source of $M_0$ | $5.551 \times 10^5$ | $\text{cell cm}^{-3} \text{ day}^{-1}$ | est. |
| $\mathcal{A}_{K_0}$ | Source of $K_0$ | $1.362 \times 10^5$ | $\text{cell cm}^{-3} \text{ day}^{-1}$ | est. |
| $d_N$ | Degradation rate of $N$ | $4.81 \times 10^1$ | $\text{day}^{-1}$ | est. |
| $d_H$ | Degradation rate of $H$ | 5.55 | $\text{day}^{-1}$ | [129] |
| $d_R$ | Degradation rate of $R$ | 1.39 | $\text{day}^{-1}$ | [130] |
| $d_{D_0}$ | Degradation rate of $D_0$ | $3.57 \times 10^{-2}$ | $\text{day}^{-1}$ | [131] |
| $d_D$ | Degradation rate of $D$ | $3.15 \times 10^{-1}$ | $\text{day}^{-1}$ | [132, 133] |
| $d_{D_L}$ | Degradation rate of $D_L$ | $3.15 \times 10^{-1}$ | $\text{day}^{-1}$ | [132] |
| $d_{T_8^0}$ | Degradation rate of $T_8^0$ | $3.22 \times 10^{-2}$ | $\text{day}^{-1}$ | [134] |
| $d_{L_8}$ | Degradation rate of $L_8$ | $9.0 \times 10^{-3}$ | $\text{day}^{-1}$ | [135] |
| $d_{T_8}$ | Degradation rate of $T_8$ | $9.0 \times 10^{-3}$ | $\text{day}^{-1}$ | [135] |
| $d_{T_4^0}$ | Degradation rate of $T_4^0$ | $4.03 \times 10^{-2}$ | $\text{day}^{-1}$ | [134] |
| $d_{L_1}$ | Degradation rate of $L_1$ | $7.97 \times 10^{-3}$ | $\text{day}^{-1}$ | [135] |
| $d_{T_1}$ | Degradation rate of $T_1$ | $7.97 \times 10^{-3}$ | $\text{day}^{-1}$ | [135] |
| $d_{L_2}$ | Degradation rate of $L_2$ | $7.97 \times 10^{-3}$ | $\text{day}^{-1}$ | [135] |
| $d_{T_2}$ | Degradation rate of $T_2$ | $7.97 \times 10^{-3}$ | $\text{day}^{-1}$ | [135] |
| $d_{L_0^r}$ | Degradation rate of $L_0^r$ | $2.2 \times 10^{-3}$ | $\text{day}^{-1}$ | [136] |
| $d_{L_r}$ | Degradation rate of $L_r$ | $6.3 \times 10^{-2}$ | $\text{day}^{-1}$ | [137] |
| $d_{T_r}$ | Degradation rate of $T_r$ | $6.3 \times 10^{-2}$ | $\text{day}^{-1}$ | [137] |
| $d_{M_0}$ | Degradation rate of $M_0$ | $7.3 \times 10^{-1}$ | $\text{day}^{-1}$ | [138] |
| $d_{M_1}$ | Degradation rate of $M_1$ | $9.9 \times 10^{-1}$ | $\text{day}^{-1}$ | [138, 139] |
| $d_{M_2}$ | Degradation rate of $M_2$ | $1.35 \times 10^{-1}$ | $\text{day}^{-1}$ | [138] |
| $d_{K_0}$ | Degradation rate of $K_0$ | $6.93 \times 10^{-2}$ | $\text{day}^{-1}$ | [112, 140] |
| $d_K$ | Degradation rate of $K$ | $6.93 \times 10^{-2}$ | $\text{day}^{-1}$ | [112, 140] |
| $d_{I_2}$ | Degradation rate of $I_2$ | $1.446 \times 10^2$ | $\text{day}^{-1}$ | [141, 142] |
| $d_{I_4}$ | Degradation rate of $I_4$ | $5.253 \times 10^1$ | $\text{day}^{-1}$ | [143] |
| $d_{I_{10}}$ | Degradation rate of $I_{10}$ | 6.16 | $\text{day}^{-1}$ | [144, 145] |
| $d_{I_{12}}$ | Degradation rate of $I_{12}$ | 2.23 | $\text{day}^{-1}$ | [146] |
| $d_{I_\gamma}$ | Degradation rate of $I_\gamma$ | $3.327 \times 10^1$ | $\text{day}^{-1}$ | [147] |
| $d_{I_\alpha}$ | Degradation rate of $I_\alpha$ | $5.484 \times 10^1$ | $\text{day}^{-1}$ | [148, 149] |
| $d_{I_\beta}$ | Degradation rate of $I_\beta$ | $4.991 \times 10^2$ | $\text{day}^{-1}$ | [150, 151] |
| $d_{PC}$ | Depletion rate of $P$ by killing $C$ | $5.196 \times 10^{-7}$ | $(\text{cell/cm}^3)^{-1} \text{day}^{-1}$ | est. |
| $d_P$ | Degradation rate of $P$ and $P^{LN}$ | 1.68 | $\text{day}^{-1}$ | [152] |
| $d_{P^{LN}L_8}$ | Depletion rate of $P^{LN}$ by killing $L_8$ | $3.347 \times 10^{-9}$ | $(\text{cell}/\text{cm}^3)^{-1}\text{day}^{-1}$ | est. |
| $d_{P^{LN}L_1}$ | Depletion rate of $P^{LN}$ by killing $L_1$ | $3.012 \times 10^{-9}$ | $(\text{cell}/\text{cm}^3)^{-1}\text{day}^{-1}$ | est. |
| $d_{P^{LN}L_2}$ | Depletion rate of $P^{LN}$ by killing $L_2$ | $3.012 \times 10^{-9}$ | $(\text{cell}/\text{cm}^3)^{-1}\text{day}^{-1}$ | est. |
| $d_{P^{LN}L_r}$ | Depletion rate of $P^{LN}$ by killing $L_r$ | $2.008 \times 10^{-9}$ | $(\text{cell}/\text{cm}^3)^{-1}\text{day}^{-1}$ | est. |
| $d_{AC}$ | Depletion rate of $A$ by killing $C$ | $5.184 \times 10^{-8}$ | $(\text{cell}/\text{cm}^3)^{-1}\text{day}^{-1}$ | est. |
| $d_A$ | Degradation rate of $A$ and $A^{LN}$ | $1.676 \times 10^{-1}$ | $\text{day}^{-1}$ | [153] est. |
| $d_{A^{LN}L_8}$ | Depletion rate of $A^{LN}$ by killing $L_8$ | $3.339 \times 10^{-10}$ | $(\text{cell}/\text{cm}^3)^{-1}\text{day}^{-1}$ | est. |
| $d_{A^{LN}L_1}$ | Depletion rate of $A^{LN}$ by killing $L_1$ | $3.005 \times 10^{-10}$ | $(\text{cell}/\text{cm}^3)^{-1}\text{day}^{-1}$ | est. |
| $d_{A^{LN}L_2}$ | Depletion rate of $A^{LN}$ by killing $L_2$ | $3.005 \times 10^{-10}$ | $(\text{cell}/\text{cm}^3)^{-1}\text{day}^{-1}$ | est. |
| $d_{A^{LN}L_r}$ | Depletion rate of $A^{LN}$ by killing $L_r$ | $2.003 \times 10^{-10}$ | $(\text{cell}/\text{cm}^3)^{-1}\text{day}^{-1}$ | est. |

We note that this reduction holds since IFN-*γ* (the slowest fast species) still evolves much faster than any slow species in the model [106].

### 2.3 Model Parameters

The model parameters are in Table 2 as estimated in Appendix A.

## 3 Initial Conditions and Steady States

Biologically, we assumed a steady state occurred in Stage IV HGSOC patients due to a maladaptive TME where immune regulation becomes dysfunctional. We chose Stage III HGSOC patients as our initial condition, as most patients are diagnosed at this stage with the cell steady states and initial conditions to be as in Table 3. The justification for the choice of these values is shown in Appendix A.1.

**Table 3:**
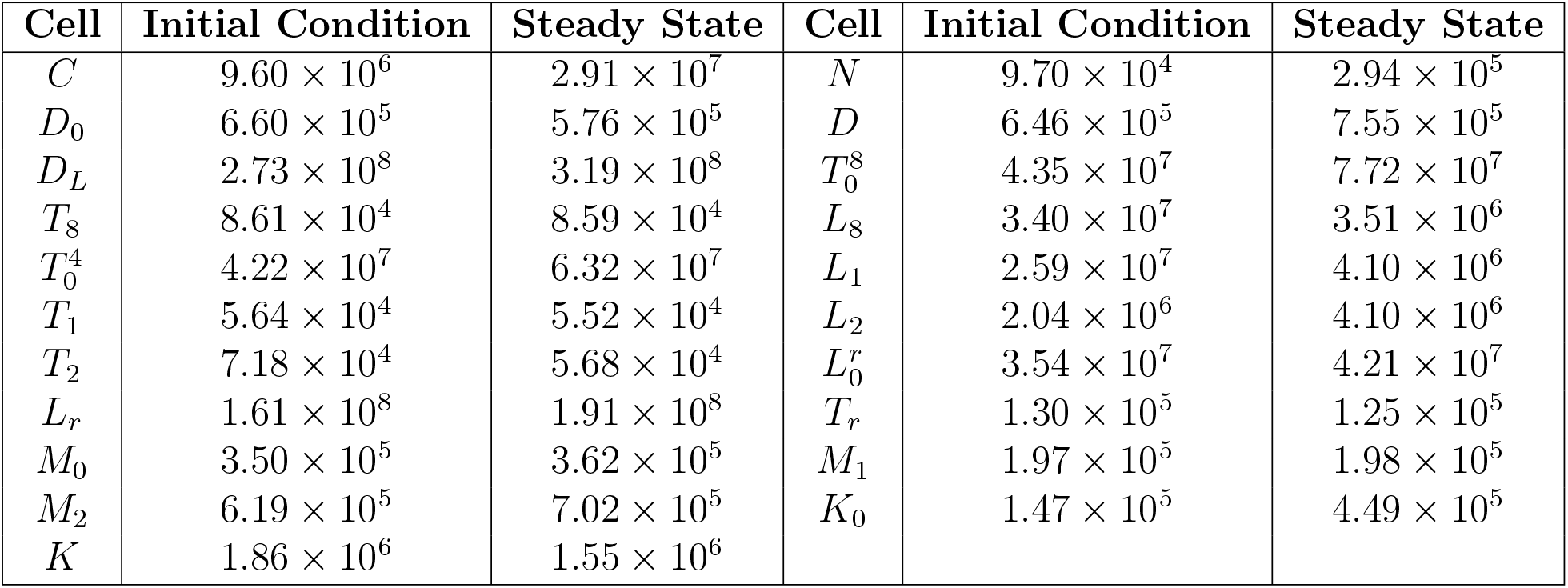
Cell steady states and initial conditions for the model. All values are in units of cells*/*cm^3^.

We chose the DAMP steady states and initial conditions to be as in Table 4. The justification for these values is explained in Appendix A.2.

**Table 4:** DAMP steady states and initial conditions for the model. All values are in units of g*/*cm^3^.

| DAMP | Initial Condition | Steady State |
| --- | --- | --- |
| $H$ | $3.15 \times 10^{-8}$ | $6.69 \times 10^{-8}$ |
| $R$ | $2.00 \times 10^{-8}$ | $4.50 \times 10^{-8}$ |

We choose the cytokine steady states and initial conditions as in Table 5. The choice of these values is explained in Appendix A.3.

**Table 5:** Cytokine initial conditions and steady states for the model. All values are in units of g*/*cm^3^.

| Cytokine | Initial Condition | Steady State |
| --- | --- | --- |
| $I_2$ | $3.05 \times 10^{-11}$ | $2.91 \times 10^{-11}$ |
| $I_4$ | $8.11 \times 10^{-12}$ | $9.46 \times 10^{-12}$ |
| $I_{10}$ | $1.03 \times 10^{-8}$ | $1.16 \times 10^{-8}$ |
| $I_{12}$ | $1.96 \times 10^{-11}$ | $3.10 \times 10^{-12}$ |
| $I_\alpha$ | $2.72 \times 10^{-12}$ | $2.62 \times 10^{-12}$ |
| $I_\gamma$ | $6.11 \times 10^{-11}$ | $5.20 \times 10^{-11}$ |
| $I_\beta$ | $1.48 \times 10^{-8}$ | $1.75 \times 10^{-8}$ |

## 4 Results

We now aim to optimise neoadjuvant paclitaxel and carboplatin therapy in advanced HGSOC. For simplicity, we assume that paclitaxel and carboplatin are given at a constant dosage and that the spacing between consecutive infusions is constant. We also assume that the patient has paclitaxel and carboplatin at *t* = 0 days, and we consider a treatment regimen lasting for 12 weeks so that the time for the latest allowed infusion occurs before *t* = 84 days, and simulate to 84 days. In our optimisation, we consider the following four objectives: efficacy, tumour volume reduction (TVR), efficiency, and toxicity.

First, we use the Calvert formula to determine the appropriate dose of carboplatin (in mg) based on the desired AUC (in mg*/*mL min) and the patient’s glomerular filtration rate (GFR) (in mL*/*min*/*1.73m^2^), taking into account the patient’s renal function [154]:

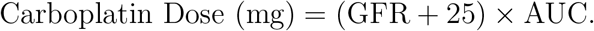

The mean GFR in OC patients was found to be 73.2 mL*/*min*/*1.73m^2^ [155], hence we take

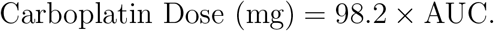

We define *V* (*t*) = *C*(*t*)+*N* (*t*) as the total cancer concentration at time *t* without treatment, and define *V* (*ξ*_*P*_; *η*_*P*_; *ξ*_*A*_; *η*_*A*_, *t*) = *C*(*ξ*_*P*_; *η*_*P*_; *ξ*_*A*_; *η*_*A*_, *t*) + *N* (*ξ*_*P*_; *η*_*P*_; *ξ*_*A*_; *η*_*A*_, *t*) as the total cancer concentration at time *t* with treatment with paclitaxel doses of *ξ*_*P*_ mg*/*m^2^ at a dosing interval of *η*_*P*_ days and carboplatin doses with a target AUC of *ξ*_*A*_ mg*/*mL min at a dosing interval of *η*_*A*_ days. We define the efficacy at time *t* from this regimen to be

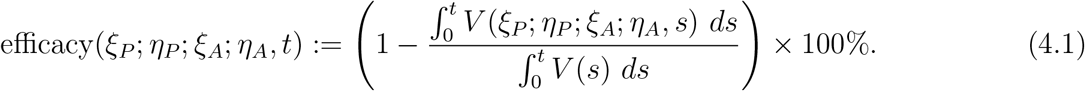

We also define the TVR similarly, as

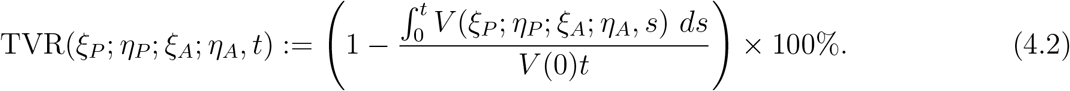

In particular, the efficacy represents the extent of tumour density shrinkage throughout its growth course in comparison to no treatment, whereas the TVR reveals how much the tumour density has reduced since the commencement of treatment. We see that the TVR and efficacy are linearly related via the formula

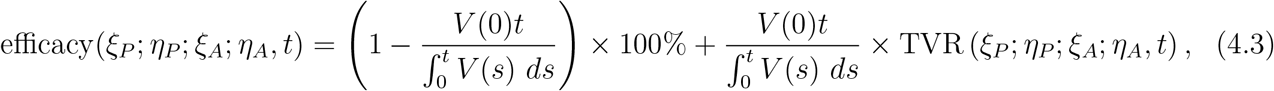

so that an increase in treatment efficacy results in increased TVR, and vice versa. As such, we calculate only the TVR, and the corresponding treatment efficacy can be calculated via (4.3).

We can also consider the efficiency of the treatment regimen, given by

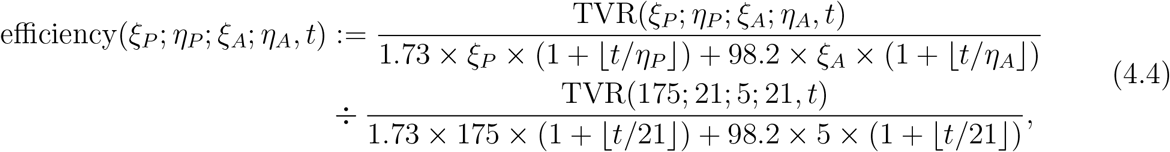

where 1.73 *× ξ*_*P*_ *×* (1 + ⌊*t/η*_*P*_⌋) and 98.2 ×*ξ*_*A*_ × (1 + ⌊*t/η*_*A*_⌋) are the total doses of paclitaxel and carboplatin, in mg, administered by time *t*, respectively, assuming a body surface area of 1.73 m^2^ and a GFR of 73.2 mL*/*min*/*1.73m^2^. The efficiency corresponds to the ratio of the TVR percentage to the total chemotherapy dose administered, comparing the treatment regimen to the standard regimen, with an efficiency greater than 1 denoting a treatment more efficient than the standard regimen and vice versa.

Finally, we can define the toxicity of the treatment regimen, noting that large enough paclitaxel and carboplatin concentrations can potentially cause peripheral neuropathy, bone marrow suppression and cardiotoxicity [156–158]. It was shown in [154] that single-agent carboplatin given at an AUC between 6–8 mg*/*mL min in previously untreated patients led to manageable haematological toxicity. It was also shown in [159] that a regimen of 550 mg*/*m^2^ of carboplatin and 200 mg*/*m^2^ of paclitaxel, administered triweekly for six courses, represents the maximum tolerated dose for previously untreated OC patients. As such, we assume that the threshold for treatment toxicity is 200 mg*/*m^2^ of paclitaxel and an AUC of 7 mg*/*mL min of carboplatin every 3 weeks, with higher doses being deemed toxic. To rigorise this notion, we define the toxicity of the treatment regimen as

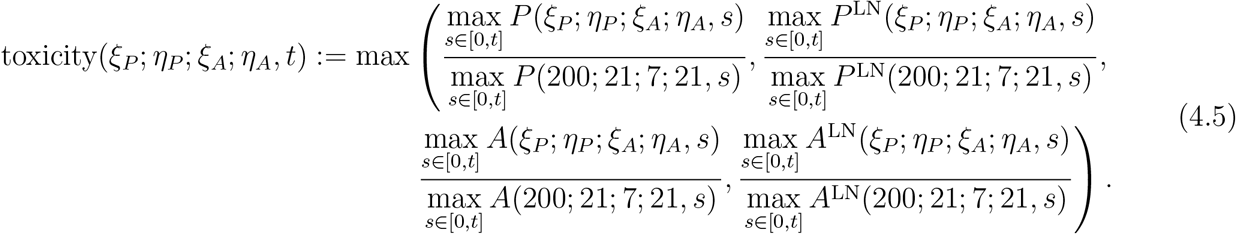

In particular, *P* (*ξ*_*P*_; *η*_*P*_; *ξ*_*A*_; *η*_*A*_, *s*) and *A*(*ξ*_*P*_; *η*_*P*_; *ξ*_*A*_; *η*_*A*_, *s*) denote the concentrations of carboplatin and paclitaxel in the TS at time *s*, respectively. In particular, the toxicity quantifies the ratio of the maximum paclitaxel and carboplatin concentrations from the regimen to those of the threshold regimen, taking the highest value of this ratio between the TDLN and TS. A toxicity greater than 1 indicates a toxic and unsafe regimen, whereas a toxicity of 1 or less signifies a non-toxic and safe regimen, with lower toxicity values corresponding to safer treatments.

We perform a sweep across the space *η*_*P*_, *η*_*A*_ ∈ {7, 14, 21, 28, 42} days. These values are integer factors of 84 days, corresponding to a distinct number of paclitaxel or carboplatin doses administered. This approach ensures practicality whilst preventing any artefacts that could occur from selecting a treatment regimen that ends at a fixed time of 12 weeks. We consider linearly spaced dosages in the domain *ξ*_P_ ∈ [0 mg*/*m^2^, 200 mg*/*m^2^] with an increment of 5 mg*/*m^2^ and *ξ*_*A*_ ∈ [0 AUC, 7 AUC] with an increment of 0.05 AUC, assuming a patient body surface area of 1.73 m^2^ and GFR of 73.2 mL*/*min*/*1.73m^2^.

Heatmaps of TVR, efficacy, efficiency, and toxicity at 12 weeks for these *η*_*P*_, *η*_*A*_, *ξ*_*P*_ and *ξ*_*A*_ values are shown in Appendix B. All simulations were done in MATLAB using the dde23 solver with the initial conditions stated in Section 3 and drug parameters as above.

We can determine the optimal paclitaxel and carboplatin therapies by considering the regimens that achieve an acceptable TVR at 12 weeks, whilst maximising treatment efficiency as much as possible and ensuring toxicity of less than 1. To ensure that the TVR of the optimal treatments is comparable to the SR, we set the threshold TVR to be 75%. Denoting the space of regimens, (*P* ^reg^, *A*^reg^), that satisfy these criteria as *S*^prac^, the optimal regimens, (*P* ^opt^, *A*^opt^), satisfy

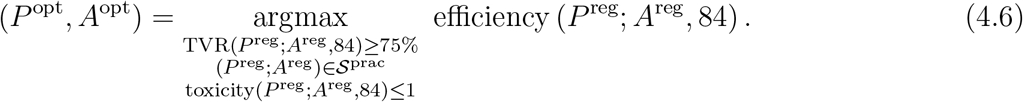

Solving leads to the optimal therapies defined in Table 6. We define a cyclic regimen, such as the 21-day cyclic regimen 101, to indicate treatment on day 1, no treatment on day 8 and treatment on day 15, and similarly for 28-day cyclic regimens.

**Table 6:** Comparison of *ξ*_*P*_, *ξ*_*A*_, TVR, efficacy, efficiency and toxicity for the optimal chemotherapy regimens. *Dose given on Day 1 and Day 8. **Dose given on Day 1, 8, 15.

| Regimen | $\xi_P$<br>(mg/m<sup>2</sup>) | $\xi_A$<br>(AUC) | TVR<br>(%) | Efficacy<br>(%) | Efficiency | Toxicity |
| --- | --- | --- | --- | --- | --- | --- |
| Standard Regimen | 175 | 5.00 | 75.85 | 89.86 | 1.00 | 0.87 |
| Optimal Triweekly | 0 | 5.75 | 75.07 | 89.54 | 1.39 | 0.82 |
| Optimal Biweekly | 0 | 3.30 | 75.46 | 89.70 | 1.62 | 0.50 |
| Optimal Weekly | 0 | 1.40 | 75.83 | 89.86 | 1.92 | 0.28 |
| Optimal 21-day cycle* | 0 | 2.00 | 75.16 | 89.57 | 2.00 | 0.37 |
| Optimal 28-day cycle** | 0 | 1.70 | 75.79 | 89.84 | 2.11 | 0.33 |

Time traces for total cancer concentration (*V*) under optimal triweekly, biweekly, weekly, 21-day cycle and 28-day cycle paclitaxel and carboplatin therapies, compared to no treatment, are shown in Figure 3. We can also compare the effect of optimal triweekly, biweekly, weekly, 21-day cyclic and 28-day cyclic regimens against no treatment, on the TME, with time traces of key variables shown in Figure 4. Figure 5 breaks down the treatment regimes into TVR, efficacy, efficiency, and toxicity. For the cycle regimens, we present heatmaps of TVR, efficacy, efficiency, and toxicity in Figure 6.

**Figure 3:**
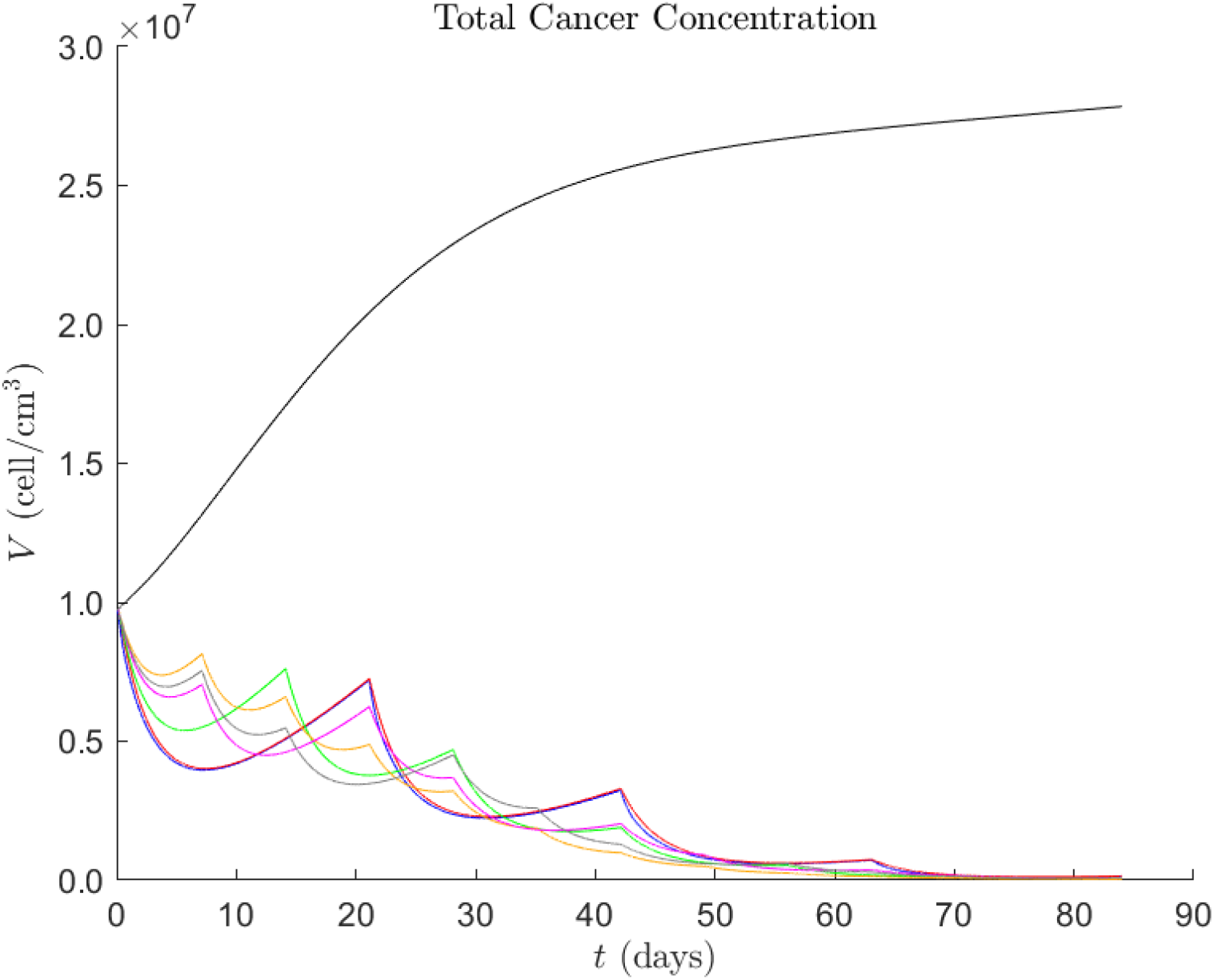
Left: time traces of *V* up to 84 days from commencement, with no treatment shown in black, SR in blue, optimal triweekly in red, optimal biweekly in green, optimal weekly in orange, optimal 21-day cycle in magenta and optimal 28-day cycle in grey.

**Figure 4:**
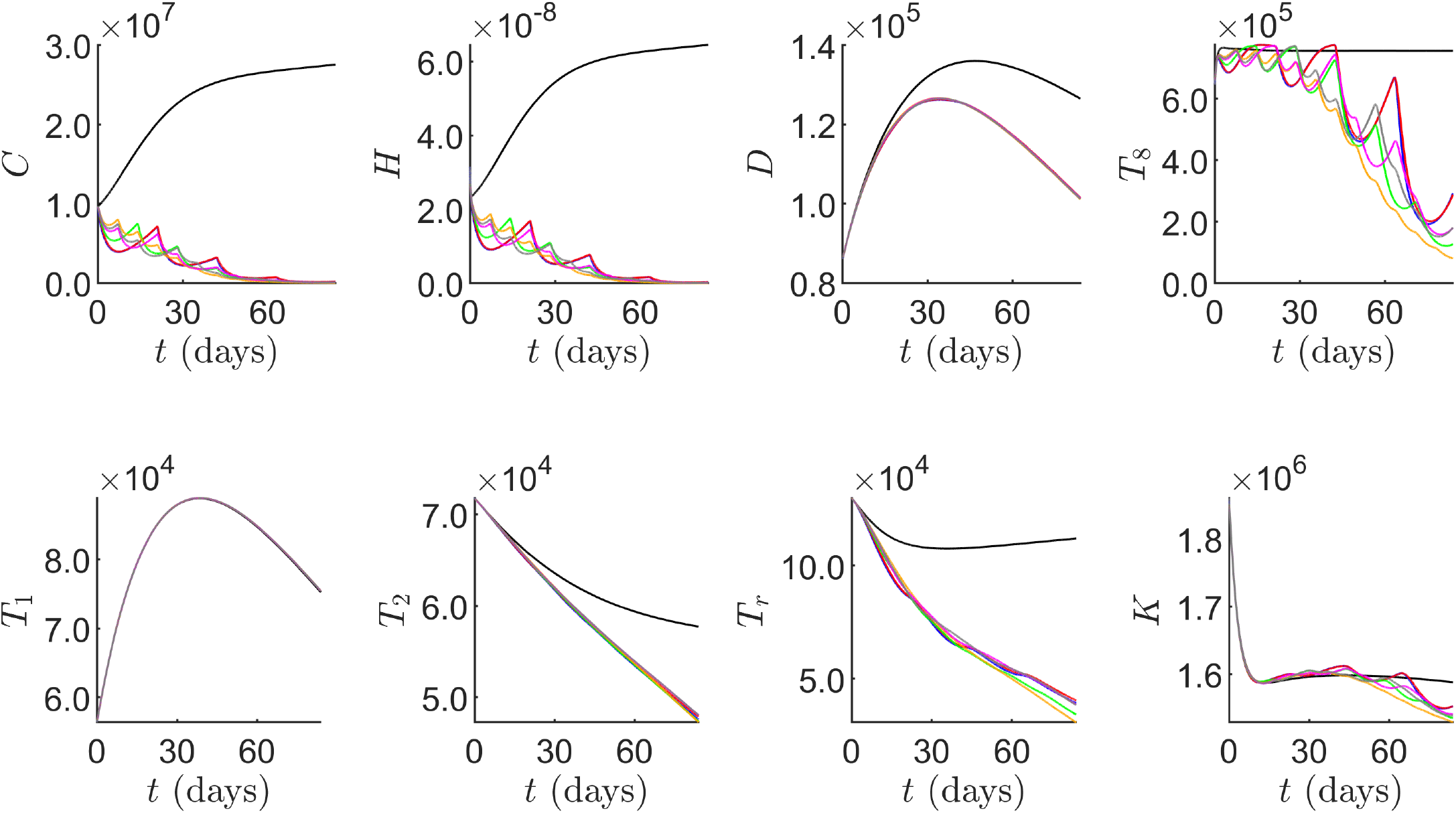
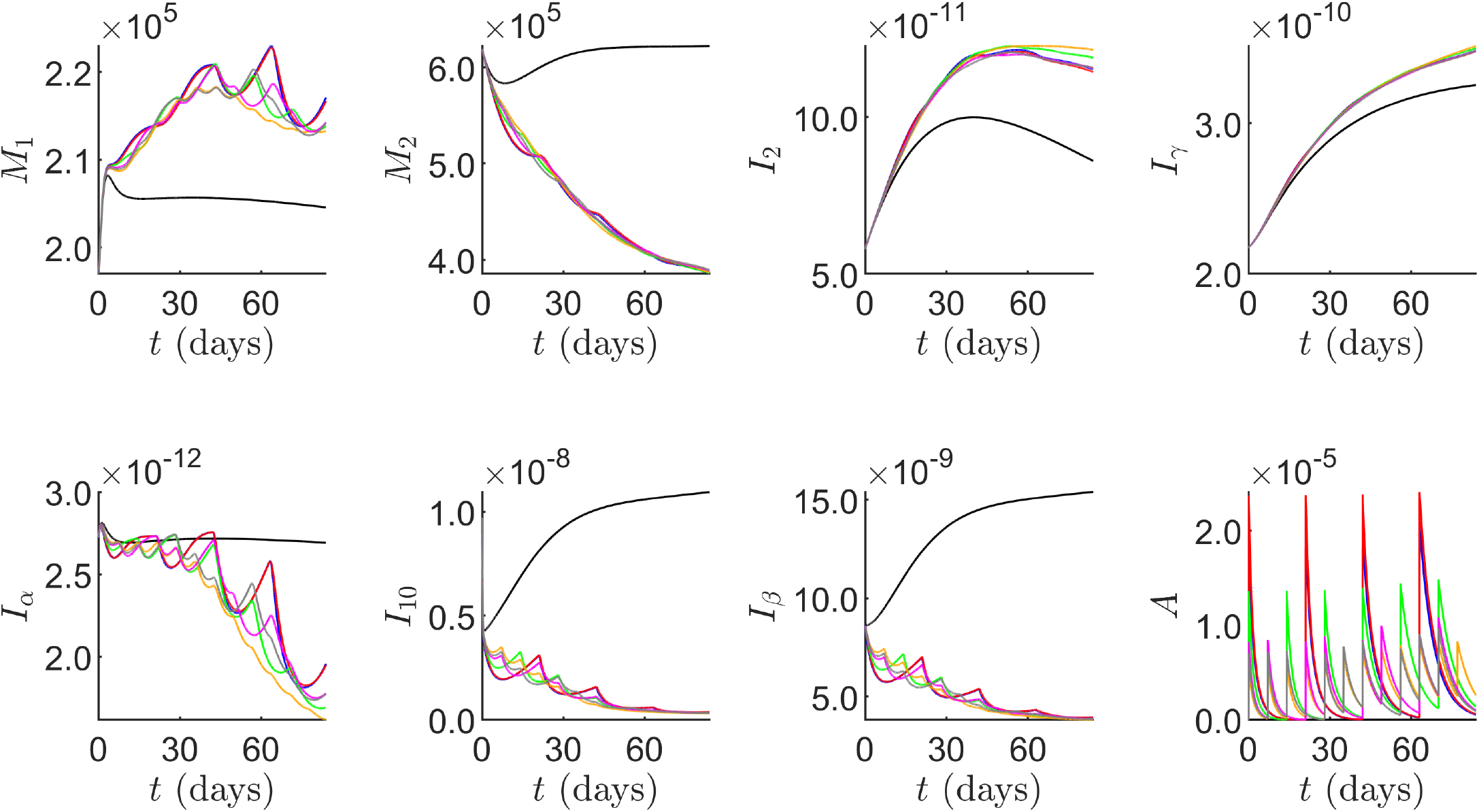
Time traces of notable variables in the model, with the units of the variables as in Table 1. Time traces with no treatment in black, SR in blue, optimal triweekly regimen in red, optimal biweekly regimen in green, optimal weekly regimen in orange, optimal 21-day cycle in magenta and optimal 28-day cycle in grey.

**Figure 5:**
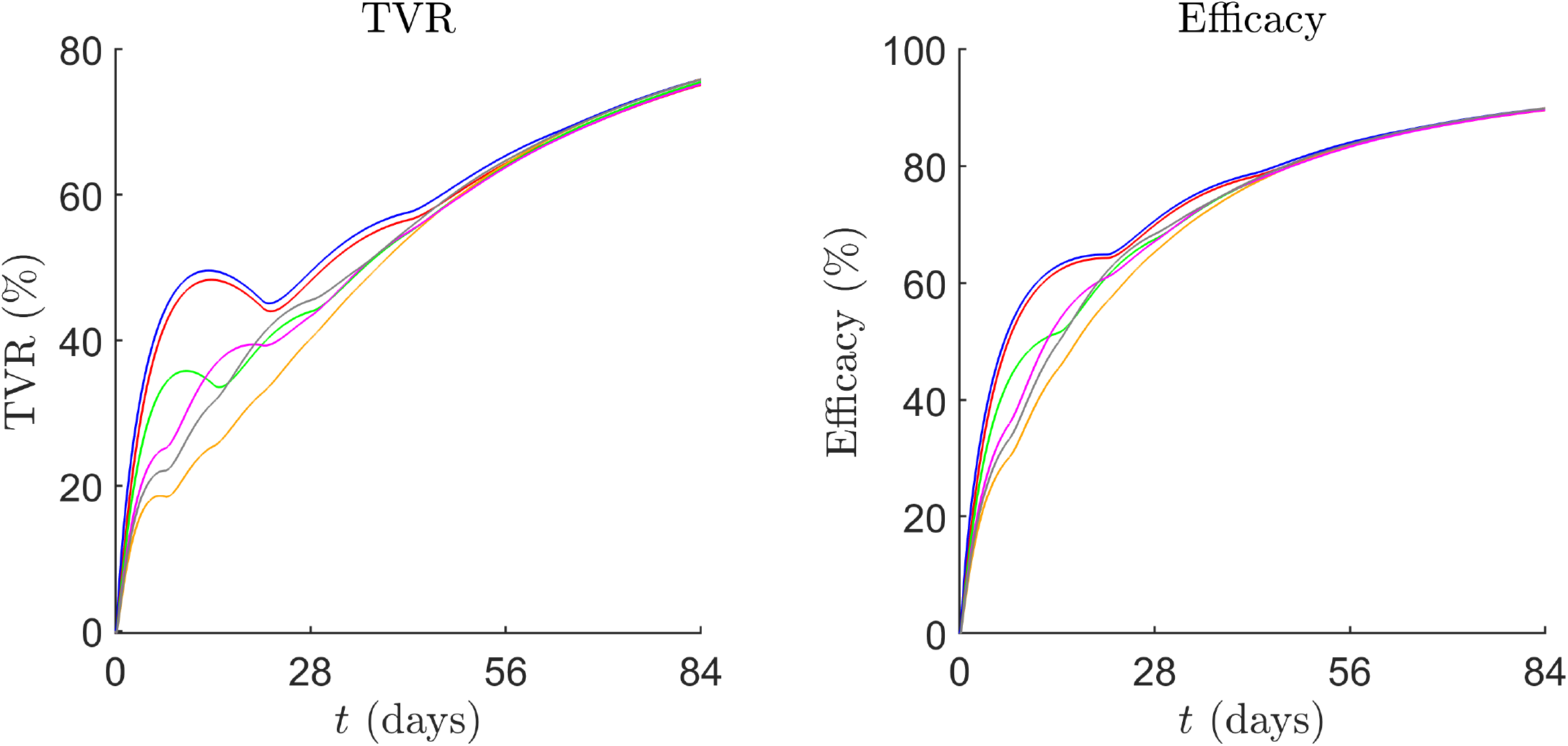
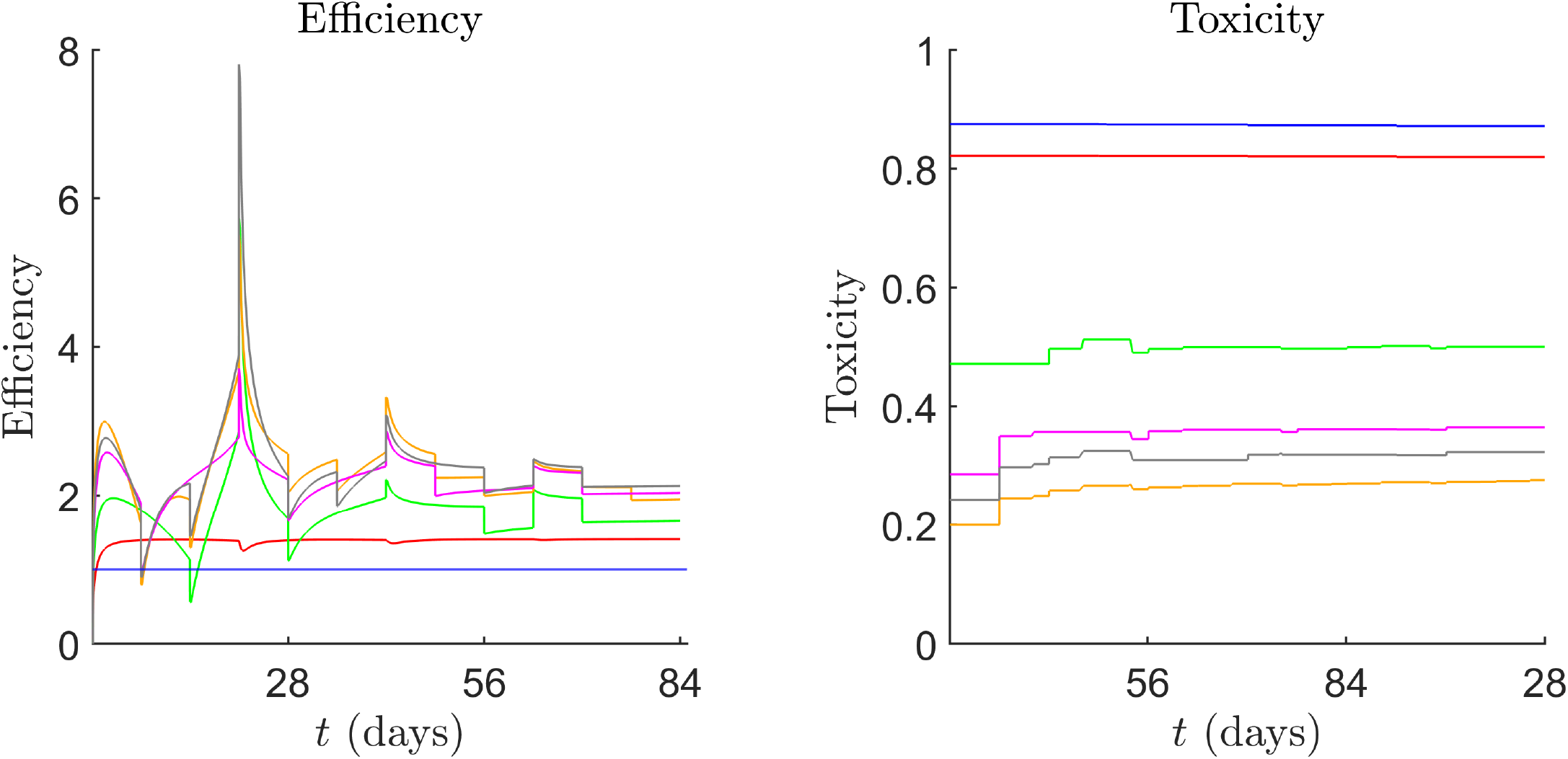
Time traces of TVR, efficacy, efficiency and toxicity. SR in blue, optimal triweekly in red, optimal biweekly in green, optimal weekly in orange, optimal 21-day cycle in magenta and optimal 28-day cycle in grey.

**Figure 6:**
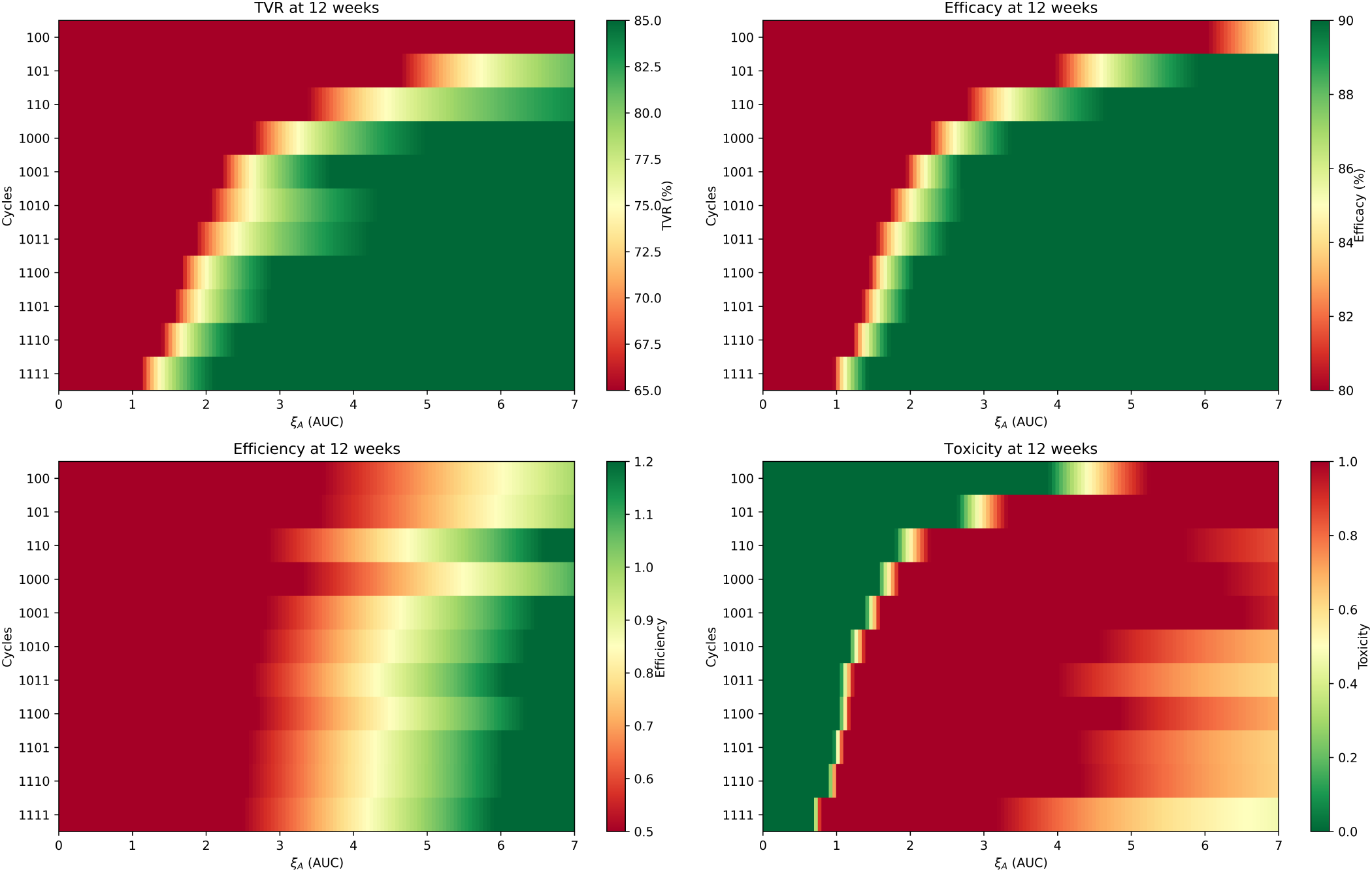
Heatmap of 21-day and 28-day cycles. The binary encodings on the y-axis correspond to infusion days 1, 8 and 15 for the 21-day cycles, and 1, 8, 15 and 21 for the 28-day cycles.

It is also difficult to compare our results to pre-existing clinical trials for first-line neoadjuvant HGSOC due to the lack of time-series data. However, we run the chemotherapy regimens from the clinical trials in Table 7 on our model and compare them with the SR and optimal weekly solutions in Table 8. We also display their TVR, efficacy, efficiency and toxicity in Figure 7.

**Table 7:** Neoadjuvant clinical trials with varying dosing regimens. Paclitaxel dose (*P* dose) is given in (mg*/*m^2^), and carboplatin dose (*A* dose) is given in AUC. *Dose given on Day 1, 8, 15.

| Trial | $P$ Dose | Freq. | $A$ Dose | Freq. | Reference |
| --- | --- | --- | --- | --- | --- |
| MITO-7 | 60 | Weekly | 2 | Weekly | [160] |
| S B Dessai et. al/ICON8 | 80 | Weekly | 2 | Weekly | [161, 162] |
| T. Safra et. al* | 80 | 28-day cycle | 2 | 28-day cycle | [161] |
| ICON8 | 80 | Weekly | 5 | Triweekly | [162] |
| GOG-0262/JGOG3016 | 80 | Weekly | 6 | Triweekly | [163, 164] |

**Table 8:** Comparison of TVR, efficacy, efficiency and toxicity for various chemotherapy regimens.

| Regimen | TVR (%) | Efficacy (%) | Efficiency | Toxicity |
| --- | --- | --- | --- | --- |
| Standard Regimen | 75.85 | 89.86 | 1.00 | 0.87 |
| Optimal Weekly | 75.83 | 89.86 | 1.92 | 0.28 |
| MITO-7 | 86.36 | 94.27 | 0.87 | 0.41 |
| S B Dessai et. al/ICON8 | 86.63 | 94.39 | 0.79 | 0.42 |
| T. Safra et. al | 84.59 | 93.53 | 0.79 | 0.41 |
| ICON8 | 79.01 | 91.19 | 0.41 | 0.71 |
| GOG-0262/JGOG3016 | 83.08 | 93.90 | 0.36 | 0.86 |

**Figure 7:**
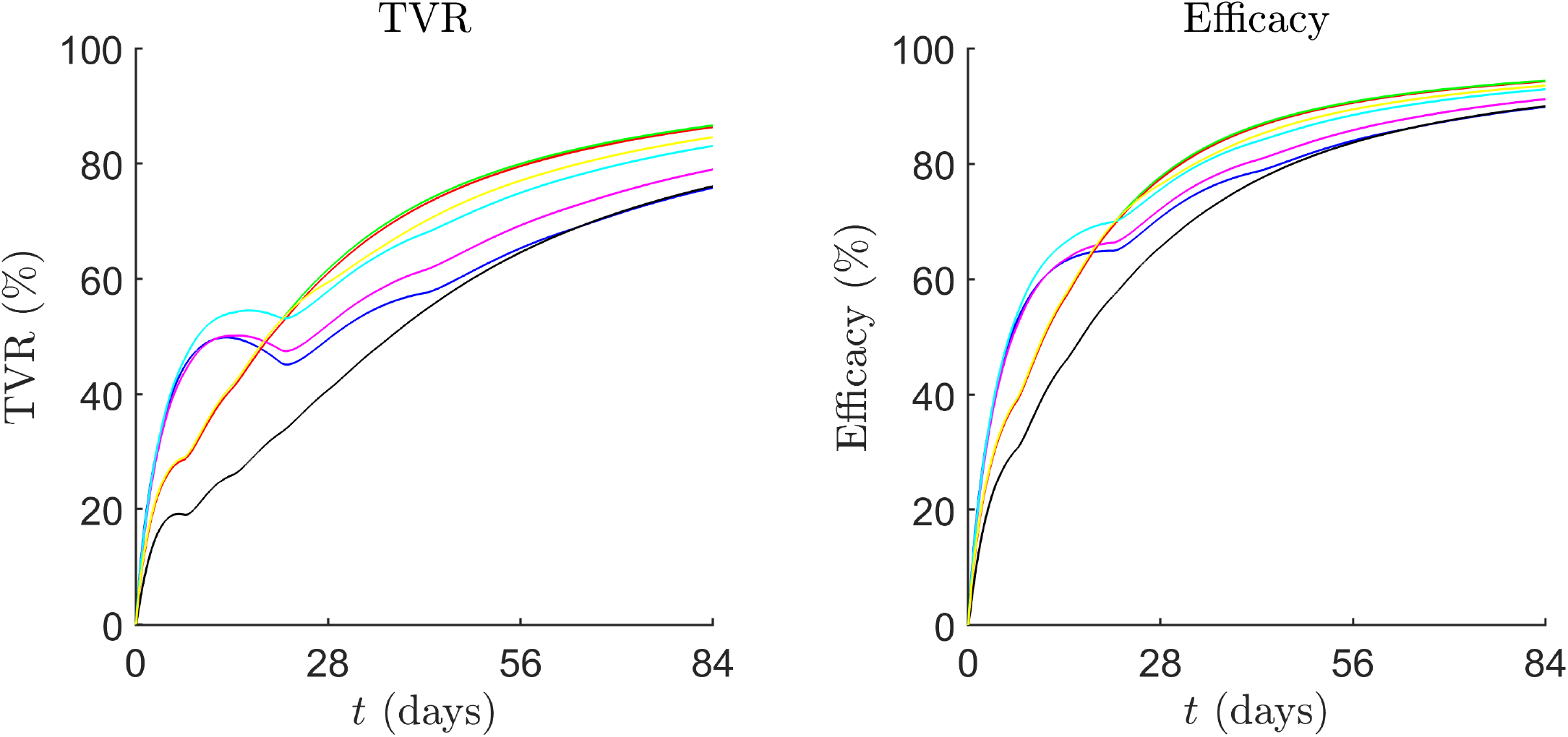
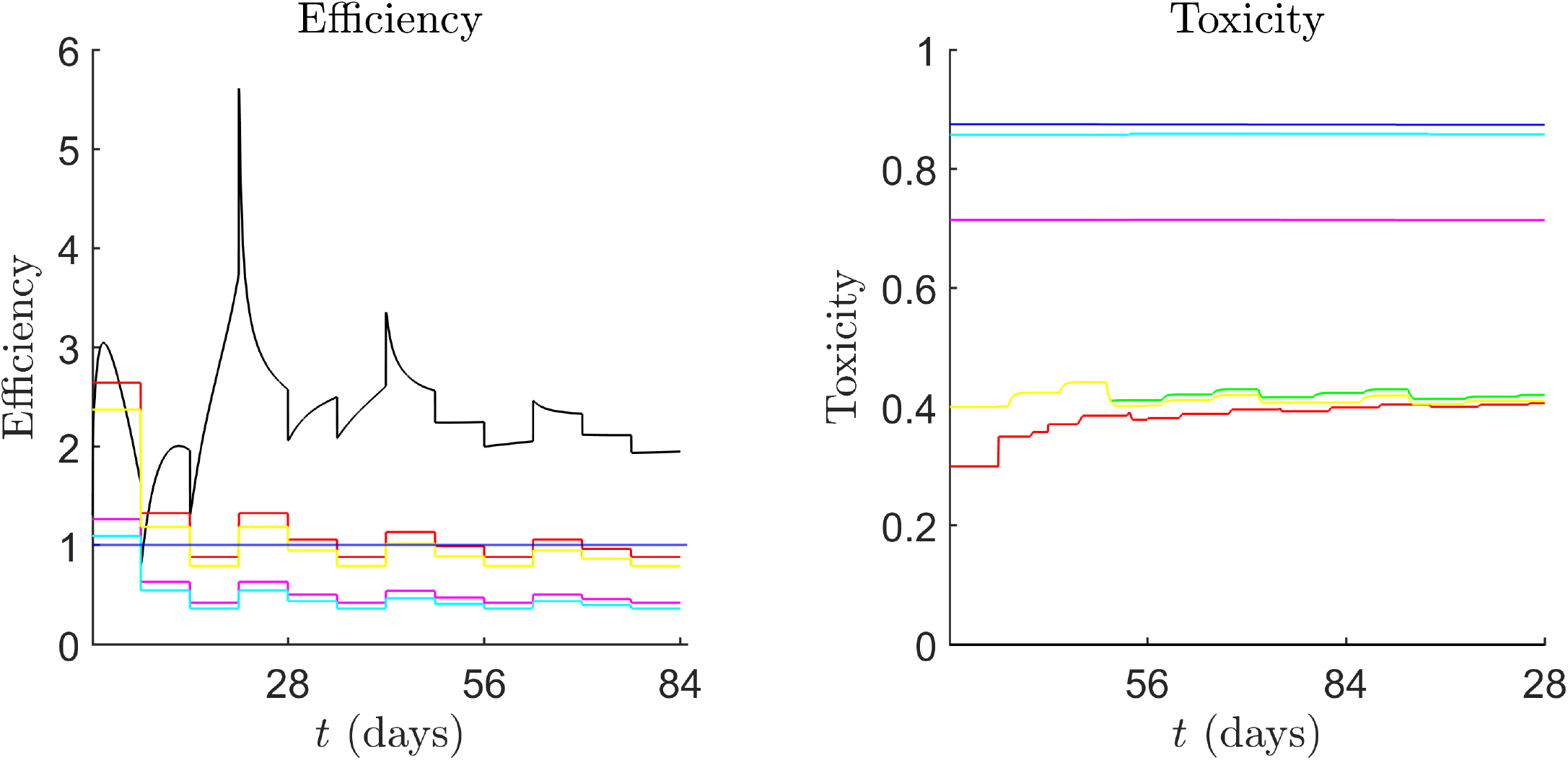
Time traces of TVR, efficacy, efficiency and toxicity. SR in blue, MITO-7 in red, S B Dessai et. al/ICON8 in green, T. Safra et. al in yellow, ICON8 trial in magenta, GOG-0262/JGOG3016 in cyan and the optimal weekly solution in black.

## 5 Discussion

To explain the behaviour of the treatment regimens, we first analyse the immune trajectories from Figure 4 to understand the TME in the absence of treatment. At first, we analyse the effects of paclitaxel and carboplatin treatment, by only considering the SR. One of the main drivers of the immune response against cancer is TNF and IFN-*γ* induced cancer cell necrosis, which triggers the release of DAMPs and the maturation of immature DCs. Elevated levels of TGF-*β* and IL-10 inhibit this maturation, causing decreased activation of CD8+ and Th1 cells in the TDLN which migrate to the TS and sustain the cytotoxic response. We observe on average elevated levels of TGF-*β* and IL-10 at 1.24 ×10^*−*6^ g*/*cm^3^ and 7.70 ×10^*−*7^ g*/*cm^3^, respectively, for no treatment, compared to 4.79 ×10^*−*7^ g*/*cm^3^ and 1.07 ×10^*−*7^ g*/*cm^3^ with treatment. As TGF-*β* promotes the proliferation of Tregs, we observe a relatively increased Treg population of 9.34 ×10^6^ cell*/*cm^3^ without treatment compared to 5.95 ×10^6^ cell*/*cm^3^ with treatment, which inhibits the cytotoxic activity of CD8+ T cells and Th1 cells at the TS. Tregs also secrete additional TGF-*β* and IL-10, promoting an immunosuppressive positive feed-forward loop. Additionally, as IL-10 and TGF-*β* drive macrophage polarisation toward the anti-inflammatory M2 phenotype, this establishes another immunosuppressive feed-forward loop as M2 macrophages secrete IL-10 and TGF-*β*. Hence, we observe an increased average M1/M2 ratio of 0.40 for the SR compared to 0.28 without treatment. This reinforces an environment where cytotoxic immune cells are functionally impaired and outnumbered by immunosuppressive components, allowing cancer to proliferate and overwhelm the immune system.

During chemotherapy treatment, the reduction in cancer concentration causes a large decrease in TGF-*β* and IL-10, which decreases the polarisation of macrophages to the M2 phenotype and the expansion of Tregs. As Tregs inhibit CD8+ and Th1 secretion of IL-2, a decreased Treg population reduces this inhibition, promoting a positive feedback loop, as IL-2 is a growth factor for CD8+ and Th1 cells.

At the same time, a reduced effector Treg population decreases the inhibition of IFN-*γ* production by CD8+ T cells and NK cells, which, in combination with increased IL-2 and decreased TGF-*β* and IL-10, leads to an increase in the polarisation and repolarisation of macrophages to the pro-inflammatory M1 phenotype. These macrophages further secrete pro-inflammatory cytokines, including IL-12, TNF and IFN-*γ*. However, the TNF concentration decreases to 2.04 ×10^*−*10^ g*/*cm^3^ with treatment at 12 weeks compared to 2.28 ×10^*−*10^ g*/*cm^3^ without treatment. This is due to a reduction in the DAMP concentration and the subsequent maturation of DCs, which leads to a decreased activation of T cells in the TDLN, and hence T cell populations at the TS relative to those without treatment. Additionally, chemotherapy, which targets proliferating cells such as T cells, further contributes to the decrease, leading to reduced secretion rates of cytokines such as TNF and IFN-*γ* as we assumed a constant influx of naïve immune cells over time. Hence, we observe that chemotherapy disrupts immunosuppressive feedback loops in the TME, allowing the immune system to shift towards a pro-inflammatory response.

Within the chemotherapy regimens, we find that lower dosage, more frequent carboplatin is more efficient and comparably as effective than higher dosages with lower frequencies (Table 6). In the absence of carboplatin resistance, we expect that optimal regimens feature carboplatin monotherapy, as carboplatin has a prolonged duration of cytotoxic activity than paclitaxel, despite having a shorter plasma half-life (1–2 hours vs 9.9 hours) [152, 153]. This is due to the behaviour of the platinum moiety, which, after hydrolysis, binds irreversibly to plasma proteins and is eliminated over several days [165]. As carboplatin is more cytotoxic per milligram than paclitaxel, the most efficient regimens that are both non-toxic and achieve an acceptable TVR do not include paclitaxel. High-frequency dosing of carboplatin results in decreased oscillation in carboplatin concentrations so that cancer cells are subjected to continuous cytotoxicity, inhibiting cancer growth between infusions, as observed in Figure 3 for the optimal triweekly regimen and SR. As shown in Figure 4, the optimal biweekly, weekly, 21-day, and 28-day cyclic regimens consistently sustain a non-zero carboplatin concentration, with each subsequent administration resulting in an increased sustained carboplatin concentration, whereas the corresponding concentrations for the optimal triweekly regimen and SR rapidly approach zero after one week. As the peak concentrations reduce even with lower dosing, these solutions exhibit a lower toxicity, with an average of 0.37 compared to the optimal triweekly regimen and SR with an average of 0.84.

Analysing the TVR can provide further insights into these results. The SR, despite its high toxicity, leads to the largest increase in TVR in the first two weeks. However, due to the longer rest periods required to tolerate this treatment, the TVR decreases due to cancer growth in the absence of sustained cytotoxicity from chemotherapy between cycles (Figure 5). The rates of growth also decrease as treatment progresses due to a lower cancer concentration and shift towards a more pro-inflammatory immune state in which cytotoxic cells and cytokines promote cancer lysis in between cycles. Although the TVR of the biweekly, weekly, 21-day and 28-day cyclic regimens have a less rapid increase in the early stages of treatment, they gradually increase and exhibit a smaller degree of growth between cycles due to more frequent infusions. We also observe that all optimal regimens are more efficient than the SR at all time points and vary according to infusion spacing and dosing. On the first day, the efficiency is inversely proportional to the dosages administered, with lower-dosage regimens displaying the highest efficiency due to similar TVRs between the regimens at the initial infusion. On days in which the SR regimen does not inject but the biweekly, weekly, or cyclic regimens do, we observe a sharp decrease in efficiency, followed by a gradual increase in efficiency due to the delay in the effects of chemotherapy in the system. For days on which all regimens include a dose coinciding with that of the SR infusion, we observe a spike in efficiency as the SR has a higher comparative drug concentration, with the magnitude of this spike decreasing as the drug concentrations over time become more similar. In combination with the similarity of the TVR time trace between regimens, this also explains the reduction in oscillations as the treatment approaches 84 days since commencement.

Additionally, 21-day and 28-day cyclic regimens can serve as effective alternative dosing strategies comparable to those with constant spacing, with the inclusion of rest weeks allowing patients to recover from side effects, potentially improving treatment adherence. From Table 6, we observe that rest weeks are compensated by higher carboplatin doses compared to regimens administered weekly. We also observe in Figure 6 that within the 21-day and 28-day cyclic treatments, regimens that administer infusions in the first two weeks can sustain lower overall carboplatin dosages while maintaining comparable TVR, efficacy, and toxicity to the optimal biweekly and weekly regimens. Notably, dosing on days 1 and 8 in the 28-day cycle appears sufficient to sustain a one to two-week break, as indicated by the similar heatmap profiles of the 28-day cycles 1101 and 1100. However, when a break is introduced on day 8 (as in cycles 1011, 1010, 1001, and 1000), higher carboplatin doses are required to achieve a comparable TVR, efficacy, and toxicity profile to regimens that include dosing on day 8. This is due to the lack of sustained cytotoxic response in the early stages of treatment, allowing the still sizeable tumour to recover before the immune response becomes sufficiently pro-inflammatory. We also note that for all 21-day and 28-day cyclic regimens, except for the 21-day cycles 100 and 101 and the 28-day cycle 1000, which omit dosing on day 8, the efficacy profiles remain similar. Although the 28-day cycles 1011, 1010, and 1001 also omit dosing on day 8, they have a lower ratio of rest to non-rest weeks, suggesting that longer rest periods combined with later administration leads to an increase in tumour concentration, necessitating higher drug doses to compensate. This, in turn, reduces overall treatment efficiency, and as a result, the optimal 21-day and 28-day cyclic regimens administer infusions in the first two and three weeks, respectively.

Heatmaps of the triweekly, biweekly and weekly regimens, shown in Figure B.1, Figure B.2 and Figure B.3, respectively, compare the TVR, efficacy, efficiency and toxicity of treatments consisting of varying carboplatin and paclitaxel doses. Of note, is that the region of acceptable TVR shifts according to the infusion frequency of carboplatin, with higher frequencies enabling lower doses of carboplatin to sustain TVRs similar to those of lower frequencies. Compared to the SR, we observe in Figure B.1, that high TVRs for triweekly regimens only occur above 5.5 AUC, and so all effective triweekly regimens will exhibit a high toxicity. Efficiency also primarily depends on the frequency and dosage of administered carboplatin, with more frequent, low-dose carboplatin leading to higher efficiency. However, for the biweekly and weekly regimens, Figure B.2 and Figure B.3 demonstrate that the efficiency of treatments with paclitaxel doses below 80 mg*/*m^2^ increases if sufficient carboplatin is administered. Interestingly, clinical trials comparing weekly paclitaxel (60 and 80 mg*/*m^2^) to weekly carboplatin at 2 AUC have reported PFS comparable to the SR, but with reduced patient toxicity [29, 166, 167]. We observe in Table 8 and Figure 7 that the addition of paclitaxel increases the TVR compared to the optimal weekly solution, from 75.83% to 86.36% and 86.63% for 60 mg*/*m^2^ and 80 mg*/*m^2^, respectively, at the expense of increased toxicity. However, Figure B.3 indicates that dosages of paclitaxel beyond 85 mg*/*m^2^ have a poorer efficiency and higher toxicity, and so, the optimal regimens will have a paclitaxel dose less than 85 mg*/*m^2^. Due to the increased efficacy of carboplatin per milligram, compared to paclitaxel, the optimal treatments, which balance TVR, efficiency and toxicity, consist solely of carboplatin. In particular, these results demonstrate diminishing increases in TVR as chemotherapy dosing increases, which are accompanied by disproportionate increases in toxicity.

It should be noted that the model has several limitations, with the potential for addressing these issues serving as exciting avenues for future research.

- Although most OC patients exhibit an initial response to first-line chemotherapy and the inclusion of platinum agent is a dominant factor in the survival of ovarian cancer, in practice, combination chemotherapy with independent mechanisms of action is preferred to minimise the evolution of platinum resistance [5, 168–170]. Future work involves extending this model to incorporate resistance and optimise regimens to maintain effective cancer reduction whilst minimising the effects of acquired resistance.
- Our model does not account for spatial effects, which could provide insights into the distribution and clustering of immune cell populations within the TME and how these spatial dynamics influence tumour progression and treatment outcomes [171, 172].
- We considered only the M1/M2 macrophage dichotomy; however, their inherent plasticity suggests that their phenotypes are better represented as a continuum, allowing them to adopt mixed functional states that encompass both pro-inflammatory M1 and anti-inflammatory M2 characteristics [173].
- We did not account for the role of cytokines in the TDLN, which can influence T cell activation and proliferation and play a significant role in effector T cell differentiation [174, 175].

In this work, we have provided a framework for mathematically modelling the immunobiology of HGSOC using delay-differential equations. We analysed the optimised paclitaxel and carboplatin therapies for tumour volume reduction, efficacy, efficiency and toxicity for first-line chemotherapy treatment, and we conclude that low-dose high-frequency carboplatin monotherapy provides comparable results to the standard regimen. Additionally, 21-day and 28-day cyclic regimens can serve as alternative dosing strategies with rest weeks, allowing patients to recover from the toxic side effects of chemotherapy. Future work will focus on extending this model to incorporate carboplatin resistance, a major challenge in HGSOC, and optimising treatment to minimise its impact while maintaining effective tumour reduction.

## 6 CRediT authorship contribution statement

**Cristina Koprinski**: conceptualisation, data curation, formal analysis, funding acquisition, investigation, methodology, project administration, resources, software, validation, visualisation, writing — original draft, writing — review & editing.

**Georgio Hawi**: conceptualisation, data curation, formal analysis, funding acquisition, investigation, methodology, project administration, resources, software, validation, visualisation, writing — original draft, writing — review & editing.

**Peter S. Kim**: conceptualisation, formal analysis, investigation, methodology, project administration, resources, supervision, validation, visualisation, writing — original draft, writing — review & editing.

Cristina Koprinski and Georgio Hawi contributed equally to this work.

## 8 Data availability

All data and procedures are available within the manuscript and its Supporting Information file.

## 9 Acknowledgements

This work was supported by an Australian Academy of Technological Sciences and Engineering Elevate Scholarship and an Australian Government Research Training Program Scholarship. PSK gratefully acknowledges support from the Australian Research Council Discovery Project (DP230100485).

