## Supplementary Information for "Optimising Chemotherapy for Advanced High-Grade Serous Ovarian Cancer via Delay-Differential Equations"

### A Parameter Estimation

We estimate all parameters, where possible, under the assumption that no chemotherapy has/will be administered. The exception to this is the parameters directly related to chemotherapy treatment, for which the assumptions are explicitly stated during estimation. We also denote 1 d to be 1 day, 1 h to be 1 hour, 1 m to be 1 minute, and 1 s to be 1 second. Many of the assumptions and techniques in this section are adopted from [1] and [2].

#### A.1 Cell Initial Conditions and Steady states

##### A.1.1 Tumour Site

Digital cytometry can be used to analyse cell populations in ovarian cancer without physical cell isolation [3, 4]. This is achieved by analysing bulk tissue transcriptomic data from individual patients, followed by deconvolution [5, 6].

We used the UCSC Xena web portal [7] to obtain RSEM normalised RNA-sequencing (RNA-seq) gene expression profiles of tumours from the TCGA OV database. The corresponding clinical and biospecimen data were downloaded from the GDC portal [8] which includes tumour dimensions, necrotic cell percentage, AJCC TNM stage and ovarian cancer subtype. The samples were filtered from primary tumours of patients with AJCC stage III or stage IV high-grade serous adenocarcinoma, with complete necrosis percentage data. We used the stage IIIA, IIIB and IIIC samples as part of our estimation for the initial conditions, and the stage IV samples for the steady state conditions. For all digital cytometry algorithms below, we take the median of the cell population estimates and then normalise them row-wise such that they sum to 1.

To estimate immune cell populations in HGSOC, we applied the ImmuCellAI [9] and CIBERSORT [6] algorithms, and combined their results to obtain estimates for all cell types in the model.

First, we used the ImmuCellAI algorithm to estimate the abundance of 24 immune cells with their data dictionary shown in Table A.1. The correspondence of these cells to the steady state variables is shown in Table A.2.

Table A.1: ImmuCellAI data dictionary.

| **Cell in ImmuCellAI** | **Description** |
|---|---|
| DC | Dendritic cells |
| Bcell | B cells |
| Monocyte | Monocyte cells |
| Macrophage | Macrophage cells |
| NK | Natural killer cells |
| Neutrophil | Neutrophil cells |
| CD4_T | Other CD4+ T cells |
| CD8_T | Other CD8+ T cells |
| NKT | Natural Killer T cells |
| Gamma_delta | $\gamma\delta$ T cells |
| CD4_naïve | Naïve CD4+ cells |
| Tr1 | Type 1 regulatory T cells |
| nTreg | Natural Tregs |
| iTreg | immature Tregs |
| Th1 | Th1 cells |
| Th2 | Th2 cells |
| Th17 | Th17 cells |
| Tfh | T follicular helper cells |
| CD8_naïve | Naïve CD8+ cells |
| Cytotoxic | CD8+ T cells |
| Exhausted | Exhausted T cells |
| MAIT | Mucosal-associated invariant T cells |

Table A.2: Mapping of state variables to ImmuCellAI cell types.

| **State Variable** | **ImmuCellAI Cell Type** |
|---|---|
| $D_0, D$ | DC |
| $T_8$ | Cytotoxic |
| $T_1$ | Th1 |
| $T_2$ | Th2 |
| $T_r$ | nTreg |
| $M_0, M_1, M_2$ | Macrophage |
| $K_0, K$ | NK |

The final aggregates from the ImmuCellAI algorithm are shown in Table A.3.

Table A.3: ImmuCellAI initial-condition (stage III) and steady-state (stage IV) cell proportions.

| **Cell** | **Initial Condition: Stage III** | **Steady State: Stage IV** |
|---|---|---|
| DC | 0.09462935 | 0.09638556 |
| Bcell | 0.07540612 | 0.07695609 |
| Monocyte | 0.09936205 | 0.09407111 |
| Macrophage | 0.08447555 | 0.09145037 |
| NK | 0.1322621 | 0.1314804 |
| Neutrophil | 0.08717281 | 0.07971137 |
| CD4_T | 0.07560978 | 0.07043560 |
| CD8_T | 0.1076817 | 0.1083324 |
| NKT | 0.08173929 | 0.09083879 |
| Gamma_delta | 0.09660578 | 0.09999272 |
| CD4_naïve | 0.0009590279 | 0.0009289585 |
| Tr1 | 0.007553911 | 0.006579816 |
| nTreg | 0.008554112 | 0.008248465 |
| iTreg | 0.004716786 | 0.004612348 |
| Th1 | 0.003714472 | 0.003637384 |
| Th2 | 0.004730036 | 0.003742143 |
| Th17 | 0.005708111 | 0.005560705 |
| Tfh | 0.006554541 | 0.005555508 |
| CD8_naïve | 0.003801827 | 0.003731572 |
| Cytotoxic | 0.005670733 | 0.005661327 |
| Exhausted | 0.007463236 | 0.006623807 |
| MAIT | 0.005628611 | 0.005463597 |

Next, we used the CIBERSORT algorithm to determine the proportions of $D_0$ and $D$, $M_0$, $M_1$, and $M_2$, and $K_0$ and $K$. CIBERSORT determines the immune cell proportions of 22 cells using the LM22 signature matrix, with the mappings between the signature matrix and steady state variables shown in Table A.4.

Table A.4: Mapping of state variables to LM22 key.

| **State Variable** | **LM22 Key** |
|---|---|
| $D_0$ | Dendritic cells resting |
| $D$ | Dendritic cells activated |
| $T_r$ | Tregs |
| $M_0$ | Macrophages M0 |
| $M_1$ | Macrophages M1 |
| $M_2$ | Macrophages M2 |
| $K_0$ | NK cells resting |
| $K$ | NK cells activated |

The final aggregates from the CIBERSORT algorithm are shown in Table A.5.

Table A.5: Immune cell proportions for the initial conditions (stage III) and steady states (stage IV) from CIBERSORT.

| **Immune Cells** | **Initial Condition: Stage III** | **Steady State: Stage IV** |
|---|---|---|
| B cells naïve | 0.03050403 | 0.01874005 |
| B cells memory | 0.01886837 | 0.01277149 |
| Plasma cells | 0.005974082 | 0.006034249 |
| T cells CD8 | 0.12853162 | 0.09345462 |
| T cells CD4 naïve | 0.000767577 | 0.000000000 |
| T cells CD4 memory resting | 0.09285640 | 0.09489761 |
| T cells CD4 memory activated | 0.02519437 | 0.04775849 |
| T cells follicular helper | 0.01352237 | 0.01264098 |
| T cells regulatory Tregs | 0.03295735 | 0.02727298 |
| T cells gamma delta | 0.003569945 | 0.005942883 |
| NK cells resting | 0.009804888 | 0.032144810 |
| NK cells activated | 0.1240267 | 0.1107462 |
| Monocytes | 0.03921748 | 0.04328349 |
| Macrophages M0 | 0.09458732 | 0.07669412 |
| Macrophages M1 | 0.05313177 | 0.04207369 |
| Macrophages M2 | 0.1671248 | 0.1488886 |
| Dendritic cells resting | 0.01289833 | 0.01175511 |
| Dendritic cells activated | 0.01261288 | 0.01540667 |
| Mast cells resting | 0.03220151 | 0.11708720 |
| Mast cells activated | 0.09394795 | 0.07613049 |
| Eosinophils | 0.002073223 | 0.002664525 |
| Neutrophils | 0.005627045 | 0.003611703 |

To estimate the proportions of subtypes within DCs, NK cells and macrophages in ImmuCellAI, we used the relative proportions of these subtypes derived from CIBERSORT. For example, for the initial conditions (stage III), the ratio of resting NK cells to total NK cells in CIBERSORT (Table A.5) is $\frac{0.009804888}{0.009804888+0.1240267} = 0.07326$. Thus, the proportion of resting NK cells in ImmuCellAI (Table A.3) is $0.07326 \times 0.1322621 = 0.0096899$. We repeat this for $D_0$, $D$, $M_0$, $M_1$, $M_2$, $K_0$ and $K$ for stage III and stage IV, with the results shown in Table A.6,

Table A.6: DC, NK cell and macrophage initial-condition (stage III) and steady-state (stage IV) estimates for ImmuCellAI from CIBERSORT.

| **Stage** | **Initial Condition: Stage III** | **Steady State: Stage IV** |
|---|---|---|
| NK cells resting | 0.0096899 | 0.02957788 |
| NK cells activated | 0.12257219 | 0.10190252 |
| Dendritic cells resting | 0.04784409 | 0.04171388 |
| Dendritic cells activated | 0.04678526 | 0.05467168 |
| Macrophages M0 | 0.02537866 | 0.02620413 |
| Macrophages M1 | 0.01425576 | 0.01437535 |
| Macrophages M2 | 0.04484113 | 0.05087089 |

Combining these results with ImmucellAI gives us Table A.7 for our initial conditions and steady-state values.

Table A.7: ImmuCellAI and CIBERSORT initial-condition (stage III) and steady-state (stage IV)cell proportions.

| Cell | Initial Condition: Stage III | Steady state: Stage IV |
|---|---|---|
| $D_0$ | 0.04784409 | 0.04171388 |
| $D$ | 0.04678526 | 0.05467168 |
| $T_0^8$ | 0.003801827 | 0.003731572 |
| $T_8$ | 0.005670733 | 0.005661327 |
| $T_0^4$ | 0.0009590279 | 0.0009289585 |
| $T_1$ | 0.003714472 | 0.003637384 |
| $T_2$ | 0.004730036 | 0.003742143 |
| $T_r$ | 0.008554112 | 0.008248465 |
| $K_0$ | 0.0096899 | 0.02957788 |
| $K$ | 0.12257219 | 0.10190252 |
| $M_0$ | 0.02537866 | 0.02620413 |
| $M_1$ | 0.01425576 | 0.01437535 |
| $M_2$ | 0.04484113 | 0.05087089 |

From [10], we have that the density of immune cells in healthy ovaries is approximately $1.34 \times 10^7$ cell/g. Assuming a tissue density of 1.03 g/mL results in a total immune cell density of $1.38 \times 10^7$ cell/cm$^3$. We also note the presence of lymphadenopathy in advanced cancer [11], which increases the total number of lymphocytes to at most 10% [10]. As such, to estimate the total density of immune cells in the TS in advanced HGSOC, we multiplied the proportions of lymphocytes (Bcell, NK, CD4, CD8, NKT, Gamma_delta, CD4_naïve, Tr1, nTreg, iTreg, Th1, Th2, Th17, Tfh, CD8_naïve, Cytotoxic, Exhausted and MAIT) in Table A.3 by 1.10 to account for lymphadenopathy. We then summed them and multiplied the totals by $1.38 \times 10^7$ cell/cm$^3$ to give us that the total immune cell density at the initial condition and steady state is $1.47 \times 10^7$ cell/cm$^3$. Thus, multiplying the values in Table A.7 by $1.47 \times 10^7$ cell/cm$^3$ gives us the initial and steady-state values in Table A.8.

Table A.8: Initial-condition (stage III) and steady-state (stage IV) cell densities. All units are in cell/cm$^3$.

| Cell | Initial Condition: Stage III | Steady state: Stage IV |
|---|---|---|
| $D_0$ | $6.60 \times 10^5$ | $5.76 \times 10^5$ |
| $D$ | $6.46 \times 10^5$ | $7.55 \times 10^5$ |
| $T_8$ | $8.61 \times 10^4$ | $8.59 \times 10^4$ |
| $T_1$ | $5.64 \times 10^4$ | $5.52 \times 10^4$ |
| $T_2$ | $7.18 \times 10^4$ | $5.68 \times 10^4$ |
| $T_r$ | $1.30 \times 10^5$ | $1.25 \times 10^5$ |
| $K_0$ | $1.47 \times 10^5$ | $4.49 \times 10^5$ |
| $K$ | $1.86 \times 10^6$ | $1.55 \times 10^6$ |
| $M_0$ | $3.50 \times 10^5$ | $3.62 \times 10^5$ |
| $M_1$ | $1.97 \times 10^5$ | $1.98 \times 10^5$ |
| $M_2$ | $6.19 \times 10^5$ | $7.02 \times 10^5$ |

From the TCGA biospecimen data, the median necrotic cancer cell percentage for stage IV HGSOC is 1%. Denoting the total immune cell density as TIC and $N_p$ as the necrotic cell percentage, we have that at steady state

$$\overline{N} = N_p(\overline{C} + \overline{N}) \implies \frac{\overline{N}}{N_p} = \frac{\overline{C}}{1 - N_p}, \tag{A.1}$$

where $\overline{X}$ denotes the steady-state value of $X$.
Similarly to [12], we assumed at steady state that the density of cancer cells is double the density of total immune cells, i.e., $\overline{C} + \overline{N} = 2 \times \text{TIC}$. Combining this with (A.1) gives us

$$\overline{C} = 2 \times \text{TIC} \times (1 - N_p), \overline{N} = 2 \times \text{TIC} \times N_p. \tag{A.2}$$

This gives us that at steady state, $C \approx 2.91 \times 10^7$ cell/cm$^3$ and $N \approx 2.94 \times 10^5$ cell/cm$^3$.

From [13], it was estimated that the doubling time of ovarian cancer cells is 2.5 months, i.e., 75 days. This matches clinical studies where it was found that the doubling time of ovarian cancer was less than 3 months [14]. Hence, we estimate 2.5 months to be the doubling time. As such, we assume that it takes approximately 4 months, i.e., 120 days, for the cancer concentrations to reach their steady-state values. This gives us the total cancer concentration, $V$, initially equal to $9.70 \times 10^6$ cell/cm$^3$.
From this,

$$C(0) + N(0) = 9.70 \times 10^6 \implies \frac{C(0)}{0.99} = \frac{N(0)}{0.01}.$$

Solving these simultaneously gives us

$$C(0) = 9.60 \times 10^6 \text{ cell/cm}^3,$$
$$N(0) = 9.70 \times 10^4 \text{ cell/cm}^3.$$

We note that, technically, ImmuCellAI is an enrichment-based method that does not provide absolute immune cell proportions but rather estimates abundances across various immune cell subtypes not reported by CIBERSORT. However, normalising these abundances provides a good approximation of the true immune cell proportions, thereby allowing ImmuCellAI to be justifiably employed to estimate immune cell steady states and initial conditions.

##### A.1.2 Tumour-draining Lymph Node

In stage III ovarian cancer, the cancer has spread to the neighbouring lymph nodes and peritoneum. Due to a lack of data in the literature on ovarian cancer T cell proportions in the tumour-draining lymph node (TDLN), we consider proportions in the peritoneum in ovarian cancer patients where there is peritoneum metastasis, as this is one way in which ovarian cancer cells can spread to the lymph nodes [15].

We used the CIBERSORT algorithm to extract the abundance of 22 immune cell types in the peritoneum for stage III and stage IV ovarian serous carcinomas using RNA-seq gene expression profiles from [16]. These results are shown in Table A.9.

Table A.9: CIBERSORT initial-condition (stage III) and steady-state (stage IV) proportions in the TDLN.

| **Immune Cell** | **Initial Condition** | **Steady State** |
|---|---|---|
| B cells naïve | 0.06987580 | 0.06578785 |
| B cells memory | 0.05876439 | 0.04333475 |
| Plasma cells | 0.1156870 | 0.1672947 |
| T cells CD8 | 0.06145052 | 0.05162227 |
| T cells CD4 naïve | 0.02966582 | 0.04585534 |
| T cells CD4 memory resting | 0.1639476 | 0.1067573 |
| T cells CD4 memory activated | 0.08072067 | 0.08735518 |
| T cells follicular helper | 0.06224750 | 0.03779109 |
| T cells regulatory Tregs | 0.08550450 | 0.08927051 |
| T cells gamma delta | 0.005110824 | 0.005414407 |
| NK cells resting | 0.06652821 | 0.05556605 |
| NK cells activated | 0.04985427 | 0.04095482 |
| Monocytes | 0.008356961 | 0.006820465 |
| Macrophages M0 | 0.019148694 | 0.007390066 |
| Macrophages M1 | 0.003045209 | 0.023260345 |
| Macrophages M2 | 0.005362171 | 0.010203703 |
| Dendritic cells resting | 0.009240445 | 0.005610673 |
| Dendritic cells activated | 0.02936432 | 0.08063584 |
| Mast cells resting | 0.007934352 | 0.008037012 |
| Mast cells activated | 0.006437191 | 0.008988437 |
| Eosinophils | 0.01610437 | 0.01133632 |
| Neutrophils | 0.04564918 | 0.04071280 |

CIBERSORT does not provide estimates for naïve CD8+ T cells ($T_0^8$), effector CD8+ T cells ($L_8$), effector Th1 and Th2 CD4+ T cells ($L_1$, $L_2$) and effector Tregs ($L_r$). Due to an absence in the literature of ovarian cancer T cell proportions in the TDLN, we consider data for endometrial cancer to estimate these values, which have overlapping sentinel lymph nodes with ovarian cancer [17]. Additionally, endometrial and ovarian cancer share similar clinical and pathologic features, likely due to a common precursor cell [18, 19].

In order to estimate $T_0^8$, $T_0^4$, $L_8$, $L_1$, $L_2$ and $L_r$ we first considered the proportions of lymphocytes in TDLNs in endometrial cancer as reported in [20]. These proportions are summarised in Table A.10 and Table A.11, and, for simplicity, we considered the early activated CD4+ cells, T regulatory CD4+ cells and most effective T regulatory CD4+ cells categorisations in [20] be Tregs for our model. Similarly, we grouped the early activated CD8+ cells and pre-effector CD8+ cells classifications to be effector CD8+ T cells for our model. However, in [20], the Th1 cell and Th2 cell proportions are reported relative to the total CD4+ population, and we multiply them by the proportion of CD4+ relative to all lymphocytes to get the Th1 and Th2 proportions relative to all lymphocytes.

Table A.10: CD4+ cell proportions in our model relative to lymphocytes in the TDLN for the initial conditions (stage III) and steady states (stage IV) based off proportions reported in [20].

| Cell | Initial Condition | Steady State |
|---|---|---|
| Naïve CD4+ T cell | 0.031 | 0.054 |
| Tregs | 0.26 | 0.363 |
| Th1 | 0.038 | 0.007 |
| Th2 | 0.003 | 0.011 |

Table A.11: CD8+ T cell proportions relative to lymphocytes in the TDLN for the initial conditions (stage III) and steady states (stage IV).

| Cell | Initial Condition | Steady State |
|---|---|---|
| Naïve CD8+ T cell | 0.032 | 0.066 |
| Effector CD8+ T cell | 0.025 | 0.003 |

The density of immune cells in the lymph nodes of an adult is approximately $1.80 \times 10^9$ cell/g, which assuming a tissue density of 1.03 g/cm$^3$, results in a total immune cell density of $1.854 \times 10^9$ cell/cm$^3$ [10]. This gives us the density of immune cells from CIBERSORT to be Table A.12.

Table A.12: CIBERSORT cell densities in the TDLN for the initial conditions (stage III) and steady states (stage IV).

| Immune Cells | Initial Condition | Steady State |
|---|---|---|
| B cells naïve | $1.30 \times 10^8$ | $1.22 \times 10^8$ |
| B cells memory | $1.09 \times 10^8$ | $8.03 \times 10^7$ |
| Plasma cells | $2.14 \times 10^8$ | $3.10 \times 10^8$ |
| T cells CD8 | $1.14 \times 10^8$ | $9.57 \times 10^7$ |
| T cells CD4 naïve | $5.50 \times 10^7$ | $8.50 \times 10^7$ |
| T cells CD4 memory resting | $3.04 \times 10^8$ | $1.98 \times 10^8$ |
| T cells CD4 memory activated | $1.50 \times 10^8$ | $1.62 \times 10^8$ |
| T cells follicular helper | $1.15 \times 10^8$ | $7.00 \times 10^7$ |
| T cells regulatory Tregs | $1.59 \times 10^8$ | $1.66 \times 10^8$ |
| T cells gamma delta | $9.47 \times 10^6$ | $1.00 \times 10^7$ |
| NK cells resting | $1.23 \times 10^8$ | $1.03 \times 10^8$ |
| NK cells activated | $9.24 \times 10^7$ | $7.60 \times 10^7$ |
| Monocytes | $1.55 \times 10^7$ | $1.26 \times 10^7$ |
| Macrophages M0 | $3.55 \times 10^7$ | $1.37 \times 10^7$ |
| Macrophages M1 | $5.64 \times 10^6$ | $4.31 \times 10^7$ |
| Macrophages M2 | $9.94 \times 10^6$ | $1.89 \times 10^7$ |
| Dendritic cells resting | $1.71 \times 10^7$ | $1.04 \times 10^7$ |
| Dendritic cells activated | $5.44 \times 10^7$ | $1.49 \times 10^8$ |
| Mast cells resting | $1.47 \times 10^7$ | $1.49 \times 10^7$ |
| Mast cells activated | $1.19 \times 10^7$ | $1.67 \times 10^7$ |
| Eosinophils | $2.99 \times 10^7$ | $2.10 \times 10^7$ |
| Neutrophils | $8.46 \times 10^7$ | $7.55 \times 10^7$ |

From Table A.9 we considered the ratio of lymphocytes (B cells naïve, B cells memory, T cells CD8, T cells CD4 naïve, T cells CD4 memory resting, T cells CD4 memory activated, T cells follicular helper, T cells regulatory Tregs, T cells gamma delta, NK cells resting and NK cells activated) to all immune cells in the TDLN. This gives us that the proportion of lymphocytes to all immune cells in the TDLN is 0.734 at the initial condition and 0.630 at steady state. Thus, recalling that the total immune cell density in the TDLN is $1.854 \times 10^9$ cell/cm$^3$, we have that the total density of lymphocytes is $1.36 \times 10^9$ cell/cm$^3$ and $1.17 \times 10^9$ cell/cm$^3$ at the initial condition and at steady state, respectively. As such, we can multiply the cell proportions in Table A.10 and Table A.11 by these values, to get estimates for $T_0^4$, $T_0^8$ and $L_8$ at the initial condition and at steady state, as shown in Table A.13.

Table A.13: Naïve CD8+, effector CD8+ and naïve CD4+ densities in the TDLN at the initial condition (stage III) and at steady state (stage IV). All units are in cell/cm$^3$.

| Variable | Initial Condition: Stage III | Steady state: Stage IV |
|---|---|---|
| $T_0^8$ | $4.35 \times 10^7$ | $7.72 \times 10^7$ |
| $L_8$ | $3.40 \times 10^7$ | $3.51 \times 10^6$ |
| $T_0^4$ | $4.22 \times 10^7$ | $6.32 \times 10^7$ |

For the estimates of effector Th1 cells, effector Th2 cells, naïve Tregs and effector Tregs, we must account for the total number of cell divisions that a newly activated T cell undergoes before becoming an effector cell. As justified in Appendix A.7.5, we assume that a naïve CD4+ T cell undergoes $N_1 = 9$ divisions until it becomes effector. Without taking into account degradation during cell division, which is relatively negligible, the number of cells doubles after every division. This gives us that with an activated Th1 cell population, $T_1^{\text{LN}}$, the number of Th1 cells that have completed clonal expansion, i.e., are effector, is,

$$L_1 = \frac{2^{N_1}}{2^{N_1+1} - 1} T_1^{\text{LN}}. \tag{A.3}$$

Table A.10 gives us that $T_1^{\text{LN}} = 5.17 \times 10^7$ cell/cm$^3$ at the initial condition, and that $T_1^{\text{LN}} = 8.19 \times 10^6$ cell/cm$^3$ at steady state. Thus, using (A.3), we get that at the initial condition $L_1 = 2.59 \times 10^7$ cell/cm$^3$ and at steady state, $L_1 = 4.10 \times 10^6$ cell/cm$^3$.

Similarly, we get that

$$L_2 = \frac{2^{N_2}}{2^{N_2+1} - 1} T_2^{\text{LN}}, \tag{A.4}$$

where $N_2 = 9$ as justified in Appendix A.7.5 and $T_2^{\text{LN}}$ is the density of activated Th2 cells in the TDLN.

Table A.10 gives us $T_2^{\text{LN}} = 4.08 \times 10^6$ cell/cm$^3$, and that $T_2^{\text{LN}} = 6.46 \times 10^6$ cell/cm$^3$ at steady state. Thus, using (A.4), we get that at the initial condition $L_1 = 2.04 \times 10^6$ cell/cm$^3$ and at steady state, $L_2 = 4.10 \times 10^6$ cell/cm$^3$.

As we do not have estimates for naïve Tregs, we assume that 10% of all Tregs in the TDLN are naïve. This gives us

$$L_0^r = \frac{T_r^{\text{LN}}}{10}, \tag{A.5}$$

$$L_r = \frac{9}{10} \frac{2^{N_r}}{2^{N_r+1} - 1} T_r^{\text{LN}}, \tag{A.6}$$

where $N_r = 6$ as justified in Appendix A.7.6 and $T_r^{\text{LN}}$ is the density of activated Tregs in the TDLN.

Table A.10 gives us $T_r^{\text{LN}} = 0.26 \times 1.36 \times 10^9 = 3.54 \times 10^8$ cell/cm$^3$ at the initial condition and $T_r^{\text{LN}} = 0.36 \times 1.17 \times 10^9 = 4.21 \times 10^8$ cell/cm$^3$ at steady state. Thus, using (A.6), we also get that at the initial condition $L_0^r = 3.54 \times 10^7$ cell/cm$^3$ and at steady state $L_0^r = 4.21 \times 10^7$. From (A.5), we get that at the initial condition $L_r = 1.61 \times 10^8$ cell/cm$^3$ and at steady state, $L_r = 1.91 \times 10^8$ cell/cm$^3$.

Combining everything gives us the results in Table A.14.

Table A.14: Effector Th1, Th2, naïve Treg and Treg initial conditions (stage III) and steady states (stage IV) in the TDLN. All units are in cell/cm$^3$.

| Variable | Initial Condition | Steady State |
|---|---|---|
| $L_1$ | $2.59 \times 10^7$ | $4.10 \times 10^6$ |
| $L_2$ | $2.04 \times 10^6$ | $4.10 \times 10^6$ |
| $L_0^r$ | $3.54 \times 10^7$ | $4.21 \times 10^7$ |
| $L_r$ | $1.61 \times 10^8$ | $1.91 \times 10^8$ |

We estimate $D_L(0)$ and $\overline{D_L}$ in Appendix A.10.1.

#### A.2 DAMP Initial Conditions and Steady States

##### A.2.1 Estimates for $H$

For HMGB1, $H$, it was found in [21] that in the supernatant of ovarian cancer cells, the HMGB1 concentration was 31.5 ng/mL = $3.15 \times 10^{-8}$ g/mL after 1 hours and 66.89 ng/mL = $6.69 \times 10^{-8}$ g/mL after 24 hours after treatment with cisplatin. The HMGB1 concentration in advanced HGSOC is higher than in early HGSOC due to increased tumour hypoxia and necrosis from cancer progression [22]. Due to the lack of a control, we consider the cisplatin data, and we take $3.15 \times 10^{-8}$ g/mL to be the initial condition for $H$ and take $6.69 \times 10^{-8}$ g/mL to be the steady state.

##### A.2.2 Estimates for $R$

Calreticulin concentration in ovarian cancer, when no drugs were introduced, was approximately $2.00 \times 10^{-2} \pm 2.5 \times 10^{-2}$ $\mu$g/mL [23]. We assume the initial condition to be $2 \times 10^{-8}$ g/cm$^3$ and the steady state to be $4.50 \times 10^{-8}$ g/cm$^3$ as surface calreticulin is produced by necrotic cancer cells which have a higher population at steady state.

#### A.3 Cytokine Initial Conditions and Steady States

We assume that the following values are in the absence of chemotherapy or immunotherapy, that ascites are representative of tumour cytokine concentrations, and that ovarian cancer has a tissue density of 1.03 g/mL [10].

##### A.3.1 Estimates for $I_2$

It was found in [24], that in ovarian cancer patients, the mean IL-2 ascites concentration in ascites was $3.05 \times 10^1$ pg/mL, with the concentration one standard deviation below the mean being

$2.91 \times 10^1$ pg/mL. We assume that the tissue concentration is the same as the ascites concentration. Furthermore, the tissue IL-2 concentration in early HGSOC patients is higher than in advanced HGSOC patients due to the pro-inflammatory nature of IL-2. We thus set the steady state of $I_2$ to $2.91 \times 10^{-11}$ g/cm$^3$.

##### A.3.2 Estimates for $I_4$

It was found in [25], that in ovarian cancer patients undergoing cytoreductive surgery, the mean IL-4 concentration in ascites was 8.11 pg/mL, with the concentration one standard deviation above the mean being 9.46 pg/mL. We assume that the tissue concentration is the same as the ascites concentration. Furthermore, the tissue IL-4 concentration in early HGSOC patients is lower than in advanced HGSOC patients due to the immunosuppressive nature of IL-4. We thus set the steady state of $I_4$ to $9.46 \times 10^{-12}$ g/cm$^3$.

##### A.3.3 Estimates for $I_{10}$

It was found in [26], that in ovarian cancer patients, the mean IL-10 tissue concentration was $1.00 \times 10^{-2}$ pg/$\mu$g, with the concentration one standard deviation above the mean being $1.13 \times 10^{-2}$ pg/$\mu$g. Furthermore, the tissue IL-10 concentration in early HGSOC patients is lower than in advanced HGSOC patients due to the immunosuppressive nature of IL-10. Assuming a tissue density of 1.03 g/mL [10], we thus set the steady state of $I_{10}$ to $1.16 \times 10^{-8}$ g/cm$^3$, with an initial condition of $1.03 \times 10^{-8}$ g/cm$^3$.

##### A.3.4 Estimates for $I_{12}$

It was found in [27] that in advanced ovarian cancer patients with 91% of patients having HGSOC, the median IL-12 tissue concentration was $1.96 \times 10^1$ pg/mL, with the minimum concentration being 3.10 pg/mL. Furthermore, the tissue IL-12 concentration in early HGSOC patients is higher than in advanced HGSOC patients due to the pro-inflammatory nature of IL-12. We thus set the steady state of $I_{12}$ to be $3.10 \times 10^{-12}$ g/cm$^3$, with an initial condition of $1.96 \times 10^{-11}$ g/cm$^3$.

##### A.3.5 Estimates for $I_\gamma$

It was found in [28] that in ovarian cancer patients, the mean IFN-$\gamma$ tissue concentration was $5.42 \times 10^1$ pg/mL, with the concentration one standard deviation below the mean being $5.20 \times 10^1$ pg/mL. Furthermore, the tissue IFN-$\gamma$ concentration in early HGSOC patients is higher than in advanced HGSOC patients due to the pro-inflammatory nature of IFN-$\gamma$. We thus set the steady state of $I_\gamma$ to $5.20 \times 10^{-11}$ g/cm$^3$.

##### A.3.6 Estimates for $I_\alpha$

It was found in [28] that in ovarian cancer patients, the mean $I_\alpha$ tissue concentration was 2.72 pg/mL, with the concentration one standard deviation below the mean being 2.62 pg/mL. Furthermore, the tissue $I_\alpha$ concentration in early HGSOC patients is higher than in advanced HGSOC patients due to the pro-inflammatory nature of $I_\alpha$. We thus set the steady state of $I_\alpha$ to be $2.62 \times 10^{-12}$ g/cm$^3$.

##### A.3.7 Estimates for $I_\beta$

It was found in [29, 30] that in ovarian cancer patients with stage III and IV, the mean TGF-$\beta$ tissue concentration was $1.48 \times 10^4$ pg/mL, with the concentration one standard deviation above the mean being $1.75 \times 10^4$ pg/mL. Furthermore, the tissue TGF-$\beta$ concentration in early HGSOC patients is lower than in advanced HGSOC patients due to the immunosuppressive nature of TGF-$\beta$. We thus set the steady state of TGF-$\beta$ to be $1.75 \times 10^{-8}$ g/cm$^3$.

#### A.4 Half-Saturation Constants

We recall that for some species $X$, $K_X$ denotes the half-saturation constant of $X$, and is a term of the form

$$\frac{X}{K_X + X}.$$

For simplicity, we assume that if $\overline{X}$ denotes the steady-state value of $X$, then

$$\frac{\overline{X}}{K_X + \overline{X}} = \frac{1}{2} \implies K_X = \overline{X}, \tag{A.7}$$

for purely biological components. This implies that

$$\begin{aligned}
K_{T_0^8 L_r} = K_{T_0^4 L_r} = K_{L_8 L_r} = K_{L_1 L_r} = \overline{L_r} &= 1.91 \times 10^8 \text{ g cm}^{-3} \\
K_{T_r I_\beta} = K_{M_2 I_\beta} = \overline{I_\beta} &= 1.75 \times 10^{-8} \text{ g cm}^{-3}, \\
K_{C I_\alpha} = K_{M_1 I_\alpha} = K_{M_2 I_\alpha} = \overline{I_\alpha} &= 2.62 \times 10^{-12} \text{ g cm}^{-3}, \\
K_{C I_\gamma} = K_{M_1 I_\gamma} = \overline{I_\gamma} &= 5.20 \times 10^{-11} \text{ g cm}^{-3}, \\
K_{DH} = \overline{S} &= 6.69 \times 10^{-8} \text{ g cm}^{-3}, \\
K_{DR} = \overline{R} &= 4.50 \times 10^{-8} \text{ g cm}^{-3}, \\
K_{T_8 I_2} = K_{T_1 I_2} = K_{K I_2} = \overline{I_2} &= 2.91 \times 10^{-11} \text{ g cm}^{-3}, \\
K_{T_2 I_4} = K_{M_2 I_4} = \overline{I_4} &= 9.46 \times 10^{-12} \text{ g cm}^{-3}, \\
K_{M_2 I_{10}} = \overline{I_{10}} &= 1.16 \times 10^{-8} \text{ g cm}^{-3}, \\
K_{K I_{12}} = \overline{I_{12}} &= 3.10 \times 10^{-12} \text{ g cm}^{-3}.
\end{aligned}$$

To estimate half-saturation constants for terms related to the chemotherapy agents, we first estimate the $C_{\text{avg}}$ concentration of paclitaxel and carboplatin in serum.

To do this, we used pharmacokinetic parameters from [31], in which advanced ovarian cancer and endometrial cancer patients received 5 mg/mL min AUC carboplatin and 175 mg/m$^2$ of paclitaxel, administered via IV infusion every 3 weeks—over 0.5 hours for carboplatin and over 3 hours for paclitaxel. Patients were also given a weekly oral administration of 60 mg selinexor. The $\text{AUC}_{0-4}$ of paclitaxel was, on average, 3677 ngh/mL whilst the $\text{AUC}_{0-2}$ of carboplatin was, on average, 22282 ngh/mL. Furthermore, it was noted that the $\text{AUC}_{0-4}$ increased almost proportionally with the dose. We assumed this to be the same in tissue, corresponding to $C_{\text{avg}}$ concentrations, which we treat as steady state concentrations, of $9.193 \times 10^{-7}$ g/cm$^3$ and $1.114 \times 10^{-5}$ g/cm$^3$ for paclitaxel and carboplatin, respectively.

For paclitaxel and carboplatin, we assume that in any term of the form $\frac{X}{K_X+X}$, where $K_X$ is the half-saturation constant, then

$$\frac{\overline{X}}{K_X + \overline{X}} = \frac{3}{4} \implies K_X = \frac{\overline{X}}{3}.$$

Thus, we have that

$$K_{CP} = \frac{\overline{P}}{3} = 3.064 \times 10^{-7} \text{ g cm}^{-3},$$
$$K_{CA} = \frac{\overline{P}}{3} = 3.713 \times 10^{-6} \text{ g cm}^{-3}.$$

#### A.5 Inhibition Constants

We recall that for some species $X$, $K_X$ is denoted by the inhibition constant $X$ in a term of the form

$$\frac{1}{1 + X/K_X}.$$

For simplicity, we assume that if $\overline{X}$ denotes the steady-state value of $X$, then

$$\frac{1}{1 + \overline{X}/K_X} = \frac{1}{2} \implies K_X = \overline{X}. \tag{A.8}$$

for purely biological components. This implies that

$$K_{T_8 I_\beta} = K_{DI_\beta} = \overline{I_\beta} = 1.75 \times 10^{-8} \text{ g cm}^{-3},$$
$$K_{T_0^8 L_r} = K_{T_0^4 L_r} = K_{L_8 L_r} = K_{L_1 L_r} = \overline{L_r} = 1.91 \times 10^{8} \text{ cell cm}^{-3},$$
$$K_{KT_r} = K_{T_8 T_r} = K_{T_1 T_r} = K_{I_2 T_r} = K_{I_\gamma T_r} = \overline{T_r} = 1.25 \times 10^{5} \text{ cell cm}^{-3},$$
$$K_{DI_{10}} = K_{T_8 I_{10}} = \overline{I_{10}} = 1.16 \times 10^{-8} \text{ g cm}^{-3}.$$

For paclitaxel and carboplatin, we assume that in any term of the form $\frac{1}{1+X/K_X}$, where $K_X$ is the inhibition constant, then

$$\frac{1}{1 + \overline{X}/K_X} = \frac{1}{4} \implies K_X = \frac{\overline{X}}{3}.$$

Thus, we have that

$$K_{L_8 P^{\text{LN}}} = \frac{\tau_8 \overline{P^{\text{LN}}}}{3} = 4.995 \times 10^{-7} \text{ (g cm}^{-3}\text{) day},$$
$$K_{L_1 P^{\text{LN}}} = \frac{\tau_1 \overline{P^{\text{LN}}}}{3} = 1.266 \times 10^{-6} \text{ (g cm}^{-3}\text{) day},$$
$$K_{L_2 P^{\text{LN}}} = \frac{\tau_2 \overline{P^{\text{LN}}}}{3} = 1.266 \times 10^{-6} \text{ (g cm}^{-3}\text{) day},$$
$$K_{L_r P^{\text{LN}}} = \frac{\tau_r \overline{P^{\text{LN}}}}{3} = 8.795 \times 10^{-7} \text{ (g cm}^{-3}\text{) day},$$
$$K_{L_8 A^{\text{LN}}} = \frac{\tau_8 \overline{A^{\text{LN}}}}{3} = 6.053 \times 10^{-6} \text{ (g cm}^{-3}\text{) day},$$
$$K_{L_1 A^{\text{LN}}} = \frac{\tau_1 \overline{A^{\text{LN}}}}{3} = 1.534 \times 10^{-5} \text{ (g cm}^{-3}\text{) day},$$
$$K_{L_2 A^{\text{LN}}} = \frac{\tau_2 \overline{A^{\text{LN}}}}{3} = 1.534 \times 10^{-5} \text{ (g cm}^{-3}\text{) day},$$
$$K_{L_r A^{\text{LN}}} = \frac{\tau_r \overline{A^{\text{LN}}}}{3} = 1.066 \times 10^{-5} \text{ (g cm}^{-3}\text{) day}.$$

#### A.6 Depletion and Degradation Rates

We recall the formula that the degradation rate of some species $X$, $d_X$, is given by

$$d_X = \frac{\ln 2}{t_{1/2}^X} \tag{A.9}$$

$$= \tau_X \ln 2, \tag{A.10}$$

where $t_{1/2}^X$ is the half-life of $X$, and $\tau_X$ is the mean lifetime of $X$.

##### A.6.1 Estimate for $d_H$

The half-life of HMGB1 was found to be 3 hours in prostate cancer [32]. We assume similar in HGSOC and estimate that,

$$d_H = \frac{\ln 2}{3\text{ h}} = 5.55\text{ d}^{-1},$$

using (A.9).

##### A.6.2 Estimate for $d_R$

The half-life of calreticulin was found to be approximately 12 hours [33]. Therefore, we estimate that

$$d_R = \frac{\ln 2}{12\text{ h}} = 1.39\text{ d}^{-1},$$

using (A.9).

##### A.6.3 Estimate for $d_{D_0}$

In murine models, the lifetime of immature dendritic cells was 28 days [34]. We assume similar in HGSOC and estimate that,

$$d_{D_0} = \frac{1}{28\text{ d}} = 3.57 \times 10^{-2}\text{ d}^{-1},$$

using (A.10).

##### A.6.4 Estimate for $d_D$ and $d_{D_L}$

The half-life of mature dendritic cells is between 1.5–2.9 days [35, 36], and we take it to be 2.2 days in HGSOC. Therefore, we estimate that

$$d_D = \frac{\ln 2}{2.2\text{ d}} = 3.15 \times 10^{-1}\text{ d}^{-1},$$

using (A.9).

We assume the half-life of mature DCs in the TDLN is the same as for mature DCs in the TS, so that

$$d_{D_L} = \frac{\ln 2}{2.2\text{ d}} = 3.15 \times 10^{-1}\text{ d}^{-1}.$$

###### A.6.5 Estimate for $d_{T_0^8}$

The half-life of naïve CD4+ cells in the lymph node is 21.5 days [37]. Therefore, we estimate that

$$d_{T_0^8} = \frac{\ln 2}{21.5 \text{ d}} = 3.22 \times 10^{-2} \text{ d}^{-1},$$

using (A.9).

###### A.6.6 Estimate for $d_{L_8}$ and $d_{T_8}$

We assume that the half-life of effector CD8+ cells in the TDLN is the same as effector CD8+ T cells in the TS, being 77 days [38]. Therefore, we estimate that

$$d_{L_8} = d_{T_8} = \frac{\ln 2}{77 \text{ d}} = 9.00 \times 10^{-3} \text{ d}^{-1},$$

using (A.9).

###### A.6.7 Estimate for $d_{T_0^4}$

The half-life of naïve CD4+ T cells in the lymph node is 17.2 days [37]. Therefore, we estimate that

$$d_{T_0^4} = \frac{\ln 2}{17.2 \text{ d}} = 4.03 \times 10^{-2} \text{ d}^{-1},$$

using (A.9).

###### A.6.8 Estimates for $d_{L_1}$, $d_{L_2}$, $d_{T_1}$ and $d_{T_2}$

We assume that the half-life of effector Th1 and Th2 cells in the TDLN are equal, and the same the half-life of effector Th1 and Th2 cells in the TS, being 87 days [38]. Therefore, we estimate that

$$d_{L_1} = d_{L_2} = d_{T_1} = d_{T_2} = \frac{\ln 2}{87 \text{ d}} = 7.97 \times 10^{-3} \text{ d}^{-1},$$

using (A.9).

###### A.6.9 Estimate for $d_{L_0^r}$

We use the degradation rate of naïve Tregs provided in [39] and assume that $d_{L_0^r} = 2.2 \times 10^{-3}$ day$^{-1}$.

###### A.6.10 Estimate for $d_{L_r}$ and $d_{T_r}$

We assume that the half-life of effector Tregs in the TDLN is the same as for those at the TS, being 11 days [40]. Therefore, we estimate that

$$d_{L_r} = d_{T_r} = \frac{\ln 2}{11 \text{ d}} = 6.30 \times 10^{-2} \text{ d}^{-1},$$

using (A.9).

##### A.6.11 Estimate for $d_{M_0}$

The lifetime of naïve macrophages is 1.37 days [41]. Therefore, we estimate that

$$d_{M_0} = \frac{1}{1.37 \text{ d}} = 7.30 \times 10^{-1} \text{ d}^{-1},$$

using (A.10).

##### A.6.12 Estimate for $d_{M_1}$

The lifetime of classically activated macrophages is between 1–2 days [42], and we take it to be 1.01 days [41]. Therefore, we estimate that

$$d_{M_1} = \frac{1}{1.01 \text{ d}} = 9.90 \times 10^{-1} \text{ d}^{-1},$$

using (A.10).

##### A.6.13 Estimate for $d_{M_2}$

The lifetime of alternatively activated macrophages is 7.41 days [41]. Therefore, we estimate that

$$d_{M_2} = \frac{1}{7.41 \text{ d}} = 1.35 \times 10^{-1} \text{ d}^{-1},$$

using (A.10).

##### A.6.14 Estimate for $d_{K_0}$ and $d_K$

We assume that the half-life of naïve and activated NK cells are the same, being between 7–14 days [43], and we take it to be 10 days [44]. Therefore, we estimate that

$$d_{K_0} = d_K = \frac{\ln 2}{10 \text{ d}} = 6.93 \times 10^{-2} \text{ d}^{-1},$$

using (A.9).

##### A.6.15 Estimate for $d_{I_2}$

The half-life of IL-2 is approximately 5–7 minutes [45], and we take it to be 6.9 minutes [46]. Therefore, we estimate that

$$d_{I_2} = \frac{\ln 2}{6.9 \text{ m}} = 1.446 \times 10^{1} \text{ d}^{-1},$$

using (A.9).

##### A.6.16 Estimate for $d_{I_4}$

The half-life of IL-4 is $19 \pm 2$ minutes [47], and we take it to be 19 minutes. Therefore, we estimate that

$$d_{I_4} = \frac{\ln 2}{19 \text{ m}} = 5.253 \times 10^{1} \text{ d}^{-1},$$

using (A.9).

##### A.6.17 Estimate for $d_{I_{10}}$

The half-life of IL-10 is approximately 2.7–4.5 hours [48], and we take it to be 2.7 hours [49]. Therefore, we estimate that

$$d_{I_{10}} = \frac{\ln 2}{2.7 \text{ h}} = 6.16 \text{ d}^{-1},$$

using (A.9).

##### A.6.18 Estimate for $d_{I_{12}}$

The half-life of IL-12 is between 5.3–9.6 hours [50], and we take it to be the average of 7.45 hours. Therefore, we estimate that

$$d_{I_{12}} = \frac{\ln 2}{7.45 \text{ h}} = 2.23 \text{ d}^{-1},$$

using (A.9).

##### A.6.19 Estimate for $d_{I_\gamma}$

The half-life of IFN-$\gamma$ is 30 minutes [51]. Therefore, we estimate that

$$d_{I_\gamma} = \frac{\ln 2}{30 \text{ m}} = 3.327 \times 10^1 \text{ d}^{-1},$$

using (A.9).

##### A.6.20 Estimate for $d_{I_\alpha}$

The half-life of TNF is approximately 10–20 minutes [52, 53], and we take it to be 18.2 minutes [54]. Therefore, we estimate that

$$d_{I_\alpha} = \frac{\ln 2}{18.2 \text{ m}} = 5.484 \times 10^1 \text{ d}^{-1},$$

using (A.9).

##### A.6.21 Estimate for $d_{I_\beta}$

The half-life of TGF-$\beta$ is between 2–3 minutes [55], and we take it to be 2 minutes [56]. Therefore, we estimate that

$$d_{I_\beta} = \frac{\ln 2}{2 \text{ m}} = 4.991 \times 10^2 \text{ d}^{-1},$$

using (A.9).

##### A.6.22 Estimate for $d_P$

The mean terminal elimination half-life of paclitaxel for a 3-hour infusion is approximately 9.9 hours [57]. Therefore, we estimate that

$$d_P = \frac{\ln 2}{9.9 \text{ h}} = 1.680 \text{ d}^{-1},$$

using (A.9).

##### A.6.23 Estimate for $d_A$

The estimation for the degradation rate of carboplatin is more complicated than other species due to the involved pharmacokinetics. The half-life of carboplatin itself is approximately 1.5 hours; however, approximately 87% of the platinum from carboplatin binds irreversibly to plasma proteins and is slowly eliminated with a minimum half-life of 5 days [58]. To accommodate for this, we follow [59] and assume the rate of carboplatin degradation to follow first-order kinetics. In particular, [60] reported that the decay rate of carboplatin was unaffected by the presence of cells and remained approximately linear in the presence of Jurkat cell concentrations below $10^7$ cells/mL. The first-order decay rate of carboplatin was found to be approximately $1.94 \times 10^{-6}\ \mathrm{s}^{-1} = 1.676 \times 10^{-1}\ \mathrm{d}^{-1}$. Thus, we estimate $d_A = 1.676 \times 10^{-1}\ \mathrm{d}^{-1}$.

##### A.6.24 Estimates for $d_{PC}$

We assume that at steady state, 90% of paclitaxel is used in eliminating cancer cells, while the remaining 10% degrades naturally. Thus,

$$\frac{d_{PC}\overline{PC}}{90\%} = \frac{d_P\overline{P}}{10\%} \implies d_{PC} = \frac{9d_P}{\overline{C}} = 5.196 \times 10^{-7}\ (\mathrm{cell/cm^3})^{-1}\mathrm{day}^{-1}.$$

##### A.6.25 Estimates for $d_{AC}$

We assume that at steady state, 90% of carboplatin is used in eliminating cancer cells, while the remaining 10% degrades naturally. Thus,

$$\frac{d_{AC}\overline{AC}}{90\%} = \frac{d_A\overline{A}}{10\%} \implies d_{AC} = \frac{9d_A}{\overline{C}} = 5.184 \times 10^{-8}\ (\mathrm{cell/cm^3})^{-1}\mathrm{day}^{-1}.$$

##### A.6.26 Estimates for $d_{P^{\mathrm{LN}}L_8}$, $d_{P^{\mathrm{LN}}L_1}$, $d_{P^{\mathrm{LN}}L_2}$, and $d_{P^{\mathrm{LN}}L_r}$

We assume that at steady state, 20% of paclitaxel is used in eliminating T cells, while the remaining 80% degrades naturally. Furthermore, we assume that the depletion rates of paclitaxel by T cells are proportional to their concentrations, so that

$$\frac{d_{P^{\mathrm{LN}}L_8}\overline{P^{\mathrm{LN}}L_8} + d_{P^{\mathrm{LN}}L_1}\overline{P^{\mathrm{LN}}L_1} + d_{P^{\mathrm{LN}}L_2}\overline{P^{\mathrm{LN}}L_2} + d_{P^{\mathrm{LN}}L_r}\overline{P^{\mathrm{LN}}L_r}}{20\%} = \frac{d_P\overline{P^{\mathrm{LN}}}}{80\%},$$
$$\frac{d_{P^{\mathrm{LN}}L_8}}{\overline{L_8}} = \frac{d_{P^{\mathrm{LN}}L_1}}{\overline{L_1}} = \frac{d_{P^{\mathrm{LN}}L_2}}{\overline{L_2}} = \frac{d_{P^{\mathrm{LN}}L_r}}{\overline{L_r}}.$$

Solving these simultaneously leads to

$$\begin{aligned}
d_{P^{\mathrm{LN}}L_8} &= 3.347 \times 10^{-9}\ (\mathrm{cell/cm^3})^{-1}\mathrm{day}^{-1},\\
d_{P^{\mathrm{LN}}L_1} &= 3.012 \times 10^{-9}\ (\mathrm{cell/cm^3})^{-1}\mathrm{day}^{-1},\\
d_{P^{\mathrm{LN}}L_2} &= 3.012 \times 10^{-9}\ (\mathrm{cell/cm^3})^{-1}\mathrm{day}^{-1},\\
d_{P^{\mathrm{LN}}L_r} &= 2.008 \times 10^{-9}\ (\mathrm{cell/cm^3})^{-1}\mathrm{day}^{-1}.
\end{aligned}$$

##### A.6.27 Estimates for $d_{A^{\mathrm{LN}}L_8}$, $d_{A^{\mathrm{LN}}L_1}$, $d_{A^{\mathrm{LN}}L_2}$, and $d_{A^{\mathrm{LN}}L_r}$

We assume that at steady state, 20% of carboplatin is used in eliminating T cells, while the remaining 80% degrades naturally. Furthermore, we assume that the depletion rates of carboplatin by T cells are proportional to their concentrations, so that

$$\frac{d_{A^{\mathrm{LN}}L_8}\overline{A^{\mathrm{LN}}}\,\overline{L_8}+d_{A^{\mathrm{LN}}L_1}\overline{A^{\mathrm{LN}}}\,\overline{L_1}+d_{A^{\mathrm{LN}}L_2}\overline{A^{\mathrm{LN}}}\,\overline{L_2}+d_{A^{\mathrm{LN}}L_r}\overline{A^{\mathrm{LN}}}\,\overline{L_r}}{20\%}=\frac{d_A\overline{A^{\mathrm{LN}}}}{80\%},$$

$$\frac{d_{A^{\mathrm{LN}}L_8}}{\overline{L_8}}=\frac{d_{A^{\mathrm{LN}}L_1}}{\overline{L_1}}=\frac{d_{A^{\mathrm{LN}}L_2}}{\overline{L_2}}=\frac{d_{A^{\mathrm{LN}}L_r}}{\overline{L_r}}.$$

Solving these simultaneously leads to

$$\begin{aligned}
d_{A^{\mathrm{LN}}L_8} &= 3.339\times10^{-10}\ (\mathrm{cell/cm^3})^{-1}\mathrm{day}^{-1},\\
d_{A^{\mathrm{LN}}L_1} &= 3.005\times10^{-10}\ (\mathrm{cell/cm^3})^{-1}\mathrm{day}^{-1},\\
d_{A^{\mathrm{LN}}L_2} &= 3.005\times10^{-10}\ (\mathrm{cell/cm^3})^{-1}\mathrm{day}^{-1},\\
d_{A^{\mathrm{LN}}L_r} &= 2.003\times10^{-10}\ (\mathrm{cell/cm^3})^{-1}\mathrm{day}^{-1}.
\end{aligned}$$

#### A.7 TDLN Subsystem Parameters

##### A.7.1 Estimate for $V_{\mathrm{TS}}$

In stage III ovarian cancer, the average tumour size is 4.60 cm [61]. Assuming sphericity the volume of the tumour is $\frac{4}{3}\times\pi\times(2.3)^3\ \mathrm{cm}^3\approx 51.0\ \mathrm{cm}^3$.

##### A.7.2 Estimate for $V_{\mathrm{LN}}$

In [62], in stage III ovarian cancer, the median length of the long and short axes of positive lymph nodes was 10.0 mm and 5.0 mm, respectively. For simplicity, we assume the lymph node to be spherical with radius equal to the geometric mean of the long and short axis, i.e., $\sqrt{5\times2.5}$. Assuming sphericity the volume of the tumour is $\frac{4}{3}\times\pi\times(\sqrt{5\times2.5})^3\ \mathrm{mm}^3\approx 0.185\ \mathrm{cm}^3$.

##### A.7.3 Estimate for $\tau_m$

In [63], dendritic cells required approximately 18 hours to migrate from a subcutaneous infusion site to the TDLN after acquiring antigen. We assume a similar migration time for dendritic cells that acquire cancer antigens at the TS, setting $\tau_m = 18\ \mathrm{h} = 0.75\ \mathrm{d}$.

##### A.7.4 Estimates for CD8+ T Cells

In [64], it was reported that activated CD8+ T cells typically undergo a minimum of 7–10 divisions during clonal expansion, however, under conditions of persistent antigen exposure, the number of divisions can increase significantly. For instance, in the case of Lymphocytic Choriomeningitis Virus (LCV), up to 19 divisions have been observed [65]. Accordingly, in our model we set the number of cell divisions, $N_8$, to be 10 for CD8+ T cells. In [66], it was found that activated CD8+ T cells required 39 hours on average to complete their first cell division, and 8.6 hours to complete subsequent divisions, though this can range from 5 to 28 hours.
To estimate $\tau_8$, the time for CD8+ T cells to complete clonal expansion, we assume a constant cell cycle time $\Delta_8$, except for the first cell division which has a cycle time of $\Delta_0^8$, i.e.,

$$\tau_8 = \Delta_0^8 + (N_8-1)\Delta_8. \tag{A.11}$$

Thus, setting $\Delta_0^8 = 39\ \mathrm{h} = 1.63\ \mathrm{d}$, and $\Delta_8$, to be $8.6\ \mathrm{h} = 0.36\ \mathrm{d}$, we have that $\tau_8 = 116.4\ \mathrm{h} = 4.85\ \mathrm{d}$.

##### A.7.5 Estimates for CD4+ T Cells

In [67], CD4+ T cells undergo approximately 9 divisions during clonal expansion in response to LCV. We assume a similar process occurs in HGSOC and set the number of cell divisions for CD4+ cells, $N_1$, to be 9. The first CD4+ T cell division, denoted, $\Delta_0^4$, typically occurs within 12 to 24 hours, and so we take it to be and we take the average, so that $\Delta_0^4 = 18.5$ h $= 0.77$ d [68]. Subsequent divisions, take place at an average rate of approximately 10 hours per division, denoting this $\Delta_4$, so that $\Delta_4 = 10$ h $= 0.42$ d.

In order to estimate $\tau_1$ and $\tau_2$ we consider,

$$\tau_1 = \tau_2 = \Delta_0^4 + (N_1 - 1)\Delta_4. \tag{A.12}$$

This gives us $\tau_1 = \tau_2 = 4.13$ d.

##### A.7.6 Estimates for Tregs

In [69], it was observed that 6 days after tumour implantation in mice, 45% of Tregs in the TDLN had undergone at least one division, with 14% having undergone more than six divisions. In order to estimate $\tau_r$ we consider,

$$\tau_r = \Delta_0^r + (N_r - 1)\Delta_r. \tag{A.13}$$

where we set $N_r$, the number of cell divisions for Treg cells, to be 6, and assume that the first and subsequent divisions rates of Tregs, $\Delta_0^r$ and $\Delta_r$, respectively, are the same as those for CD4+ T helper cells. This gives us $\Delta_0^r = 0.77$ d and $\Delta_r = 0.42$ d. Consequently, we have that $\tau_r = 2.87$ d.

##### A.7.7 Estimates for $\tau_a$, $\tau_b$, $\tau_c$, $\tau_d$

In [70], it was found that T cells in the TDLN migrate at speeds between 11–14 $\mu$m/min, while DCs migrate at speeds between 3–6 $\mu$m/min. We assume these migration speeds are the same for travel from the TDLN to the TS. Using this, we obtain the ratio: $\tau_a = \frac{4.5}{12.5}\tau_m \approx 0.27$ d. We thus set $\tau_a = \tau_b = \tau_c = \tau_d = 0.27$ d.

#### A.8 DAMP Parameters

##### A.8.1 Estimates for $H$

Considering (2.3) at steady state, we have that

$$\lambda_{NH}\overline{N} - d_S\overline{S} = 0 \implies \lambda_{NH} = 1.263 \times 10^{-12}\,(\text{g/cell})\,\text{day}^{-1}.$$

##### A.8.2 Estimates for $R$

Considering (2.4) at steady state, we have that

$$\lambda_{NR}\overline{N} - d_R\overline{R} = 0 \implies \lambda_{NR} = 2.128 \times 10^{-13}\,(\text{g/cell})\,\text{day}^{-1}.$$

#### A.9 Cytokine Production Parameters

##### A.9.1 Estimates for $I_2$

Using (2.24), we get that at steady state

$$\lambda_{I_2T_1}\frac{\overline{T_1}}{2}+\lambda_{I_2T_8}\frac{\overline{T_8}}{2}-d_{I_2}\overline{I_2}=0.$$

Using values from [71] we get that

$$\frac{\lambda_{I_2T_1}}{2\times 0.335763694}=\frac{\lambda_{I_2T_8}}{2\times 0.114615876}.$$

Solving these simultaneously gives us

$$\lambda_{I_2T_1}=9.957\times 10^{-14}\ \ (\text{g/cell})\,\text{day}^{-1},$$
$$\lambda_{I_2T_8}=3.399\times 10^{-14}\ \ (\text{g/cell})\,\text{day}^{-1}.$$

Consequently, considering (2.35), we have that

$$I_2(0)=\frac{1}{d_{I_2}}\left[\left(\lambda_{I_2T_8}T_8(0)+\lambda_{I_2T_1}T_1(0)\right)\frac{1}{1+T_r(0)/K_{I_2T_r}}\right]=3.05\times 10^{-11}\ \text{g/cm}^3.$$

##### A.9.2 Estimates for $I_4$

Using (2.25), we get that at steady state

$$\lambda_{I_4T_2}\overline{T_2}-d_{I_4}\overline{I_4}=0\implies\lambda_{I_4T_2}=8.749\times 10^{-15}\ \ (\text{g/cell})\,\text{day}^{-1}.$$

Consequently, considering (2.36), we have that

$$I_4(0)=\frac{\lambda_{I_4T_2}}{d_{I_4}}T_2(0)=8.11\times 10^{-12}\ \text{g/cm}^3.$$

##### A.9.3 Estimates for $I_{10}$

Using (2.26), we get that at steady state

$$\lambda_{I_{10}T_r}\overline{T_r}+\lambda_{I_{10}M_2}\overline{M_2}+\lambda_{I_{10}C}\overline{C}-d_{I_{10}}\overline{I_{10}}=0.$$

Using values from [71] we get that

$$\frac{\lambda_{I_{10}T_r}}{0.472157631}=\frac{\lambda_{I_{10}M_2}}{1}$$

In ovarian cancer, myeloid cells are the main producers of IL-10 [72]. Ovarian cancer cells also produce IL-10, and we assume they produce half the IL-10 of macrophages. Hence,

$$\frac{\lambda_{I_{10}C}}{1}=\frac{\lambda_{I_{10}M_2}}{2}$$

Solving these simultaneously gives us

$$\lambda_{I_{10}T_r}=2.204\times 10^{-15}\ \ (\text{g/cell})\,\text{day}^{-1},$$
$$\lambda_{I_{10}M_2}=4.667\times 10^{-15}\ \ (\text{g/cell})\,\text{day}^{-1},$$
$$\lambda_{I_{10}C}=2.334\times 10^{-15}\ \ (\text{g/cell})\,\text{day}^{-1}.$$

##### A.9.4 Estimates for $I_{12}$

Using (2.27), we get that at steady state

$$\lambda_{I_{12}D}\overline{D} + \lambda_{I_{12}M_1}\overline{M_1} - d_{I_{12}}\overline{I_{12}} = 0.$$

Using values from [71] we get that

$$\frac{\lambda_{I_{12}D}}{0.330046991} = \frac{\lambda_{I_{12}M_1}}{0.37592202}.$$

Solving these simultaneously gives us

$$\begin{aligned} \lambda_{I_{12}D} &= 7.05 \times 10^{-18} \ \ (\text{g/cell})\,\text{day}^{-1}, \\ \lambda_{I_{12}M_1} &= 8.03 \times 10^{-18} \ \ (\text{g/cell})\,\text{day}^{-1}. \end{aligned}$$

##### A.9.5 Estimates for $I_\gamma$

Using (2.28), we get that at steady state,

$$\lambda_{I_\gamma T_1}\frac{\overline{T_1}}{2} + \lambda_{I_\gamma K}\frac{\overline{K}}{2} + \lambda_{I_\gamma T_8}\overline{T_8} - d_{I_\gamma}\overline{I_\gamma} = 0.$$

Using values from [71] we get that

$$\frac{\lambda_{I_\gamma T_1}}{2 \times 0.018892673} = \frac{\lambda_{I_\gamma K}}{2 \times 1} = \frac{\lambda_{I_\gamma T_8}}{0.053997331}.$$

Solving these simultaneously gives us

$$\begin{aligned} \lambda_{I_\gamma T_1} &= 4.202 \times 10^{-17} \ \ (\text{g/cell})\,\text{day}^{-1}, \\ \lambda_{I_\gamma K} &= 2.224 \times 10^{-15} \ \ (\text{g/cell})\,\text{day}^{-1}, \\ \lambda_{I_\gamma T_8} &= 6.001 \times 10^{-17} \ \ (\text{g/cell})\,\text{day}^{-1}. \end{aligned}$$

Consequently, considering (2.37), we have that

$$I_\gamma(0) = \frac{1}{d_{I_\gamma}}\left[\big(\lambda_{I_\gamma T_1}T_1(0) + \lambda_{I_\gamma K}K(0)\big)\frac{1}{1 + T_r(0)/K_{I_\gamma T_r}} + \lambda_{I_\gamma T_8}T_8(0)\right] = 6.11 \times 10^{-11} \ \text{g/cm}^3.$$

##### A.9.6 Estimates for $I_\alpha$

Using (2.2.26), we get that at steady state

$$\lambda_{I_\alpha T_8}\overline{T_8} + \lambda_{I_\alpha T_1}\overline{T_1} + \lambda_{I_\alpha K}\overline{K} + \lambda_{I_\alpha M_1}\overline{M_1} + \lambda_{I_\alpha D}\overline{D} - d_{I_\alpha}\overline{I_\alpha} = 0.$$

Using values from [71] we get that

$$\frac{I_\alpha T_8}{0.065444378} = \frac{\lambda_{I_\alpha T_1}}{0.108187215} = \frac{\lambda_{I_\alpha K}}{0.114108294} = \frac{\lambda_{I_\alpha M_1}}{0.039657574} = \frac{\lambda_{I_\alpha D}}{0.199223795}.$$

Solving these simultaneously gives us

$$\lambda_{I_\alpha T_8} = 2.712 \times 10^{-17} \ \ (\text{g/cell})\,\text{day}^{-1},$$

$$\lambda_{I_\alpha T_1} = 4.483 \times 10^{-17} \text{ (g/cell) day}^{-1},$$
$$\lambda_{I_\alpha K} = 4.729 \times 10^{-17} \text{ (g/cell) day}^{-1},$$
$$\lambda_{I_\alpha M_1} = 1.643 \times 10^{-17} \text{ (g/cell) day}^{-1},$$
$$\lambda_{I_\alpha D} = 8.256 \times 10^{-17} \text{ (g/cell) day}^{-1}.$$

Consequently, considering (2.38), we have that

$$I_\alpha(0) = \frac{1}{d_{I_\alpha}} \left(\lambda_{I_\alpha T_8} T_8(0) + \lambda_{I_\alpha T_1} T_1(0) + \lambda_{I_\alpha K} K(0) + \lambda_{I_\alpha M_1} M_1(0) + \lambda_{I_\alpha D} D(0)\right) = 2.72 \times 10^{-12} \text{ g/cm}^3.$$

##### A.9.7 Estimates for $I_\beta$

Using (2.2.27), we get that at steady state

$$\lambda_{I_\beta T_r}\overline{T_r} + \lambda_{I_\beta C}\overline{C} + \lambda_{I_\beta M_2}\overline{M_2} + \lambda_{I_\beta K}\overline{K} - d_{I_\beta}\overline{I_\beta} = 0.$$

Using values from [71] we get that

$$\frac{\lambda_{I_\beta T_r}}{0.507677682} = \frac{\lambda_{I_\beta M_2}}{0.630703572} = \frac{\lambda_{I_\beta F}}{0.175283003} = \frac{\lambda_{I_\beta K}}{0.429072356}.$$

Solving these simultaneously gives us

$$\lambda_{I_\beta T_r} = 1.191 \times 10^{-12} \text{ (g/cell) day}^{-1},$$
$$\lambda_{I_\beta C} = 2.057 \times 10^{-13} \text{ (g/cell) day}^{-1},$$
$$\lambda_{I_\beta M_2} = 1.48 \times 10^{-12} \text{ (g/cell) day}^{-1},$$
$$\lambda_{I_\beta K} = 1.007 \times 10^{-12} \text{ (g/cell) day}^{-1}.$$

Consequently, considering (2.39), we have that

$$I_\beta(0) = \frac{1}{d_{I_\beta}} \left(\lambda_{I_\beta T_r} T_r(0) + \lambda_{I_\beta C} C(0) + \lambda_{I_\beta M_2} M_2(0) + \lambda_{I_\beta K} K(0)\right) = 1.48 \times 10^{-8} \text{ g/cm}^3.$$

#### A.10 Parameters for DCs, Macrophages, and NK Cells

##### A.10.1 Estimates for $D_0$, $D$ and $D_L$

Considering (2.5) at steady state, we have that

$$\mathcal{A}_{D_0} - \zeta_{DH}\frac{\overline{D_0}}{8} - \zeta_{DR}\frac{\overline{D_0}}{8} - d_{D_0}\overline{D_0} = 0.$$

We assume that HMGB1 is a more potent inducer of DC maturation than surface calreticulin. As such, at steady state, we assume that

$$\zeta_{DH}/10 = \zeta_{DR}.$$

Considering (2.6) at steady state, we have that

$$\zeta_{DH}\frac{\overline{D_0}}{8} + \zeta_{DR}\frac{\overline{D_0}}{8} - \lambda_{D_L D}\overline{D} - d_D\overline{D} = 0.$$

Considering (2.7) at steady state, we have that,

$$\frac{V_{\text{TS}}}{V_{\text{LN}}} e^{-d_D \tau_m} \lambda_{D_L D} \overline{D} - d_{D_L} \overline{D_L} = 0$$

In [73], it was found that the rate of migration of DCs through afferent lymph is $10^4$–$10^5$ (cell cm$^3$)$^{-1}$ h$^{-1}$. We assume this is the same rate at which immature DCs enter the TS, so that

$$\begin{aligned}\mathcal{A}_{D_0} &= 3.00 \times 10^4 \text{ cell cm}^{-3} \text{ h}^{-1}\\ &= 7.20 \times 10^5 \text{ cell cm}^{-3} \text{ day}^{-1}.\end{aligned}$$

Solving these simultaneously gives us

$$\begin{aligned}\zeta_{DH} &= 8.831 \text{ day}^{-1},\\ \zeta_{DR} &= 0.883 \text{ day}^{-1},\\ \lambda_{D_L D} &= 6.114 \times 10^{-1} \text{ day}^{-1}.\end{aligned}$$

Considering (2.7), we get that at steady state

$$\frac{V_{\text{TS}}}{V_{\text{TS}}} e^{-d_D \tau_m} \lambda_{D_L D} \overline{D} - d_{D_L} \overline{D_L} = 0.$$

Solving this gives us

$$\begin{aligned}\overline{D_L} &= \frac{\frac{V_{\text{TS}}}{V_{\text{TS}}} e^{-d_D \tau_m} \lambda_{D_L D} \overline{D}}{d_{D_L}},\\ &= 3.190 \times 10^8 \text{ cell/cm}^3.\end{aligned}$$

To estimate the initial condition of $D_L$, we assume that the ratio of mature DCs in the TDLN at the initial condition to steady state is the same as the ratio of mature DCs in the TS at the initial condition to steady state. Thus we have that

$$\frac{D_L(0)}{\overline{D_L}} = \frac{D(0)}{\overline{D}},$$

so that

$$\begin{aligned}D_L(0) &= \frac{6.60 \times 10^5}{7.55 \times 10^5} \times 3.190 \times 10^8\\ &= 2.729 \times 10^8 \text{ cell/cm}^3.\end{aligned}$$

##### A.10.2 Estimates for $M_0$, $M_1$, and $M_2$

Adding (2.19), (2.20), and (2.21) at steady state, leads to

$$\mathcal{A}_{M_0} - d_{M_0} \overline{M_0} - d_{M_1} \overline{M_1} - d_{M_2} \overline{M_2} = 0 \implies \mathcal{A}_{M_0}.$$

Using values from [71], and considering (2.19) at steady state, leads to the equations

$$\mathcal{A}_{M_0} - \zeta_{M_1 I_\gamma} \frac{\overline{M_0}}{2} - \zeta_{M_1 I_\alpha} \frac{\overline{M_0}}{2} - \zeta_{M_2 I_4} \frac{\overline{M_0}}{2} - \zeta_{M_2 I_{10}} \frac{\overline{M_0}}{2} - d_{M_0} \overline{M_0} = 0,$$

$$\frac{\zeta_{M_1 I_\gamma}/2}{12.54} = \frac{\zeta_{M_1 I_\alpha}/2}{10.77} = \frac{\zeta_{M_2 I_4}/2}{7.39} = \frac{\zeta_{M_2 I_{10}}/2}{6.81}.$$

Considering (2.20) at steady state, we have that

$$\zeta_{M_1 I_\gamma}\frac{\overline{M_0}}{2} + \zeta_{M_1 I_\alpha}\frac{\overline{M_0}}{2} + \zeta_{M I_\gamma}\frac{\overline{M_2}}{2} + \zeta_{M I_\alpha}\frac{\overline{M_2}}{2} - \zeta_{M I_\beta}\frac{\overline{M_1}}{2} - d_{M_1} M_1 = 0,$$

As in [1], we also assume,

$$\frac{1}{\overline{M_2}}\left(\frac{\zeta_{MI_\gamma}}{2} + \frac{\zeta_{MI_\alpha}}{2}\right) = \frac{1}{\overline{M_1}}\frac{\zeta_{MI_\beta}}{2},$$

and that the rate of re-polarisation of M1 to M2 macrophages by IFN-$\gamma$ is slightly more potent than TNF. Hence,

$$\frac{\lambda_{MI_\gamma}}{2\times 5} = \frac{\lambda_{MI_\beta}}{2\times 6}.$$

Solving these simultaneously gives us

$$\begin{aligned}
\mathcal{A}_{M_0} &= 5.551\times 10^{5} \text{ cell cm}^{-3} \text{ day}^{-1},\\
\zeta_{M_1 I_\gamma} &= 5.371\times 10^{-1} \text{ day}^{-1},\\
\zeta_{M_1 I_\alpha} &= 4.613\times 10^{-1} \text{ day}^{-1},\\
\zeta_{M_2 I_4} &= 3.165\times 10^{-1} \text{ day}^{-1},\\
\zeta_{M_2 I_{10}} &= 2.197\times 10^{-1} \text{ day}^{-1},\\
\zeta_{M I_\alpha} &= 2.154\times 10^{-2} \text{ day}^{-1},\\
\zeta_{M I_\beta} &= 1.337\times 10^{-2} \text{ day}^{-1},\\
\zeta_{M I_\gamma} &= 2.585\times 10^{-2} \text{ day}^{-1}.
\end{aligned}$$

##### A.10.3 Estimates for $K_0$ and $K$

Using (2.22), we get that at steady state

$$\mathcal{A}_{K_0} - \zeta_{KI_2}\frac{\overline{K_0}}{2} - \zeta_{KI_{12}}\frac{\overline{K_0}}{2} - d_{K_0}\overline{K_0} = 0.$$

Using values from [71] gives us

$$\frac{\zeta_{KI_2}/2}{15.4} = \frac{\zeta_{KI_{12}}/2}{14.83}.$$

Using (2.23), we get that at steady state

$$\zeta_{KI_2}\frac{\overline{K_0}}{2} + \zeta_{KI_{12}}\frac{\overline{K_0}}{2} + \lambda_{KI_2}\frac{\overline{K}}{2} + \lambda_{KI_{12}}\frac{\overline{K}}{2} - d_K\overline{K} = 0.$$

We assume that the above ratio for NK cell activation is maintained for proliferation at steady state, so that

$$\frac{\lambda_{KI_2}/2}{15.4} = \frac{\lambda_{KI_{12}}/2}{14.83}.$$

In healthy adults, the proliferation rate of NK cells is 4.3% per day [44]. However, in the ovarian cancer tumour micro-environment, NK cell function is impaired [74]. We assume that it impairs the proliferation rate by 50%. This gives us

$$\frac{\zeta_{KI_2}K_0/2 + (\zeta_{KI_{12}}K_0)/2}{0.9785} = \frac{(\lambda_{KI_2}K/2) + (\lambda_{KI_{12}}K/2)}{0.0215}$$

Solving these simultaneously gives us

$$\begin{aligned}
\mathcal{A}_{K_0} &= 1.362 \times 10^5 \text{ cell cm}^{-3} \text{ day}^{-1}, \\
\zeta_{KI_2} &= 2.385 \times 10^{-1} \text{ day}^{-1}, \\
\zeta_{KI_{12}} &= 2.297 \times 10^{-1} \text{ day}^{-1}, \\
\lambda_{KI_2} &= 1.518 \times 10^{-3} \text{ day}^{-1}, \\
\lambda_{KI_{12}} &= 1.462 \times 10^{-3} \text{ day}^{-1}.
\end{aligned}$$

#### A.11 T Cell Parameters

##### A.11.1 Estimates for $T_0^8$, $L_8$ and $T_8$

Considering (2.8) at steady state, we have that

$$\mathcal{A}_{T_0^8} - \frac{\zeta_{L_8 D_L} \overline{T_0^8}\,\overline{D_L}}{2} - d_{T_0^8} \overline{T_0^8} = 0.$$

Considering (2.9) at steady state without chemotherapy, we have that

$$\frac{e^{-d_{L_8}\tau_8} 2^{N_8} \zeta_{L_8 D_L} \overline{T_0^8}\,\overline{D_L}}{2} - \zeta_{T_8 L_8} \overline{L_8} - d_{L_8} \overline{L_8} = 0.$$

Considering (2.10) at steady state, we have that,

$$\frac{V_{\text{LN}}}{V_{\text{TS}}} e^{-d_{L_8}\tau_a} \zeta_{T_8 L_8} \overline{L_8} + \frac{\lambda_{T_8 I_2} \overline{T_8}}{4} - \frac{d_{T_8} \overline{T_8}}{2} = 0.$$

As in [1], we also assume that at steady state, 5% of positive $T_8$ growth is due to IL-2 induced proliferation and 95% is due to $L_8$ migration to the TS. Hence, we have that,

$$\frac{\frac{V_{\text{LN}}}{V_{\text{TS}}} e^{-d_{L_8}\tau_a} \zeta_{T_8 L_8} \overline{L_8}}{0.95} = \frac{\lambda_{T_8 I_2} \overline{T_8}/4}{0.05}.$$

Solving these simultaneously at steady state gives us

$$\begin{aligned}
\mathcal{A}_{T_0^8} &= 2.486 \times 10^{6} \text{ cell cm}^{-3} \text{ day}^{-1}, \\
\zeta_{L_8 D_L} &= 1.103 \times 10^{-14} \text{ (cell cm}^{-3})^{-1} \text{day}^{-1}, \\
\zeta_{T_8 L_8} &= 2.891 \times 10^{-2} \text{ day}^{-1}, \\
\lambda_{T_8 I_2} &= 9.00 \times 10^{-4} \text{ day}^{-1}.
\end{aligned}$$

##### A.11.2 Estimates for $T_0^4$, $L_1$, $T_1$, $L_2$ and $T_2$

Considering (2.11) at steady state, we have that

$$\mathcal{A}_{T_0^4} - \frac{\zeta_{L_1 D_L} \overline{T_0^4}\,\overline{D_L}}{2} - \frac{\zeta_{L_2 D_L} \overline{T_0^4}\,\overline{D_L}}{2} - d_{T_0^4} \overline{T_0^4} = 0.$$

Considering (2.12) at steady state without chemotherapy, we have that

$$\frac{e^{-d_{L_1}\tau_1} 2^{N_1} \zeta_{L_1 D_L} \overline{T_0^4}\,\overline{D_L}}{2} - \zeta_{T_1 L_1} \overline{L_1} - d_{L_1} \overline{L_1} = 0.$$

Considering (2.13) at steady state, we have that

$$\frac{V_{\text{LN}}}{V_{\text{TS}}} e^{-d_{L_1}\tau_b} \zeta_{T_1 L_1} \overline{L_1} + \frac{\lambda_{T_1 I_2} \overline{T_1}}{2} - d_{T_1} \overline{T_1} = 0$$

As in [1], we also assume that at steady state, 5% of positive $T_1$ growth is due to IL-2 induced proliferation and 95% is due to $L_1$ migration to the TS. Hence, we have that

$$\frac{\frac{V_{\text{LN}}}{V_{\text{TS}}} e^{-d_{L_1}\tau_b} \zeta_{T_1 L_1} \overline{L_1}}{0.95} = \frac{\lambda_{T_1 I_2} \overline{T_1}/2}{0.05}.$$

Considering (2.14) at steady state without chemotherapy, we have that

$$\frac{e^{-d_{L_2}\tau_2}2^{N_2}\zeta_{L_2D_L}\overline{T_0^4D_L}}{2} - \zeta_{T_2L_2}\overline{L_2} - d_{L_2}\overline{L_2} = 0.$$

Considering (2.15) at steady state without chemotherapy, we have that

$$\frac{V_{\text{LN}}}{V_{\text{TS}}}e^{-d_{T_2}\tau_c}\zeta_{T_2L_2}\overline{L_2} + \frac{\lambda_{T_2I_4}\overline{T_2}}{2} - d_{T_2}\overline{T_2} = 0.$$

We also assume that at steady state, 5% of positive $T_2$ growth is due to IL-4 induced proliferation and 95% is due to $L_2$ migration to the TS. Hence, we have that

$$\frac{\frac{V_{\text{LN}}}{V_{\text{TS}}}e^{-d_{L_2}\tau_c}\zeta_{T_2L_2}\overline{L_2}}{0.95} = \frac{\lambda_{T_2I_4}\overline{T_2}/2}{0.05}.$$

Solving these simultaneously gives us

$$\begin{aligned}
\mathcal{A}_{T_0^4} &= 2.547 \times 10^6 \text{ cell cm}^{-3} \text{ day}^{-1},\\
\zeta_{L_1D_L} &= 2.967 \times 10^{-14} \text{ (cell cm}^{-3})^{-1}\text{day}^{-1},\\
\zeta_{L_2D_L} &= 1.517 \times 10^{-14} \text{ (cell cm}^{-3})^{-1}\text{day}^{-1},\\
\zeta_{T_1L_1} &= 2.816 \times 10^{-2} \text{ day}^{-1},\\
\zeta_{T_2L_2} &= 2.898 \times 10^{-2} \text{ day}^{-1},\\
\lambda_{T_1I_2} &= 7.970 \times 10^{-4} \text{ day}^{-1},\\
\lambda_{T_2I_4} &= 3.985 \times 10^{-4} \text{ day}^{-1}.
\end{aligned}$$

##### A.11.3 Estimates for $L_0^r$, $L_r$ and $T_r$

Considering (2.16) at steady state, we have that

$$\mathcal{A}_T - \zeta_{L_rD_L}\overline{L_0^rD_L} - d_{L_0^r}\overline{L_0^r} = 0.$$

Considering (2.17) at steady state without chemotherapy, we have that

$$e^{-d_{L_r}\tau_r}2^{N_r}\zeta_{L_rD_L}\overline{L_0^rD_L} - \zeta_{T_rL_r}\overline{L_r} - d_{L_r}\overline{L_r} = 0.$$

Considering (2.18) at steady state, we have that,

$$\frac{V_{\text{LN}}}{V_{\text{TS}}}e^{-d_{T_r}\tau_d}\zeta_{T_rL_r}\overline{L_r} + \frac{\lambda_{T_rI_\beta}\overline{T_r}}{2} - d_{T_r}\overline{T_r} = 0.$$

We also assume that at steady state, 5% of positive $T_r$ growth is due to TGF-$\beta$ induced proliferation and 95% is due to $L_r$ migration to the TS. Hence, we have that,

$$\frac{\frac{V_{\text{LN}}}{V_{\text{TS}}}e^{-d_{L_r}\tau_d}\zeta_{T_rL_r}\overline{L_r}}{0.95} = \frac{\lambda_{T_rI_\beta}\overline{T_r}/2}{0.05}.$$

Solving these simultaneously gives us

$$\begin{aligned}
\mathcal{A}_T &= 3.572 \times 10^5 \text{ (cell cm}^{-3})^{-1}\text{day}^{-1},\\
\zeta_{L_rD_L} &= 1.970 \times 10^{-11} \text{ (cell cm}^{-3})^{-1}\text{day}^{-1},\\
\zeta_{T_rL_r} &= 1.098 \times 10^{-2} \text{ day}^{-1},\\
\lambda_{T_rI_\beta} &= 6.30 \times 10^{-3} \text{ day}^{-1}.
\end{aligned}$$

#### A.12 Cancer Cell Parameters

##### A.12.1 Estimates for $C$

As in [1], we assume that CD8+ T cells and NK cells kill cancer cells with similar potency, so we approximate.

$$\frac{\lambda_{CK}}{4} = \frac{\lambda_{CT_8}}{4} \implies \lambda_{CK} = \lambda_{CT_8}.$$

We also assume that the rate that TNF induces tumour necroptosis is larger than that for IFN-$\gamma$, so we approximate

$$\frac{\lambda_{CI_\alpha}}{2 \times 5} = \frac{\lambda_C I_\gamma}{2}.$$

We also assume that CD8+ T cells kill cancer cells with greater potency than Th1 cells, so that

$$\frac{\lambda_{CT_8}/4}{16} = \frac{\lambda_{CT_1}}{2}.$$

Solving these simultaneously leads to

$$\begin{aligned}
\lambda_{CK} &= \lambda_{CT_8}, \\
\lambda_{CT_1} &= \frac{\lambda_{CT_8}}{32}, \\
\lambda_{CI_\gamma} &= \frac{\lambda_{CI_\alpha}}{5}.
\end{aligned}$$

We note that not considering (2.1) at steady state as part of the parameter estimation process results in more accurate trajectories for the time period of interest. However, this may result in reduced accuracy for trajectories extending significantly beyond this timeframe.

##### A.12.2 Estimates for $N$

Considering (2.2) at steady state without chemotherapy, we have that

$$\frac{\lambda_{CI_\alpha} C}{2} + \frac{\lambda_{CI_\gamma} C}{2} - d_N \overline{N} = 0.$$

Thus, we have that

$$\begin{aligned}
d_N &= \frac{1}{\overline{N}} \left( \frac{\lambda_{CI_\alpha} C}{2} + \frac{\lambda_{CI_\gamma} C}{2} \right) \\
&= \frac{3\lambda_{CI_\alpha} \overline{C}}{5\overline{N}}.
\end{aligned}$$

##### A.12.3 Fitting $r_C$, $\lambda_{CT_8}$, $\lambda_{CI_\alpha}$ and $C_0$

We fit $r_C$, $\lambda_{CT_8}$, $\lambda_{CI_\alpha}$ and $C_0$ by choosing the values such that the steady-state value of $C$ and $N$ is reached at 84 days, ensuring that $C$ and $N$ reach steady state at exactly 84 days. Furthermore, we expect monotonicity in the growth of the total cancer population $(C + N)$ as the cancer progresses without treatment, and so we aim to minimise

$$\text{Objective} = \max \left( \frac{|C(84) - \overline{C}|}{\overline{C}}, \frac{|N(84) - \overline{N}|}{\overline{N}}, \frac{|C(84) + N(84) - (\overline{C} + \overline{N})|}{\overline{C} + \overline{N}} \right), \tag{A.14}$$

subject to

$$\max_{t\in[0,84]}(C(t)+N(t)) \le \overline{C}+\overline{N}. \tag{A.15}$$

We perform a parameter sweep to minimise (A.14) subject to (A.15), and set the parameter space to be $r_C \in (0 \text{ day}^{-1}, 2 \text{ day}^{-1}]$, $\lambda_{CT_8} \in (0 \text{ day}^{-1}, 1\times10^{-6} \text{ day}^{-1}]$, and $C_0 \in (8\times10^7 \text{ cell/cm}^3, 10^{11} \text{ cell/cm}^3]$, ensuring that all model parameters are positive. The optimal values of $\lambda_C$, $\lambda_{CT_8}$, and $C_0$ were found to be

$$\begin{aligned}
r_C &= 0.534 \text{ day}^{-1},\\
\lambda_{CT_8} &= 4.00\times10^{-8}\ \left(\text{cell/cm}^3\right)^{-1}\text{day}^{-1},\\
\lambda_{CI_\alpha} &= 0.704 \text{ day}^{-1},\\
C_0 &= 1.75\times10^{8} \text{ cell/cm}^3,
\end{aligned}$$

which implies that

$$\begin{aligned}
\lambda_{CK} &= 4.00\times10^{-8},\ \left(\text{cell/cm}^3\right)^{-1}\text{day}^{-1},\\
\lambda_{CI_\gamma} &= 14.08\times10^{-2} \text{ day}^{-1},\\
\lambda_{CT_1} &= 1.25\times10^{-9}\ \left(\text{cell/cm}^3\right)^{-1}\text{day}^{-1},\\
d_N &= 4.81\times10^{1} \text{ day}^{-1}.
\end{aligned}$$

To estimate the values of $\lambda_{CP}$, $\nu_{CP}$, and $\lambda_{CA}$, we first note that as $P\to\infty$ and $A\to\infty$, we reasonably expect that all cancer growth is stopped, but not reversed, so that $\lambda_{CP}+\nu_{CP}+\lambda_{CA}=1$. We assume that as $P\to\infty$ and $A\to\infty$, 70% of cancer growth inhibition and killing arise due from carboplatin, whilst 30% arises from paclitaxel, so that $\lambda_{CA}=0.7$, whilst $\lambda_{CP}+\nu_{CP}=0.3$. We further assume that paclitaxel induces 20 times more apoptosis of cancer cells than necrosis, so that $\lambda_{CP}=20\nu_{CP}$. Solving these simultaneously leads to $\lambda_{CP}=2.857\times10^{-1}$, $\lambda_{CA}=7.000\times10^{-1}$ and $\nu_{CP}=1.429\times10^{-2}$.

#### A.13 Drug Estimates

To determine $f_X$, where $X\in\{A,P\}$, we use the formula

$$f_X=\frac{C_{\max}(X)}{\xi_X}, \tag{A.16}$$

where $C_{\max}(X)$ corresponds to the maximum serum concentration of $X$ after a dose, $\xi_X$ mg, of $X$ is administered.

We use data from [31], where advanced ovarian cancer and endometrial cancer patients received 5 mg/mL min AUC carboplatin and 175 mg/m$^2$ of paclitaxel, administered via IV infusion every 3 weeks—over 0.5 hours for carboplatin and over 3 hours for paclitaxel. Patients were also given a weekly oral administration of 60 mg selinexor. The $C_{\max}$ of paclitaxel was, on average, 3159 ng/mL whilst the $C_{\max}$ of carboplatin was, on average, 20765 ng/mL. We assume these concentrations to be the same in tissue, and assume a patient surface area of 1.73m$^2$, the reference body surface area for an average 70 kg human. This results in $f_P=1.043\times10^{-8}\ \left(\text{g/cm}^3\right)/\text{mg}$.

To estimate $f_A$, we first use the Calvert formula which determines the appropriate dose of carboplatin (in mg) based on the desired AUC (in mg/mL min) and the patient's glomerular filtration rate

(GFR) (in mL/min/1.73m$^2$), taking into account the patient's renal function [75].

The Calvert formula states that

$$\text{Carboplatin Dose (mg)} = (\text{GFR} + 25) \times \text{AUC}.$$

The mean GFR in OC patients was found to be 73.2 mL/min/1.73m$^2$ [76] and we assume that this is the case here so that

$$\text{Carboplatin Dose (mg)} = 98.2 \times \text{AUC}.$$

This results in $f_A = 4.229 \times 10^{-8}\ (\text{g/cm}^3)/\text{mg}$. We note that it was reported that the $C_{\max}$ was similar to those observed in other clinical trials where the chemotherapy agents were administered as monotherapy.

### B Heatmaps of TVR, Efficacy, Efficiency and Toxicity for Various Chemotherapy Regimens

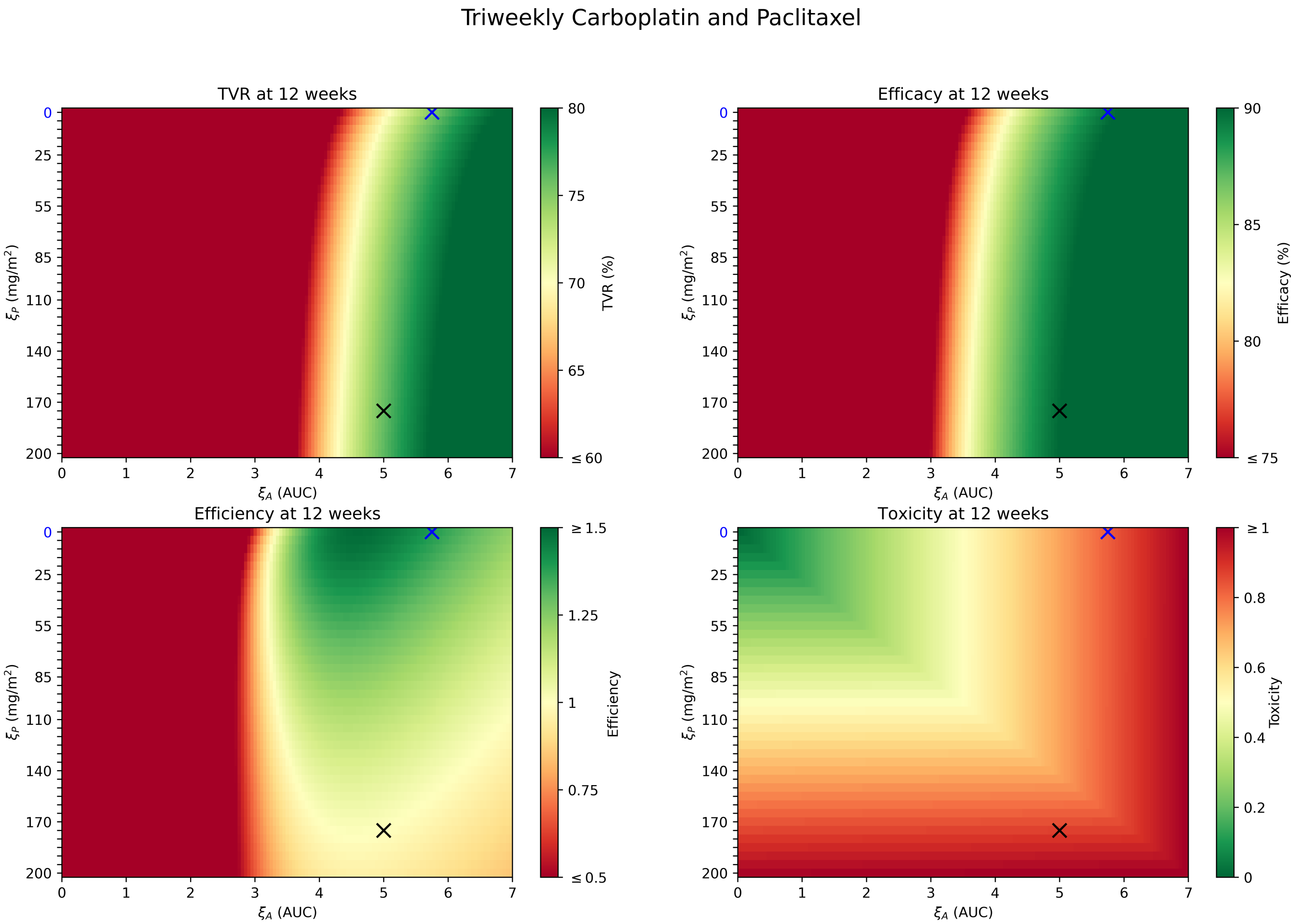


Figure B.1: Heatmaps of TVR, efficacy, efficiency and toxicity for triweekly carboplatin and paclitaxel. The blue cross indicates the optimal regimen, and the grey cross indicates the SR.

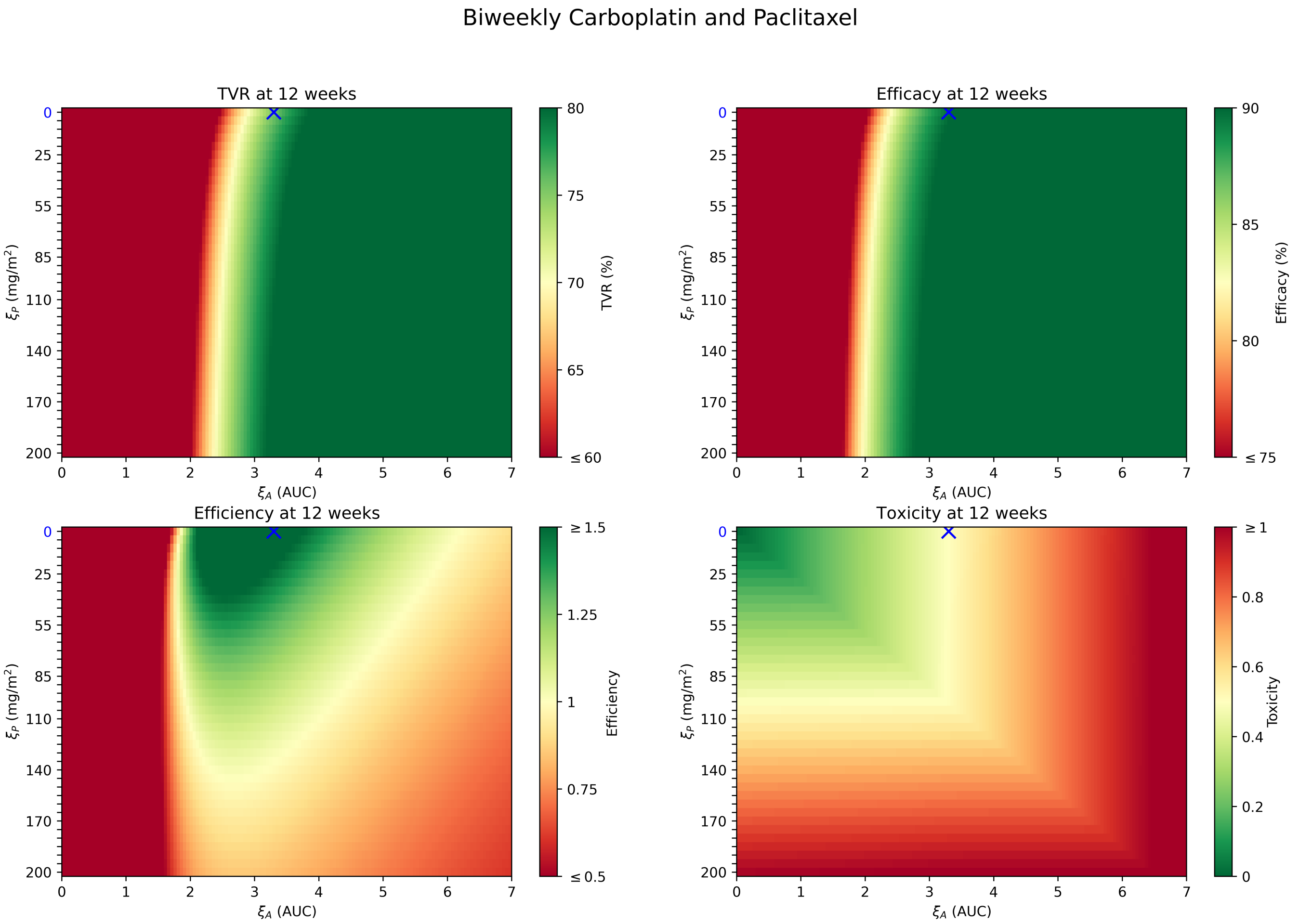


Figure B.2: Heatmaps of TVR, efficacy, efficiency and toxicity for biweekly carboplatin and paclitaxel. The blue cross indicates the optimal regimen.

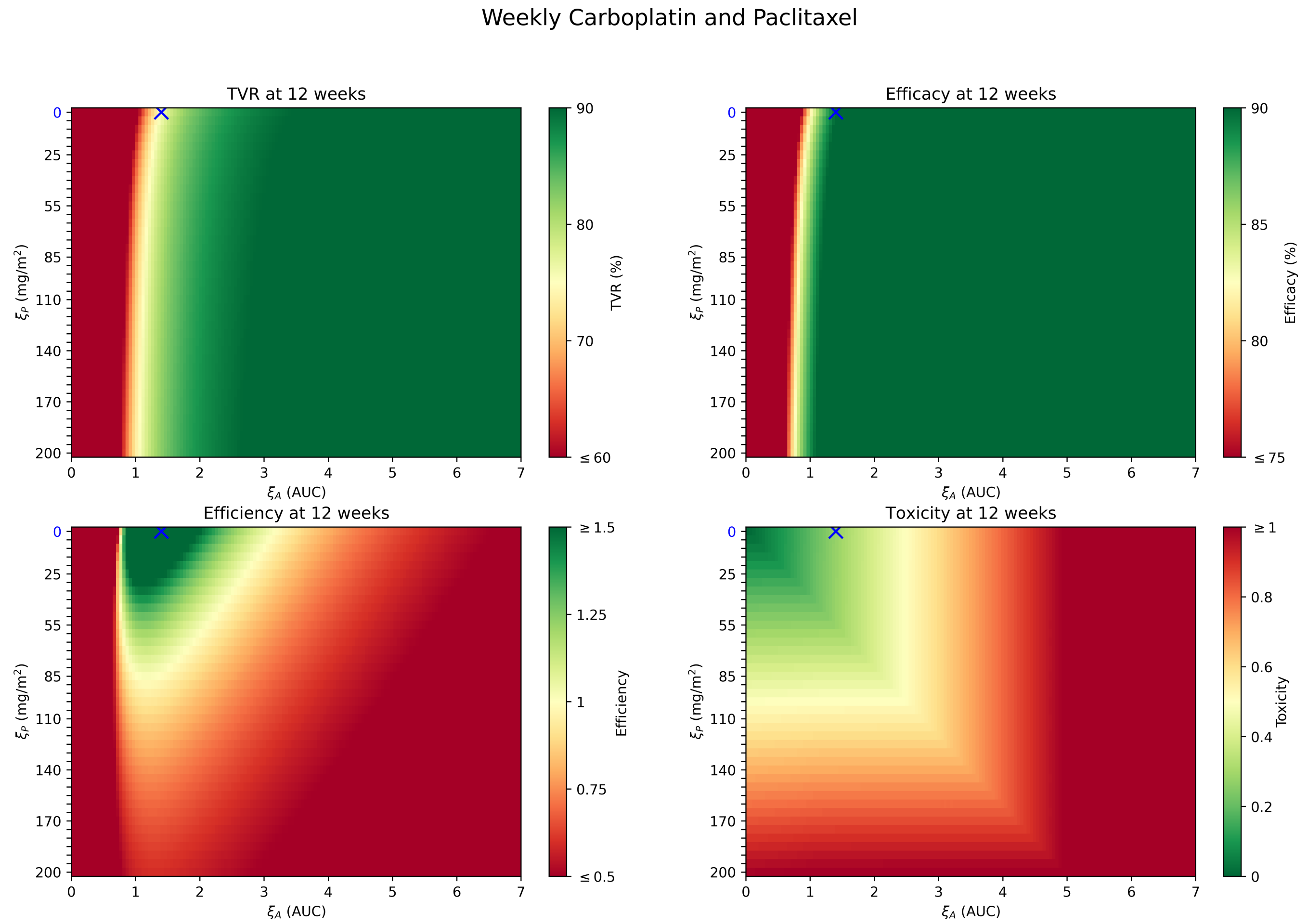


Figure B.3: Heatmaps of TVR, efficacy, efficiency and toxicity for weekly carboplatin and paclitaxel. The blue cross indicates the optimal regimen.

[5] Liebner DA, Huang K, Parvin JD. MMAD: microarray microdissection with analysis of differences is a computational tool for deconvoluting cell type-specific contributions from tissue

samples. Bioinformatics. 2013 Oct;30(5):682–689. Available from: https://doi.org/10.1093/bioinformatics/btt566.

[52] Chapman PB, Lester TJ, Casper ES, Gabrilove JL, Wong GY, Kempin SJ, et al. Clinical pharmacology of recombinant human tumor necrosis factor in patients with advanced cancer. Journal

of Clinical Oncology. 1987 Dec;5(12):1942–1951. Available from: http://dx.doi.org/10.1200/JCO.1987.5.12.1942.

[70] Anaya JM, Shoenfeld Y, Rojas-Villarraga A, Levy RA, Cervera R. Autoimmunity: from bench to bedside [Internet]. El Rosario University Press; 2013.
